# Deep generative embeddings of gene expression and splicing reposition the interpretation of single-cell transcriptomic signatures

**DOI:** 10.64898/2026.09.23.753495

**Authors:** Friedrich-Maximilian Weberling, Ioakeim Ampartzidis, Irina Mohorianu, Florian Hollfelder

## Abstract

Single-cell transcriptomic analysis predominantly derives cell identity from gene expression analysis, while alternative splicing is processed separately despite its fundamental role for cell homeostasis. To overcome the limits of separate investigations, we developed a probabilistic deep learning framework, *Crecerelle*, enabling resolution of the contributions of gene expression and alternative splicing in each cell. *Crecerelle* learns cell embeddings from gene expressions and alternative splicing isoforms, to decipher their mutually dependent impact on the functional characterisation of cells in a data-driven manner, exemplified for the Tabula Muris dataset. This is enabled through a zero-and-N-inflated Dirichlet-Multinomial for a variational autoencoder that learns cell embeddings solely from splicing profiles, as well as a bi-modal variational autoencoder with a relevance-weighted mixture-of-experts variational posterior to consolidate the modality-specific contribution at single-cell level. *Crecerelle* reveals cell-type-specific isoform markers as well as subpopulations with unique isoforms and uncovers regulatory and disease-associated pathways not detected by gene expression analyses alone. This scalable and interpretable framework thus allows a more holistic study of transcriptomic regulation and will open a route to modality-relevance-weighted investigations across single-cell multiomics datasets and their influence on cellular homeostasis, tissue development and disease phenotypes.

## 1 Introduction

Recent advances in single-cell RNA sequencing (scRNA-seq) are enabling the increasingly comprehensive characterisation of RNA processing, including coding and non-coding RNA as well as alternatively spliced transcript isoforms through full-length, long-read, and higher-coverage technologies such as Smart-seq2 [1], Nanopore sequencing [2], Smart-seq3xpress [3], and Vasa-seq [4]. As sequencing costs decline, increasing sequencing depth further expands the potential to resolve transcriptomic and isoform-level variation at single-cell resolution. However, most analytical approaches applied to datasets from these technologies take only overall gene transcript abundance into account for the inference of cellular phenotypes [5, 6]. This prevents the assessment of the regulatory and functional roles of spliced isoforms that are foundational for cell homeostasis. Indeed, post-transcriptional alternative splicing (AS) is present in up to 94 % of multi-exon human genes [7], up to 60 % in mouse [8], and up to 80 % in plants [9]. By generating diverse mRNA isoforms, AS expands the regulatory and functional output of individual genes by enabling cell- and tissue-specific functionalities [10]. Dysregulation of this key driver of evolution and cell homeostasis [11] has been linked to a wide spectrum of pathologies in humans, including Alzheimer’s and Parkinson’s disease, amyotrophic lateral sclerosis and several other neurodegenerative diseases [12], cancer [13], and cardiovascular pathologies [14]. In plants, aberrant AS has been implicated in pathogen-specific transcriptomic changes [15, 16].

AS-focused analyses, mainly carried out on datasets of bulk RNA-seq [17, 18] and scRNA-seq [19–21], are typically analysed separately from the overall gene expression, limiting the ability to jointly capture gene expression and transcript usage at single cell resolution. Thus, current data-driven transcriptomic workflows only assess one of two information inputs, despite evidence that these two transcriptomic layers can provide mutually dependent interpretations of cellular phenotype and regulation [11, 22]. Currently, no large-scale method exists that combines these two layers of analysis to test and understand the synergistic impact of gene expression levels and AS on development and disease.

One approach to integrate total unspliced and spliced transcript abundances to model cell-state transitions, first through ordinary differential equations under a steady-state assumption [23] and subsequently through dynamical [24] and deep generative frameworks [25, 26] is called ‘RNA velocity’. However, RNA velocity remains too coarse-grained for deeper analysis of AS influence on development and disease: while AS can produce multiple isoforms with distinct regulatory and functional implications, RNA velocity only considers the ratio between spliced or unspliced transcripts. Thus, existing RNA velocity methods cannot integrate isoform usage patterns or explicitly model the interplay between gene expression levels and splicing-derived transcript diversity.

To address this gap, our study aims to expand single-cell transcriptomic analysis beyond assessment of gene expression by developing a method that can analyse and interpret the regulatory interplay between gene expression levels and isoform usage. This allows to better resolve cell states and gain insight into how AS programs contribute to cell homeostasis, phenotypic heterogeneity, and disease-associated transcriptional changes [27].

Deep generative modelling [28] has been shown to be a successful approach to learn cell embeddings from single-cell gene expression [6, 29] but remains under-exploited for learning directly from single-cell transcript usage profiles shaped by AS. This is due to the challenges that arise when adopting conventional single-cell gene expression analysis pipelines. Gene expression is typically represented as one abundance value per gene and cell, but transcript usage profiles encode a one-to-many relationship between genes and isoforms, as one gene can give rise to multiple isoforms. The exon junction reads describing the spliced variants may be compositional, gene-constrained, and sparse, due to coordinated exon inclusion, intron excision, alternative transcript structure, or incomplete transcript capture. Current approaches only partially address these challenges, either by augmenting transcript usage profiles in ways that can obscure cell-specific isoform patterns [30], or by integrating AS with gene expression through multimodal variational autoencoders without directly exposing the relative contributions of each transcriptomic layer [31, 32].

To overcome the lack of a scalable and interpretable method for directly modelling sparse, compositional AS-induced single-cell transcript usage profiles and to integrate them with corresponding gene expression data while preserving the contribution of each transcriptomic layer, we introduce the deep generative framework *Crecerelle*^1^ to learn and interpret isolated and joint relevance-weighted cell embeddings from gene expression and AS-induced transcript usage profiles. *Crecerelle* models transcript usage data directly with a zero-and-N-inflated Dirichlet-Multinomial observation model [33] and removes the need for data augmentation or over-reliance on gene expression data by probabilistically accounting for the sparsity and compositionality of single-cell splicing profiles directly. Furthermore, *Crecerelle* integrates gene expression with transcript usage profiles through a relevance-weighted mixture-of-experts variational posterior, designed as bi-modal Gaussian mixture model with learnable parameters, allowing a data-driven interpretation of the importance of each transcriptomic facet in the consolidated representation of a single cell. Applied to heart and brain tissues from the mouse cell atlas *Tabula Muris* [34], *Crecerelle* identifies cell-type-specific information encoded by isoform usage. This approach allowed us to test whether AS provides a complementary, discriminative signal beyond gene expression across tissues, cell types, and sub-cell-type states. Indeed, with *Crecerelle*, we found cell-type-specific isoform markers, isoform-specific sub-cell type populations, recover biological pathways not identified in a gene expression-only approach, and derive disease-relevant markers from splicing-aware representations. Taken together, this deep generative framework monitors, models and distinguishes the regulatory interplay of AS and gene expression on a single-cell level. The interpretations emanating from *Crecerelle* do not only enhance the functional and regulatory characterisation of cell differentiation but may also contribute to novel AS-targeted therapeutic approaches by increasing our understanding of the disease-relevance of AS dysregulation.

## 2 Results

### 2.1 *Crecerelle*: A unified deep generative framework for gene expression and alternative splicing analysis at single cell level resolution

To analyse gene expression and AS as mutually dependent layers of transcriptomic regulation [11, 27, 35], we developed *Crecerelle*, a deep generative framework that combines the published scVI model [29] with two newly developed variational autoencoders, tuVI and TRVI, to learn cell embeddings from gene expression, AS-induced transcript usage or both transcriptomic layers. Existing methods model AS-induced transcript usage (TU) independently [30] or jointly with gene expression (GE) [32], but these models have not been unified with established gene expression modelling in a single workflow that supports both modality-specific and interpretable joint cell representations.

To retain interpretability when integrating gene expression and transcript usage, we first considered how modality-specific cell embeddings are combined in the broader field of single-cell multiomics (Fig. 1a). Multimodal VAEs [36] combine modality-specific cell embeddings modelled as Gaussians using distributional averaging [31], using a product of experts (PoE) [37], or using a fixed mixture of experts (MoE) [38] [38]. A PoE combines modality-specific cell embeddings into a normalised products of Gaussians not providing explicit, normalised estimates of their individual contributions. A fixed-weight MoE retains the modality-specific cell embedding components but assigns them predetermined contributions that are not learnt from the data directly. Learnt cell-specific weighted averages have been explored by MultiVI [31], but they essentially average two cell embeddings into a single Gaussian cell embedding losing the flexibility and interpretability of a multiple component Gaussian mixture model and do not learn the weights from the single-cell profiles directly. We therefore developed for the integration of gene expression and AS-induced transcript usage data a modality-relevance-weighted MoE cell embedding that combines gene expression and transcript usage cell embeddings using cell-specific modality-relevance weights, *π*^(GE)^(**x**^(GE)^) and *π*^(TU)^(**x**^(TU)^), constrained such that *π*^(GE)^(**x**^(GE)^) + *π*^(TU)^(**x**^(TU)^) = 1, and learnt from the single-cell profiles directly. This formulation retains a two-component Gaussian mixture posterior while estimating the relative contribution of each transcriptomic layer for every cell.

**Fig. 1.**
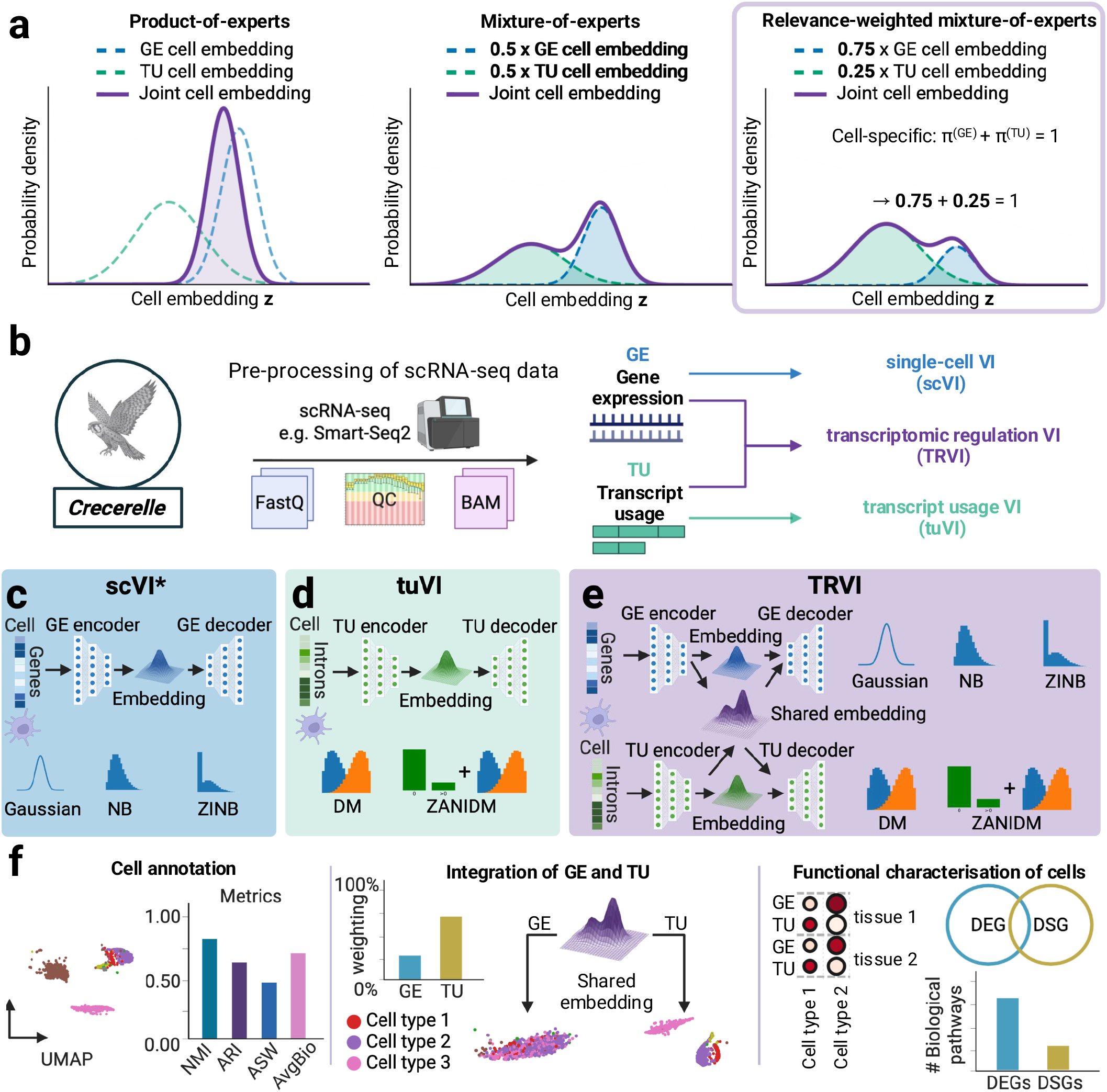
*Crecerelle* is a single-cell-centric suite of deep generative models enabling the analysis of changes in transcriptomic signatures based on gene expression and AS-induced transcript usage. **a** Learning joint cell embeddings from different modalities is done via product-of-experts, or fixed mixture-of-experts approaches that fall short due to obscuring the contribution of each modality to the joint cell embedding. Our modality-relevance-weighted mixture-of-experts approach learns for each cell an embedding where the importance of each modality for the joint representation is learnt. **b** Pre-processing and quality checks of scRNA-seq yield a gene expression (GE-) and transcript usage (TU-) matrix which are then used to learn cell embeddings through the variational autoencoders single-cell VI (scVI), transcript usage VI (tuVI), and transcriptomic regulation VI (TRVI). **c** Cell embeddings based solely on gene expression data are learnt through scVI (* published in [29]). **d** Cell embeddings based solely on AS-induced transcript usage data are learnt through tuVI. **e** Joint modality-relevance-weighted cell embeddings (GE-TU) of gene expression and transcript usage are learnt using TRVI. **f** *Crecerelle* then employs, the GE, TU, or GE-TU cell embeddings for data-driven cell annotation, cell-type clustering, differential gene expression analysis, differential splicing analysis, integrated modelling of gene expression and transcript-usage profiles, and functional cell characterisations.

To learn gene expression, AS-induced transcript usage, and joint modality-relevance-weighted cell embeddings, *Crecerelle* requires matched single-cell gene expression and transcript usage matrices quantifying gene expression counts and exon-junction counts (Fig. 1b). The input matrices for this pipeline consist of curated datasets generated in the following ways: (i) By representing each cell through a set of highly variable genes determined by ClustAssess [39], we constructed a gene expression matrix analogously to those commonly used in scRNA-seq analyses. (ii) To dissect AS, each cell was analysed for its alternatively spliced genes, each partitioned into independent isoform groups (intron groups), to give rise to the transcript usage matrix. Each isoform group consists of a set of exon junctions, whose counts quantify the relative usage of AS outcomes within that group. This pipeline follows transcript usage formulations used in bulk and single-cell splicing analysis [18, 30]. From these paired inputs, *Crecerelle* uses the three variational autoencoders ‘single-cell Variational Inference’ (scVI) [29], ‘transcript usage Variational Inference’ (tuVI), and ‘Transcriptomic Regulation Variational Inference’ (TRVI) to learn modality-specific cell embeddings from gene expression or AS-induced transcript usage alone or modality-relevance-weighted joint embeddings that integrate both layers.

For gene expression cell embeddings, we incorporated the published scVI model [29] (Fig. 1c). For AS-induced transcript usage, we developed tuVI (Methods 4.2), which learns cell embeddings directly from grouped exon-junction counts (Fig. 1d). In existing AS-focused models, the likelihood of observed transcript usage data has been modelled with a Dirichlet-Multinomial (DM) distribution [18, 30, 32]. Although the DM distribution captures the grouped compositional structure and overdispersion of transcript usage counts, it does not explicitly account for the excess zeros and boundary observations prevalent in sparse single-cell exon-junction data (Supplementary Note 2.1). Therefore, based on a finite mixture distribution [33], we developed the zero-and-*N*-inflated Dirichlet-Multinomial distribution (ZANIDM) as the observation model for the transcript usage profiles (Methods 4.2.5, Supplementary Note 2.2) to overcome the data-modelling constraints (Supplementary Note 2). Consequently, tuVI can learn cell embeddings directly from transcript usage without requiring data augmentation or paired gene expression information, as used by existing approaches [30, 32]. Because exact variational autoencoder training under this observation model is only partially parallelisable, we further developed a heuristic surrogate of the ZANIDM log-likelihood (Methods 4.2.5, Supplementary Note 2.3) that we named zero-inflated Dirichlet-Multinomial (ZIDM). The performance and fidelity of the heuristic ZIDM was evaluated with a sample dataset (*Tabula Muris*) [34], resulting in a reduction of the training time (compared to using the full ZANIDM distribution) by 27-fold, from 24 hours to 53 minutes, while preserving the geometry of the learnt latent space in terms of relative distance between the cell embeddings (Supplementary Note 5).

To integrate gene expression and AS-induced transcript usage, we developed TRVI (Fig. 1e, Methods 4.3, Supplementary Note 3). Its modality-specific encoders parameterise gene expression cell embeddings and transcript usage cell embeddings, which are combined through the modality-relevance-weighted MoE formulation described above. We trained TRVI by maximising a Monte Carlo estimate of the evidence lower bound and applied load-balancing and entropy regularisation to the modality weights. The resulting cell embeddings of TRVI provide a joint GE-TU cell embedding together with modality-specific components and per-cell modality-relevance weights that quantify the importance of gene expression or transcript usage for the individual cell.

Taken together, *Crecerelle* forms an end-to-end single-cell gene expression and AS framework built upon the deep generative models of scVI, tuVI, and TRVI to unify data-driven cell annotation, cell-type clustering, differential gene expression analysis, differential splicing analysis, and integrated modelling of gene expression and transcript usage profiles (Fig. 1f). *Crecerelle* is easy to use and seamlessly integrates with existing single-cell libraries such as Scanpy [40] and Seurat [41] while maintaining the interpretability and possibility for human interaction.

### 2.2 Alternative splicing captured through transcript usage profiles faithfully predicts cell types

With *Crecerelle* in hand, we first probed whether gene expression data alone are predictive of cell types. To this end we trained scVI with a zero-inflated Negative Binomial observation model (scVI-ZINB) on the *Tabula Muris* [34] gene expression dataset (Fig. 1c). The learnt gene expression embeddings resulted in well-separated, cell-type-specific UMAP clusters that largely coincided with the reference cell-type annotation (Fig. 2a). This observation demonstrates that variation in expression levels alone capture sufficient biological signal to distinguish major cell types in the dataset.

**Fig. 2.**
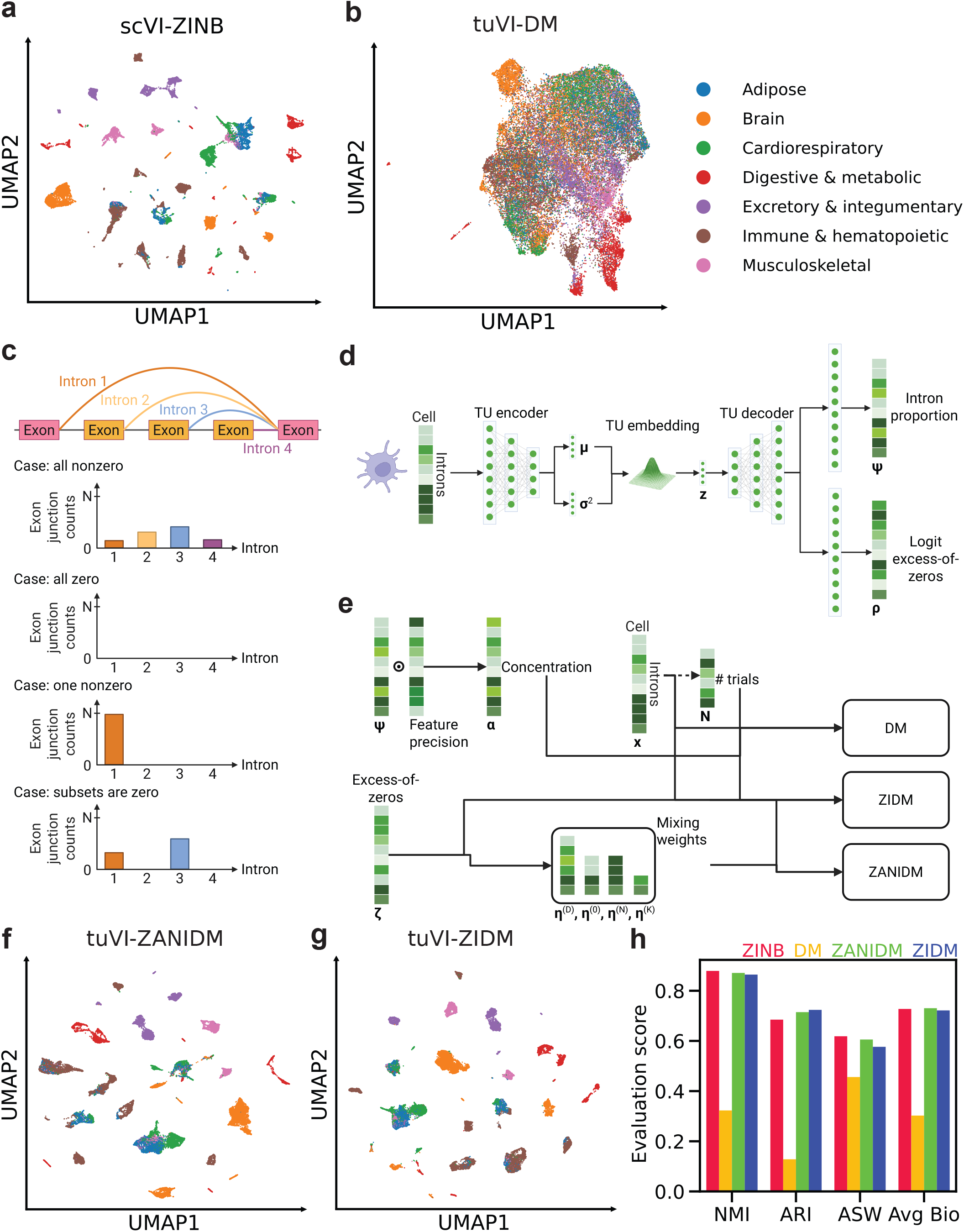
tuVI learns cell embeddings from transcript usage patterns. **a**,**b** UMAPs of *Tabula Muris* (TM) cell embeddings learnt from gene expression using scVI-ZINB (**a**) or transcript usage (TU) using tuVI with a Dirichlet–multinomial observation model (tuVI-DM; **b**). Colours denote organ systems. Compared with the separated scVI-ZINB clusters, tuVI-DM produces a compact embedding with limited biological structure. **c** Schematic of a complex splicing event comprising three alternative exons and four introns quantified by exon-junction counts. TU profiles are classified as all nonzero, all zero, one nonzero or partially nonzero. The standard DM pipeline excludes all-zero profiles, although these represent the most frequent case. **d** tuVI architecture: a TU encoder maps intron counts to a latent embedding, from which the decoder predicts intron proportions ***ψ*** and the excess-of-zeros probabilities logit ***ρ*** = logit(***ζ***). The standard DM uses only ***ψ***, whereas zero-inflated models also represent all-zero profiles. **e** Construction of the observation-model parameters from predicted intron proportions ***ψ***, excess-of-zeros probabilities ***ζ*** and observed junction counts **x.** The DM is parameterised by the concentration vector ***α*** and total count *N*; ZANIDM additionally uses mixture weights *η*^(*D*)^, *η*^(0)^, *η*^(*N*)^ and 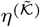; and ZIDM provides an efficient heuristic surrogate of the ZANIDM log-likelihood using the concentration ***α***, the number of exon junction counts *N* of the isoform group and the excess-of-zeros parameter ***ζ*. f**,**g** UMAPs of TM cell embeddings learnt by tuVI-ZANIDM (**f**) and tuVI-ZIDM (**g**). Both models recover cell-type specific clusters, with ZIDM providing a computationally efficient surrogate to ZANIDM. **h** Biological-conservation scores based on normalised mutual information (NMI), adjusted Rand index (ARI), average silhouette width (ASW) and their mean (AvgBio). tuVI-ZANIDM and tuVI-ZIDM outperform tuVI-DM and perform competitively with gene expression cell embeddings learnt by scVI-ZINB.

Assuming that AS is a key driver of cell differentiation, we next asked whether AS-induced transcript usage profiles may also provide similarly informative cell representations. We therefore trained tuVI on *Tabula Muris* [34] transcript usage profiles using a Dirichlet-Multinomial (DM) observation model (Fig. 1d). The resulting transcript usage embeddings formed a UMAP cloud with largely overlapping clusters (except for distinct clusters of brain and digestive cells), making it difficult to infer any functional assignments (Fig. 2b). In contrast, the individual clusters inferred from scVI are well separated. These contrasting outcomes may suggest that AS-derived transcript usage profiles are not sufficiently predictive of cell types.

Indeed, closer examination revealed that the standard DM observation model is mathematically unable to support isoform groups (intron groups) with zero-inflated exon junction counts, a pervasive feature of single cell transcript usage data (Supplementary Note 2.1). This mismatch between the model assumption and ground truth data can explain the inability to discover patterns predictive of cell types. To resolve this mismatch, we considered the possible observation cases for an isoform group (intron group) of a spliced gene (Fig. 2c). For a hypothetical isoform group containing four exon junctions, a cell can exhibit four broad scenarios: all junctions have reads, no junction has reads, exactly one junction has reads, or only a subset of junctions has reads. Across *Tabula Muris*, 92% of observed transcript usage profiles contained no exon-junction reads, 7.2% contained reads for exactly one junction, 0.5% contained reads for all junctions, and 0.3% contained reads for a subset of junctions (Supplementary Note 4 for a benchmarking discussion). As the standard DM likelihood does not explicitly model the dominant all-zero case, it fails to capture the most frequent observation pattern in sparse single-cell transcript usage data, leading to a poor statistical description of transcript usage. However, our newly developed zero-and-*N*-inflated Dirichlet-Multinomial (ZANIDM) observation model allows the modelling of all four possible scenarios. ZANIDM uses - in addition to the intron usage proportion (used by the DM) - the excess-zero logits to capture zero inflation (Fig. 2d). These two parameters define the full ZANIDM likelihood as well as a heuristic surrogate (ZIDM), which simplifies the log-likelihood computation and enables efficient batch optimisation (Supplementary Note 5 for a benchmarking discussion).

We then asked whether tuVI combined with our ZANIDM observation model (tuVI-ZANIDM) would result in better cell annotation than the tuVI-DM model. To resolve this question, we trained our model on *Tabula Muris* transcript usage profiles using either the full ZANIDM observation model (tuVI-ZANIDM) or its ZIDM surrogate (tuVI-ZIDM). Strikingly, both tuVI-ZANIDM and tuVI-ZIDM produced distinct and well-separated transcript usage clusters that corresponded to annotated cell types (Fig. 2f,g) and significantly outperformed tuVI-DM. The UMAP of tuVI-ZIDM embeddings showed a highly similar cluster organisation when compared to tuVI-ZANIDM, suggesting that tuVI-ZIDM is a computationally efficient alternative (Fig. 2g) producing similar results at improved speed. Notably, the transcript usage clusters (Fig. 2f,g) differed from the gene expression clusters in their relative size and organisation, suggesting that AS captures cell-type diversity through a transcriptomic signal that is related to - but not identical - to gene expression.

To quantify the extent to which transcript usage embeddings preserve cell-type identity, we compared our four approaches (scVI-ZINB, tuVI-DM, tuVI-ZANIDM, and tuVI-ZIDM). Performance was assessed using the average biological score (AvgBio), an aggregated metric that summarises multiple measures of biological signal conservation (Fig. 2h, Supplementary Tab. 6). Across all models examined, the tuVI-DM performed substantially worse than the alternative models, achieving an AvgBio score at 0.30 and confirming that the standard DM observation model is insufficient for learning cell-type-predictive transcript usage embeddings from sparse single-cell datasets. In contrast, tuVI-ZANIDM and tuVI-ZIDM models achieved similarly high performance, each reaching AvgBio scores of 0.73 and 0.72 respectively. Notably, these scores are competitive to the one achieved by gene expression embeddings learnt by the scVI model (AvgBio 0.72), demonstrating that transcript usage alone can recover biologically meaningful cell-type structure when equipped with an appropriate observation model.

Together, these findings demonstrate that AS-induced transcript usage profiles can be as informative for cell type identification as gene expression profiles, provided that their zero-dominated and compositional structure is processed by a model that takes into account idiosyncratic data features in this case zero-inflated exon junction counts.

### 2.3 Spliced isoform markers complement marker genes in cell annotation

Data-driven cell annotation strategies based on gene expression [5], rely on clustering and differential analysis of expressed genes to determine cluster-specific marker genes that ultimately reveal cellular phenotypes. With *Crecerelle*, we sought to (i) determine whether the same framework applied to AS-induced transcript usage profiles recapitulates the cellular profiles inferred from gene expression and to (ii) identify divergence across the two modalities. We thus compared cell embeddings, differentially expressed genes (DEGs), differentially spliced genes (DSGs), and cell-type-specific sets of marker genes and isoforms. We focused on non-myeloid brain cells from *Tabula Muris*, since AS has previously been reported to be particularly prominent in brain, heart, and reproductive organs [11]. [An additional case study on heart tissue is provided in Extended Data Fig. A1 and Supplementary Note 8.]

Using the trained scVI-ZINB and tuVI-ZIDM models, we inferred gene expression and transcript usage embeddings for the same individual cells. The UMAP of the gene expression embeddings revealed five distinct clusters corresponding to the five annotated cell types in the dataset: astrocytes, brain pericytes, endothelial cells (ECs), oligodendrocytes (OCs), and oligodendrocyte precursor cells (OPCs) (Fig. 3a). Transcript usage embeddings produced a similarly well-separated organisation of the same major cell types (Fig. 3b). However, a comparison of the corresponding clusters revealed differences in substructure. For example, OC gene expression embeddings (red, Fig. 3b) are compact whereas the corresponding transcript usage embeddings reveal two distinct subpopulations. This suggests that transcript usage may capture cell-state variation related to gene expression, but not identical to it.

**Fig. 3.**
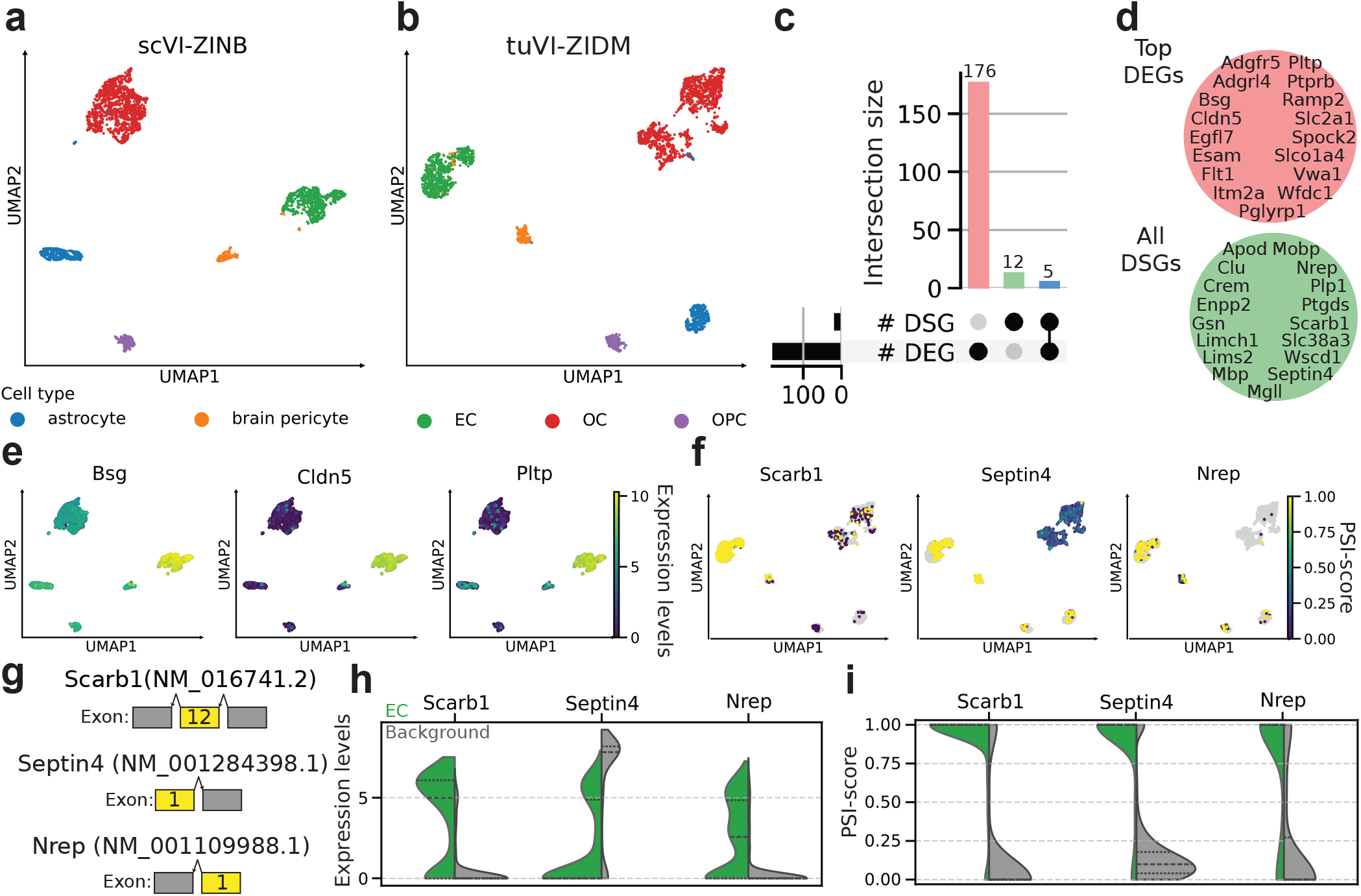
Improvement in cell annotation due to comparative application of gene expression or transcript usage, using *Crecerelle*. **a**,**b** UMAPs of gene expression embeddings learnt by scVI-ZINB (**a**) and transcript usage embeddings learnt by tuVI-ZIDM (**b**) from Tabula Muris (TM) brain-non myeloid cells, coloured by annotated cell type. Both embeddings recover the reference cell types but show modality-specific cluster structure. Intra-cluster differences are visible for oligodendrocytes (OCs) where two sub-populations have emerged for transcript usage embeddings. **c** UpSet plot comparing all significantly differentially spliced genes (DSGs) and differentially expressed genes (DEGs) for endothelial cells (ECs). **d** Venn diagram showcasing that all 17 DSGs and the top 17 DEGs or mutually exclusive i.e. the top DEGs and DSGs are not the same. **e** UMAP-projected gradient of expression for the top three DEGs (*Bsg, Cldn5, Pltp*) showcases their discriminative power as EC cell type markers. **f** UMAP projected gradient of transcript usage for the top three DSG isoforms (*Scarb1, Septin4, Nrep*) shows they are highly discriminative of ECs as well. The colour bar summarises the PSI score. **g** Exon structure of the identified cell-type specific isoforms of *Scarb1, Septin4*, and *Nrep*. **h** Violin plots displaying the gene expression level distribution of the top three DSGs across EC compared to all other cells shows that *Scarb1* and *Nrep* are also differentially expressed for ECs whereas *Septin4* is not. **i** Violin plots visualising the PSI-score distribution of the top three DSG isoforms across EC compared to all other cells indicate that the three spliced isoforms are significantly differentially spliced as well. *Scarb1* and *Nrep* are both DEG and DSG whereas *Septin4* is an example of a DSG that is not a differentially expressed.

We next focused on the EC cluster to evaluate how gene expression and transcript usage representations support cell-type annotation at the marker level. We performed differential gene expression analysis from the gene expression embeddings (giving DEGs) and differential splicing analysis from the transcript usage embeddings (giving DSGs). We identified 176 significant DEGs, 12 significant DSGs, and 5 genes that were both differentially expressed and differentially spliced (Fig. 3c). Comparing the 17 EC-associated DSGs with the 17 top-ranked DEGs reveals that the two sets are mutually exclusive. Also, for the other cell types we found none or a limited overlap (Supplementary Note 7). This observation indicates that EC marker genes and EC-associated spliced isoforms may capture complementary molecular features of cell identity.

Consistent with EC gene expression profiling [42], the top-ranked EC DEGs (*Bsg, Cldn5, Pltp*) showed reduced expression in other brain non-myeloid cell types when projected onto the gene expression UMAP (Fig. 3e). In contrast, the top-ranked EC DSGs (*Scarb1, Septin4, Nrep*) defined a distinct splicing signal. Projection of the percent-spliced-in scores (PSI-scores) of the corresponding isoforms onto the transcript usage UMAP showed that these isoforms were also highly specific to ECs (Fig. 3f). The EC-associated *Scarb1* isoform was majorly reduced in OCs and OPCs, where alternative isoforms were favoured, and was detected only in small subpopulations of brain pericytes. Similarly, the EC-associated *Septin4* isoform was prominent in ECs but reduced in OCs in favour of another isoform. The *Nrep* isoform was enriched and also partially observed in OPCs, brain pericytes, and astrocytes. Together, these isoforms formed a discriminative EC-specific splicing signature. Similarly, cell-type-specific sets of isoforms were identified for the cell type cluster of astrocytes, brain pericytes, OCs, and OPCs (Supplementary Note 7). For the two OC clusters, we found that most isoform markers are shared except a *Lims2* isoform that is unique to the lower left cluster only indicating a potential isoform-specific OC subpopulation (Supplementary Fig. 9).

Finally, to understand if the three spliced isoform markers (Fig. 3g) are only specific with regard to AS or also to expression pattern, we compared the gene expression distributions (Fig. 3h) of the top DSGs with the PSI-score distributions (Fig. 3i) of their EC-associated isoforms. *Scarb1* and *Nrep* showed elevated expression in ECs and a clear increase in PSI-score, consistent with their classification as both DEGs and DSGs. In contrast, *Septin4* did not show EC-specific upregulation at the gene expression level, but its EC-associated isoform showed a significant increase in PSI-score relative to the background. To showcase the generalisability of this cell-type specific isoform marker identification across tissues, we also identified with *Fmo1, Fbln2*, and *Lims2* spliced isoform markers for cardiac fibroblasts (Fbs) in heart tissue (Extended Data Fig. A1, Supplementary Note 8).

Taken together, these observations emphasise that DSGs are not necessarily DEGs, and that differential splicing can reveal cell-type-specific molecular signals that are not evident from gene expression data alone. We conclude that an isoform-centric view of data-driven cell annotation is accessible with tuVI in *Crecerelle*.

### 2.4 Cell-type-specific gene expression and splicing patterns can be resolved through TRVI

We next sought to investigate potential interactions between gene expression and AS, because this layer of regulation can be uniquely captured by simultaneous analysis. To this end, we extended *Crecerelle* to learn joint cell embeddings from matched gene expression and transcript usage profiles. Specifically, we developed a relevance-weighted mixture-of-experts variational autoencoder for joint modelling of gene expression and transcript usage, TRVI (Fig. 4a, Methods 4.3). For each cell, gene expression and transcript usage profiles are encoded by separate deep neural networks into three different embeddings: gene expression embeddings (P-GE), transcript usage embeddings (P-TU), and shared gene expression-transcript usage embeddings (GE-TU). P-GE and P-TU capture variations found only either at the gene expression or at the transcript usage levels. In contrast, the shared GE-TU embedding models all gene expression and transcript usage profiles jointly, weighing them according to their learnt relevance. In this way, *Cre-cerelle* analyses expression levels and spliced isoforms within a shared latent representation, while retaining cell-specific estimates of the relative contribution of each transcriptomic layer.

**Fig. 4.**
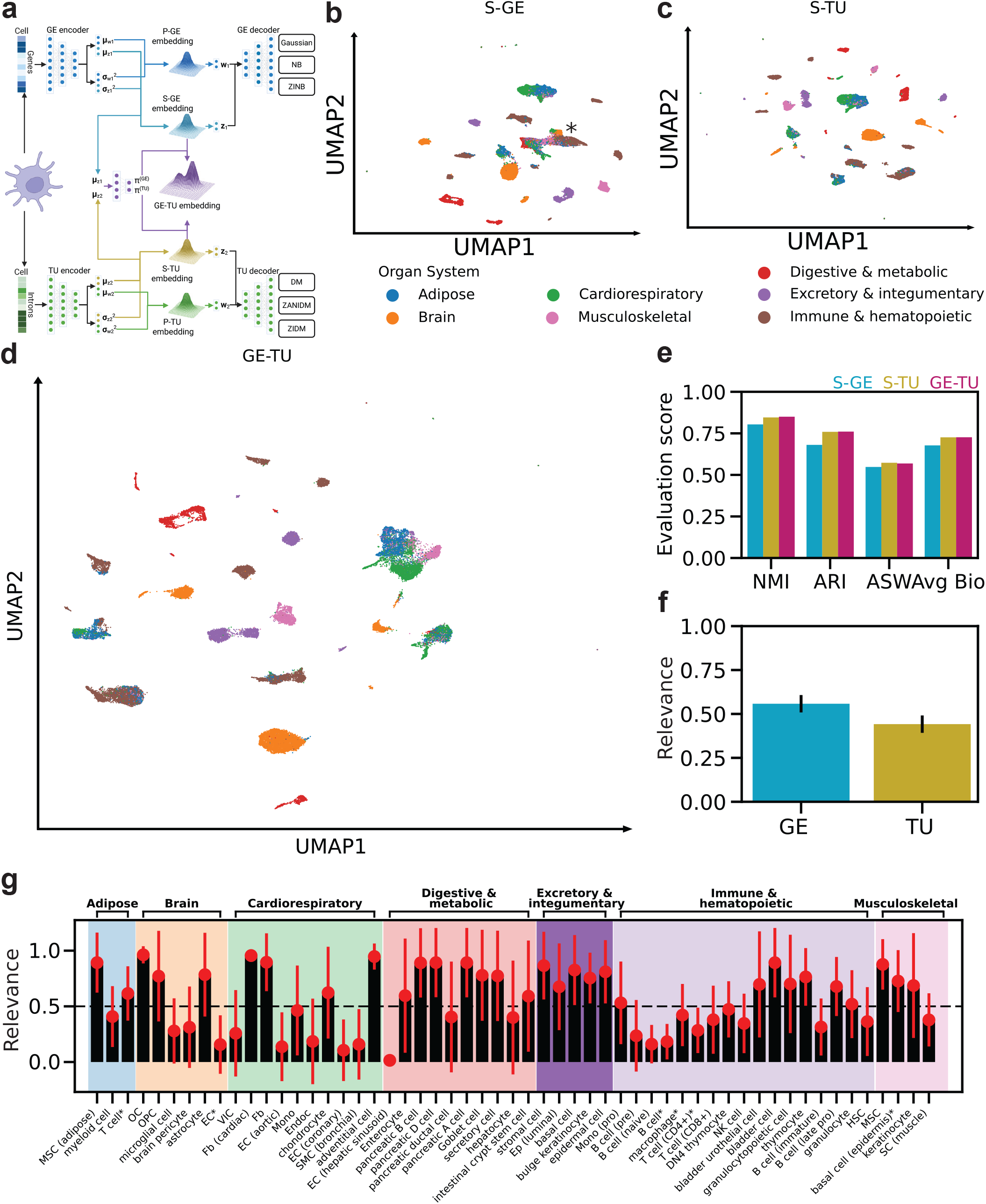
Joint relevance-weighted cell embeddings learnt by TRVI, assess the biological signal in splicing profiles and gene expression with single-cell resolution. **a** TRVI uses a mixture-of-experts variational autoencoder with private gene expression (P-GE) and transcript usage (P-TU) latent spaces and a shared bimodal Gaussian-mixture variational posterior. Its two modes define shared gene expression (S-GE) and transcript usage (S-TU) embeddings, which are combined into joint GE-TU embeddings using learnt GE and TU modality-relevance weights (*π*^(*GE*)^ and *π*^(*TU*)^, respectively). **b–d** UMAPs of S-GE (**b**), S-TU (**c**) and GE-TU (**d**) cell embeddings of *Tabula Muris* (TM) cells, coloured by organ system. All embeddings recover well-separated cell-type clusters; S-GE and S-TU capture complementary structure that is relevance-weighted in GE-TU. **e** Normalised mutual information (NMI), adjusted Rand index (ARI), average silhouette width (ASW) and their mean (Avg Bio) show comparable cell-type predictiveness, with similar performance for GE-TU and S-TU followed closely by S-GE. **f** Across cells, the mean modality-relevance weights are 55.8 % for gene expression and 44.2 % for transcript usage. **g** Cell-type-specific GE modality-relevance weights (*π*^(*GE*)^), averaged across random seeds, reveal modality-dependent importance: for example, transcript usage contributes more to B cells (early–late pro), whereas gene expression contributes more to OCs, pancreatic A, B and D cells, and keratinocytes.

To demonstrate the relevance of such gene expression and AS-induced transcript usage integration, we trained TRVI on *Tabula Muris* and projected the learnt embeddings using UMAPs to compare the cell-type structure captured by each latent representation. We first analysed the modality-specific P-GE and P-TU embeddings which formed densely occupied regions with limited separation by annotated cell type (Supplementary Note 9.2). We then analysed the GE-TU embedding, which we deconvoluted into its two components: shared gene expression (S-GE) (Fig. 4b) and shared transcript usage (S-TU) (Fig. 4c). The S-GE and S-TU UMAPs showed distinctly different cluster organisation but corresponded to the cell types annotated in *Tabula Muris*. In the S-GE embedding, several cell types formed a large, interconnected cell cluster (Fig. 4b; * indicates interconnected cell cluster). The same cell clusters separated more clearly in the S-TU embedding (Fig. 4c). These differences indicate that gene expression and transcript usage encode related but non-identical transcriptomic regulation patterns within the shared latent space.

Next, we assessed the relevance-weighted GE-TU embeddings (Fig. 4d). The GE-TU representation produced well-separated clusters corresponding to annotated cell types but differed from both the S-GE and S-TU embeddings. Notably, the interconnected region observed in the S-GE latent space was resolved in the joint representation. Using the AvgBio score, we benchmarked the recovery of reference cell-type annotations across S-GE, S-TU, and GE-TU embeddings (Fig. 4e, Supplementary Tab. 7). The GE-TU and S-TU embeddings achieved almost identical AvgBio scores of 0.73, whereas the S-GE embeddings performed slightly lower (AvgBio 0.68). These results suggest that AS-induced transcript usage contributes cell-type-discriminative information comparable to gene expression within the shared bimodal GE-TU cell embedding.

The contribution of gene expression and transcript usage to the GE-TU cell embedding was quantified through their learnt modality-relevance weights. We thus aggregated the modality weights across the full cell atlas (Fig. 4f, Supplementary Tab. 8). On average, gene expression contributed slightly more to the joint embedding than transcript usage, with mean weights of 0.558 for gene expression and 0.442 for transcript usage. However, the near-balanced contribution of the two modalities indicates that AS-induced transcript usage provides a transcriptomic regulation pattern that is similarly informative for cellular heterogeneity at atlas scale.

We then wished to understand whether there were cell-type specific variations in the modality weights. To this end, we examined the distribution of modality-relevance weights across cell types (Fig. 4g, Supplementary Tab. 10). We found that OCs, pancreatic A, B and D cells, and keratinocytes were represented predominantly by the gene expression signal, as well as digestive and metabolic cell types. In contrast, endothelial cells (ECs) and their sub types, endocardial cells, and bronchial smooth muscle cells (SMCs (bronchial)) relied more heavily on transcript usage signals. Similarly, B lymphocytes, their developmental subtypes, and many cardiorespiratory cell types were better represented by transcript usage.

Together, assigning single-cell modality-relevance weights to each transcriptomic layer by TRVI reveals where gene expression or AS provides the more informative characteristics of cell-types enabling a comprehensive, dual-perspective analysis of transcriptional abundance and isoform regulation. The ability to quantify their relative contributions at single-cell resolution makes this approach scalable and useful for analysis of transcriptomic regulation, AS programs, and disease patterns.

### 2.5 Unravelling the interplay between gene expression and alternative splicing programs

To evaluate the impact of a consolidated analysis of transcription patterns and isoform regulation on cell characterisation, we again focused on non-myeloid brain cells from *Tabula Muris* and projected the learnt joint GE-TU embedding together with its S-GE and S-TU components using UMAPs (Fig. 5a). While all three representations recapitulated the cell-type annotation of *Tabula Muris*, they unexpectedly differed in their ability to separate closely related populations. Focussing on astrocytes and oligodendrocyte precursor cells (OPCs), two macroglia cell types, we observed differential patterns across the embeddings: S-GE embedding gave an entangled region (Fig. 5a, asterisk/marked region). However, both S-TU embedding and relevance-weighted GE-TU embedding resulted in a clear separation of two clusters (Fig. 5a). We then assessed pairwise distance matrices of the S-GE, S-TU, and GE-TU embeddings and found five distinct cell-type blocks for S-TU and GE-TU but only four for S-GE (Extended Data Fig. A3). To further understand the construction of the GE-TU cell embeddings UMAP we investigated the modality-relevance weights that learnt the relevance of the gene expression and transcript usage profiles with single-cell resolution (Fig. 5b). For astrocytes, OPCs, and brain pericytes the modality-relevance weight for transcript usage was high (0.941, 1.000, 0.899) whereas for OCs and ECs the expression levels were more informative (Supplementary Tab. 11). Visualising the weights on the GE-TU cell embedding UMAP (Fig. 5c), shows that TRVI constructed the GE-TU cell embedding by merging its S-GE and S-TU components based on the learnt modality-relevance weights. Thereby, the joint embedding resolved the separation of OPCs and astrocytes, further supporting the conclusion that TRVI learns cell-type-dependent contributions from both gene expression and transcript usage.

**Fig. 5.**
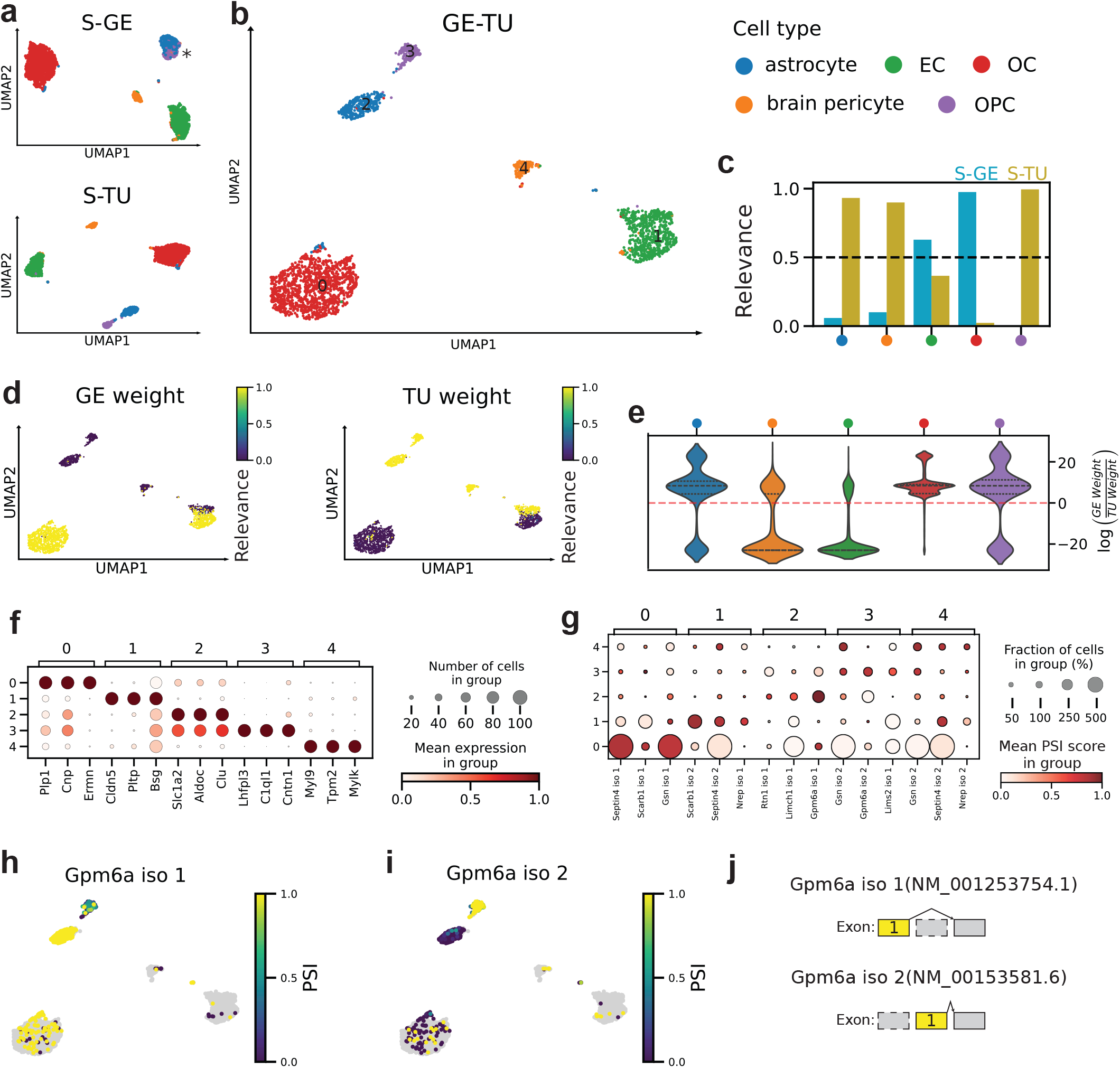
TRVI efficiently deconvolutes the signal from gene expression and transcript usage. **a**,**b**, UMAPs of the shared gene expression (S-GE) and transcript usage (S-TU) cell embeddings (**a**) and joint GE-TU cell embedding (**b**) of *Tabula Muris* (TM) brain non-myeloid cells, coloured by annotated cell type. Astrocytes and oligodendrocyte precursor cells (OPCs), which overlap in S-GE, separate in S-TU and GE-TU. **c** Mean S-GE and S-TU modality-relevance weights by cell type; the dashed line indicates equal weighting. High S-TU weights for astrocytes, brain pericytes and OPCs indicate a greater contribution of transcript usage to their joint representations, whereas oligodendrocytes (OCs) are primarily represented by gene expression. **d** GE-TU UMAPs coloured by cell-level GE and TU modality-relevance weights. The astrocytes and OPCs have high TU weights. **e** To evaluate the robustness of the modality-relevance weights, the violin plots depict the distribution of the log ratio of GE and TU modality-relevance weights per cell type across random seeds revealing GE (*>* 0) or TU (*<* 0) as driving factor for the inferred cell embeddings. For OCs, gene expression yields more cell-type discriminative patterns whereas for brain pericytes and ECs transcript usage is more important. **f**,**g** Dot plots of the top three significantly differentially expressed genes (DEGs; **f**) and differentially spliced gene (DSG) isoforms (**g**) for Leiden clusters 0–4 (0, OCs; 1, endothelial cells; 2, astrocytes; 3, OPCs; 4, brain pericytes). Astrocyte DEGs are also highly expressed in OPCs, whereas DSG isoforms distinguish these populations. *Gpm6a* is differentially spliced in both cell types. **h**,**i** UMAPs coloured by PSI score for two *Gpm6a* isoforms specific to astrocytes (**h**) and OPCs (**i**). **j** Exon structures of the two mutually exclusive, cell-type-specific *Gpm6a* isoforms.

Given that scRNA-seq datasets are inherently noisy [43], potentially affecting deep generative models, we evaluated the robustness of the learnt modality-relevance weights (GE and TU weights) across ten random seeds, corresponding to different initialisation parameters of the model. Some cell types, such as astrocytes, OPCs, and ECs exhibited variable modality contributions when assessed by using different parameter initialisations, whereas others, including brain pericytes and OCs, were highly conserved (Fig. 5c). To determine which transcriptomic layer predominantly contributed to each cell type, we analysed the distribution of log-ratio between gene expression and transcript usage modality-relevance weights across random parameter initialisations (Fig. 5e). OCs showed consistently positive logratios, indicating gene expression-dominated representations, whereas brain pericytes and ECs exhibited predominantly negative log-ratios, indicating transcript usage-dominated representations. Together, these results indicate that the modality-relevance weight distributions learnt from an ensemble of TRVI trained with different parameter initialisations provide a robust data-driven measure of the relative contribution of gene expression and transcript usage to cell-type representation within single-cell atlases.

We then asked whether the joint GE-TU embedding improved downstream cell characterisation in non-myeloid brain tissue. For the five Leiden clusters (0: OCs, 1: ECs, 2: astrocytes, 3: OPCs, 4: brain pericytes) we identified significant DEGs and DSGs (Supplementary Note Fig. 23). The mean expression of the top three DEGs per Leiden cluster revealed cell-type-specific marker-gene programs consistent with known annotations (Fig. 5f). However, the top three astrocyte marker genes (*Slc1a2, Aldoc*, and *Clu*) also exhibit high expression in OPCs, highlighting the close transcriptional similarity between these two macroglia populations (Supplementary Fig. 23).

In contrast, the top-ranked DSG isoforms provided a greater discrimination between cell types (Fig. 5g). Astrocytes and OPCs showed distinct sets of DSG isoforms (*Rtn1* iso1, *Limch1* iso1, *Gpm6a* iso1 for astrocytes and *Gsn* iso2, *Gpm6a* iso2, *Lims2* iso1 for OPCs), consistent with the high transcript usage weights assigned to these populations. We then projected their PSI-score onto the UMAP embeddings which further supported our hypothesis that transcript usage contributed to the separation of these closely related cell types (Extended Data Fig. A4c,d). OCs, ECs, and brain pericytes also exhibited distinct DSGs (Extended Data Fig. A4a,b,e), showing that each cell type could be distinguished by a compact and highly specific set of isoform markers.

At the isoform level within the same gene, we also observed cell-type-specific isoform regulation that would be missed by considering gene expression only. *Gpm6a* is a DSG in both astrocytes and OPCs, but the two cell types exhibited different mutually exclusive isoforms (Fig. 5h–j). We observed a similar regulation in heart tissue for cardiac Fb and coronary EC-specific isoforms of Fbln2 (Extended Data Fig. A2h–j). Together, these findings demonstrate that for scenarios in which the same gene contributes to the identity of multiple cell types, inclusion of different AS programs can facilitate distinction.

### 2.6 *Crecerelle* maps transcript usage-informative cell states and DSG-associated biological pathways

To investigate the atlas-wide contribution of transcriptional abundance and isoform regulation on cellular heterogeneity, across tissue heterogeneity, we analysed the relevance-weighted integration of gene expression and AS-induced transcript usage learnt by *Crecerelle*. Using TRVI trained on *Tabula Muris*, we examined the learnt gene expression and transcript usage modality-relevance weights across individual cells, cell clusters, and tissues. To visualise the relative contribution of each transcriptomic layer across cell types and tissues, we projected tissues as well as the gene expression and transcript usage modality-relevance weights on the UMAP joint GE-TU cell embeddings (Fig. 6a–c). As both weights are complementary, these projections identify cell populations whose joint representation is dominated either by transcriptional abundance or by isoform usage. In many cases, entire clusters showed consistently higher gene expression or transcript usage weights. Unexpectedly, however, substantial intra-cluster variation was also observed, indicating that the relative contributions of gene expression and transcript usage can vary within a single tissue type.

**Fig. 6.**
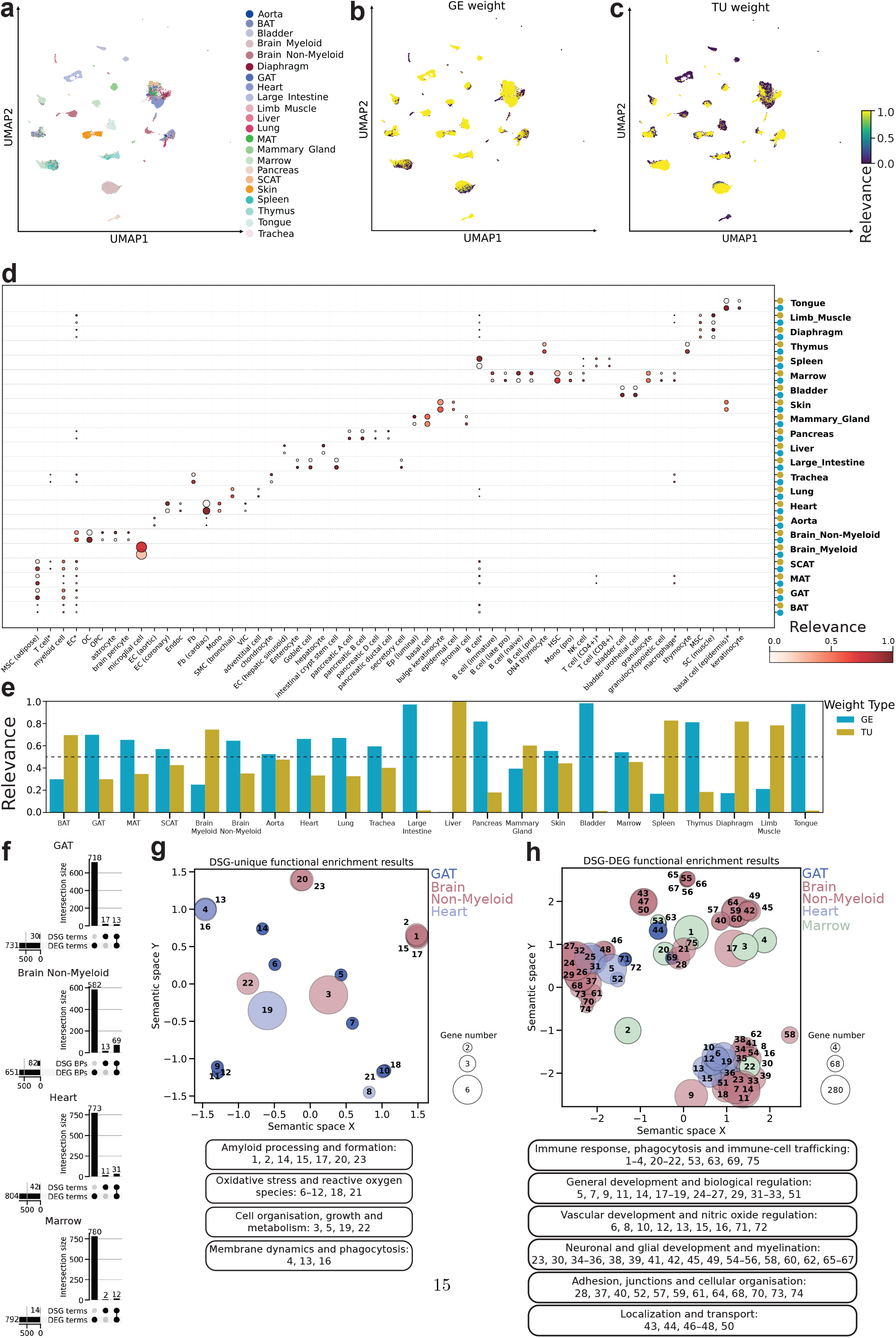
TRVI expands the functional characterisation of cells across tissues and emphasises DSG and DEG specific biological pathways. **a** Joint GE–TU cell embeddings of the Tabula Muris (TM) atlas, coloured by tissue. **b**,**c** The same UMAP coloured by per-cell (**b**) gene expression (GE) and (**c**) transcript usage (TU) modality-relevance weights. **d** Dot plot of mean GE and TU modality-relevance weights across TM cell types and tissues. Each cell type for a tissue is represented through two dots (for gene expression and transcript usage). The learnt modality-relevance weights reveal cell-type and sub-cell type dependent patterns of the learnt relevance of gene expression and transcript usage profiles across the different tissues of the cell atlas. **e** Tissue-level mean GE (blue) and TU (green) weights demonstrate that modality relevance varies not only across cell-types but also tissues. **f** UpSet plots comparing significantly enriched functional terms associated with DEGs and DSGs in GAT, brain non-myeloid, heart and marrow. DSG-based analysis identifies processes missed by DEG-only analysis. **g** Semantic-similarity dot plot of significantly enriched biological pathways (*p*_adj_ *<* 0.05) identified only by DSGs in GAT, brain non-myeloid and heart. **h** Corresponding dot plot of pathways identified by both DSGs and DEGs in GAT, brain non-myeloid, heart and marrow. In (**g**,**h**), proximity indicates semantic similarity, bubble size represents the number of associated genes, colour denotes tissue and numbers refer to the pathway groups listed below. DSG-unique pathways include amyloid processing, oxidative stress and phagocytosis, whereas shared pathways include neuronal development, immune responses and phagocytosis.

To gain further insight into these patterns, we analysed the mean gene expression and transcript usage modality-relevance weights per cell type and sub-cell type across the different tissues (Fig. 6d). Throughout the B-cell lineage, from late pro-B to naïve B-cell stages, the transcript usage weights were consistently high, while late pro-B cells showed a higher contribution of gene expression. Endothelial cells generally exhibited high transcript usage weights across several tissue contexts, including aortic, coronary, and hepatic sinusoid endothelial populations. However, for ECs in brain non-myeloid we observed a tissue-specific modality preference for gene expression.

We then compared modality-relevance weights at the tissue level, by averaging gene expression and transcript usage weights across all cells within each tissue (Fig. 6e, Supplementary Tab. 9). Previously associated tissues with prominent AS activity, including brain and heart [11], showed balanced or increased transcript usage contributions. In bone marrow, transcript usage was also similarly relevant as gene expression, suggesting that AS-induced isoform usage may contribute substantially to the organisation of immune and hematopoietic cell states.

We next asked whether incorporating AS-induced transcript usage expands the functional characterisation of cellular heterogeneity. To achieve this, we performed functional enrichment analysis, using the DEGs and DSGs previously identified in brain non-myeloid, heart, bone marrow, and gonadal adipose tissue (GAT) (Fig. 5, Extended Data Fig. A2, Supplementary Note 10) and compared the significantly (*p*_adj_ *<* 0.05) enriched functional terms recovered from DEGs and DSGs (Fig. 6f). Overall, DEGs yielded substantially more enriched functional terms than DSGs across all tissues, as expected given the larger number of DEGs included in the enrichment analysis. However, each tissue exhibited a set of terms that were only discovered by DSGs and a focussed sets of terms associated with both DEGs and DSGs.

To gain insights into regulatory mechanisms associated with each set, we filtered the DSG-unique and DEG-DSG enrichment terms only to biological pathways. For the DSG-only biological pathways (Fig. 6g), we identified crucial regulatory pathways involved in amyloid processing and formation, oxidative stress, cell growth and metabolism, and membrane dynamics and phagocytosis. Furthermore, the DSG and DEG associated pathways (Fig. 6h) consisted of groups of crucial regulatory processes such as immune response, neuronal development and myelination. Furthermore, beyond these pathways, human phenotypes associated with short-term memory impairment and brain dysfunction have also been identified by these sets of DEGs and DSGs This observation showcases that *Crecerelle* expands the hypothesis space for the interpretation of disease-relevant contexts to include both gene expression and alternative splicing (Supplementary Data Tab. 10). Combined, these findings suggest that AS provides complementary functional insights beyond gene expression analysis, particularly in brain and heart tissue.

Overall, these analyses revealed that TRVI can provide an atlas-wide map of the relative gene expression and transcript usage contributions able to accurately capture the cellular heterogeneity. The combination of modality-relevance-weighted gene expression and transcript usage integration with DEG and DSG functional enrichment analysis, allows *Crecerelle* to identify tissue-wide AS programs and biological pathways.

## 3 Discussion

In this study, we developed *Crecerelle*, a deep generative framework that takes advantage of a previously published model scVI [29] combined with two models we generated herein, tuVI, and TRVI. AS-induced isoform information is analysed independently by tuVI which learns cell embeddings directly from single-cell transcript usage profiles, while TRVI quantifies the relative contribution of each transcriptomic layer at single-cell resolution. Together, Crecerelle extends single-cell transcriptomic analysis beyond transcript abundance and provides a quantitative framework for assessing how isoform regulation contributes to cell identity, tissue organisation, and disease-relevant cellular functions.

A central obstacle for AS-aware single-cell analysis is the sparse and compositional nature of AS-induced transcript usage profiles. Previous approaches relied on paired gene expression information or data augmentation [30, 32], which can limit the recovery of subtle, cell-specific isoform usage patterns such as isoform-specific sub-populations in OCs (Fig. 3b) and cell-type specific isoform markers of cell types such as ECs (*Scarb1, Septin4, Nrep*) (Fig. 3g) or cardiac Fbs (*Fmo1, Fbln2, Lims2*) (Extended Data Fig. A1g). Here, we found that replacing the standard Dirichlet-Multinomial observation model [18] with the ZANIDM observation model (combined with a computationally efficient heuristic surrogate) enabled *Crecerelle* to learn biologically meaningful cell embeddings solely from AS-conditioned transcript usage. Compared to previous studies that made use of only 8% of transcript usage data, the ZANIDM observation model exploits the data more rigorously by also interpreting the missing 92%, suggesting that any conclusions derived from it are much better informed (Supplementary Fig. 3, Supplementary Note 4). These results establish splicing-derived transcript usage as a scalable and informative modality for data-driven cell annotation (Supplementary Fig. 3, Supplementary Note 4), with an Avg Bio score performance of 0.730 outperforming scVI-derived gene expression cell embeddings (Avg Bio 0.727) while maintaining robustness (Fig. 2h, Supplementary Tab. 6).

TRVI enables the interpretability of jointly modelled single-cell modalities through cell-specific ‘modality-relevance weights’. Multimodal product-of-experts variational autoencoders commonly used for single-cell multi-omics integration [31, 32, 44], including MultiVI and SpliceVI, combine modality-specific cell embeddings into a joint unimodal Gaussian variational posterior, whereas existing mixture-of-experts approaches [38] only use fixed modality-specific weights impeding the interpretation of each modality’s contribution to the multi-omic cell representation. By contrast, our joint modelling with TRVI uses a Gaussian-mixture variational posterior with learnable single-cell modality-relevance weights, supported by Monte Carlo sampling and targeted regularisation. This is a unique feature enabling *Crecerelle* to provide a quantitative estimate of the relative contribution of gene expression and transcript usage to each joint cell representation. Across the *Tabula Muris* dataset, gene expression and transcript usage made near-balanced contributions to the joint embedding, but their relative importance varied across cell types (Fig. 4g). For example, B-cell type populations and several cardiorespiratory cell types were strongly represented by transcript usage, while the gene expression signal dominated in other cell types, including pancreatic A, B, C cells, fibroblasts, and MSCs. The patterns observed here support the standing hypothesis that AS is a fundamental, yet heterogeneous, driver of B lymphocyte differentiation and activation [45–47]. Since AS depends on lineage, tissue context, and cellular state [11], *Crecerelle* provides a quantitative framework to monitor and distinguish AS programs alongside gene expression. This dual analysis at the single cell level in heterogeneous samples provides unmatched biological and functional transparency, enabling the identification of complex regulatory scenarios arising from co-varying but distinct AS and gene expression programs that are hard to disentangle using expression-based analyses alone.

This transcript usage information can also be leveraged for cell-type annotation, providing isoform-level markers that complement conventional marker genes. We identified *Bsg, Cldn5*, and *Pltp* as well-known EC marker genes [42]. In parallel, we identified EC-associated isoform markers from the differentially spliced genes *Scarb1, Septin4*, and *Nrep*. These are not established EC markers [42] in the literature but are linked to broader functional and disease-associated contexts [48, 49]. Importantly, some of these signatures were not evident at the transcript level alone. Our findings suggest that isoform-centric annotation complements marker gene analysis by highlighting cell-type-associated splicing events with potential functional and disease relevance.

Another biological scenario that can now be discerned concerns the interplay of differentially spliced genes that may not be differentially expressed. Previous bulk RNA-seq studies of tissues reported limited overlap between tissue-specific sets of differentially expressed genes (DEGs) and differentially spliced genes (DSGs) [22, 35, 50], but could not resolve cell-type-specific alternative-splicing (AS) programs. With Crecerelle, we confirm and extend this observation at single-cell resolution: we observed that cell-type-specific marker genes and isoforms are not the same, as shown for cells in brain non-myeloid (Fig. 3e-f) and in heart tissue (Extended Data Fig. A1e-f). Instead, we observed cell-type-specific regulation at the isoform level within the same gene. For example, *Gpm6a* and *Fbln2* exhibited expression across multiple cell types, but distinct isoforms of these genes were found in different cell populations (Fig. 5h-j, Extended Data Fig. A2h-j). Thus, cell identity that is not solely determined by gene expression levels but also by the regulation of alternative isoforms can be uncovered through *Crecerelle*. This underlines that dual analysis of AS and gene expression unravels layers of cell-type-specific molecular control.

Going beyond single cell identity, we combined relevance-weighted cell embeddings with differential gene expression and differential splicing analyses at tissue level to identify underrepresented biological pathways. Notably, some of these pathways could be recovered solely from DSGs as observed in brain non-myeloid cells, where DSG-associated enrichments highlighted processes related to amyloid-beta regulation, short-term memory impairment, and brain dysfunction. Similar enrichment patterns in heart, bone marrow, and gonadal adipose tissue suggest that incorporating AS into tissue-level analyses can reveal tissue-specific regulatory pathways that may be overlooked be gene expression only workflows. Given the established involvement of AS dysregulation in neurodegenerative disease and cancer [12, 13], *Crecerelle* provides a framework for identifying disease-associated isoform programs across existing and future single-cell datasets, potentially revealing regulatory mechanisms that are not captured by expression-based analyses alone.

These observations suggest that extending our multimodal analysis beyond gene expression and AS-indued transcript usage could provide further resolution for functional cell assignment. Additional modalities (e.g. chromatin accessibility (ATAC), surface-protein abundance, transcription-factor binding, epigenetic modifications) have been modelled before [31, 37, 38, 44, 51] and could thus technically be incorporated into *Crecerelle*’s TRVI by extending its modality-relevance-weighted mixture-of-experts variational posterior from a bi-modal to a multimodal formulation (expand |*ℳ*| = 2 to 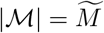 with *ℳ* being the set number of single-cell modalities; compare derivation of multimodal TRVI in Supplementary Note 3). The resulting modality-relevance weights established here will provide a quantitative measure of the contribution of each modality, potentially informing cell-type assignment and functional cluster interpretation and thereby illuminating complex biological scenarios. Importantly, this extension requires only modest additional computational effort, rendering our model scalable and capable to manage multiomic analysis of single cell discoveries. Our model approach would thus support the investigation of complex interdependencies, including how genomic context and genome organisation influence splicing and transcriptomic output [52], by jointly relating AS programs to gene-level abundance and other molecular modalities. Other single-cell multiomics deep learning approaches such as MultiVI ([31], Cobolt ([37], or scMM [38] do not accommodate such correlations.

Finally, *Crecerelle* is an open-source (https://github.com/Hollfelder-Lab/crecerelle) and easy-to-use framework compatible with widely used single-cell analysis ecosystems, such as Scanpy [6, 40] and Seurat [41]. Thus, *Crecerelle* facilitates seamless integration into existing analysis workflows, making it an immediately useful tool. Taken together, these features position *Crecerelle* as a scalable, interpretable, and readily deployable probabilistic deep learning framework for multimodal single-cell analysis, with the potential to enhance the resolution at which cellular identity and regulatory programs can be interrogated and to rapidly attach biological meaning to high throughput single-cell experiments.

## 4 Methods

### 4.1 Data processing, filtering, and quality control

We analysed the Smart-seq2 component of the *Tabula Muris* single-cell atlas [34], processed using the bioinformatics pipeline described by [30]. The initial dataset comprised a gene expression count matrix of 44,518 cells by 27,348 genes and a transcript usage count matrix of the same 44,518 cells by 29,924 exon junctions. The exon junctions were assigned to 13,084 intron groups representing splicing patterns in 7,637 genes.

Gene expression data were processed using Scanpy [40]. We excluded cells with at least 2,000,000 total gene counts, at least 10% mitochondrial counts, at least 15% ribosomal counts or fewer than 200 detected genes. Genes detected in fewer than three cells were also removed. To ensure sufficient representation for downstream analyses, we excluded cell types represented by fewer than 100 cells across the complete atlas and, within each tissue, cell types represented by fewer than 50 cells. Highly variable genes (HVGs) were selected using ClustAssess [39]. After evaluating different feature-set sizes for *Tabula Muris*, we selected 3,000 HVGs based on the stability and noise levels of the resulting low-dimensional representations and clusterings (Supplementary Fig. 1). These filtering and feature-selection steps yielded the gene expression count matrix 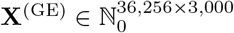. Raw gene counts were retained in a separate AnnData layer, and a normalised, (log(1 + *x*))-transformed representation was generated for downstream visualisation and analysis.

To maintain one-to-one correspondence between modalities, cells removed during gene expression quality control were also removed from the transcript usage matrix. Transcript usage features were subsequently restricted to intron groups associated with the selected HVGs. This resulted in a transcript usage count matrix 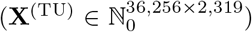, comprising 2,319 exon junctions from 600 alternatively spliced HVGs. Raw exon-junction counts were retained in a separate AnnData layer, and a normalised, (log(1 + *x*))-transformed representation was generated for downstream visualisation and analysis.

This dataset **X** = {**X**^(*GE*)^, **X**^(*TU*)^} of single-cell gene expression (*m* = *GE*) and transcript usage profiles (*m* = *TU*) was split randomly into training, validation, and test sets comprising 29,028, 3,600, and 3,628 cells, respectively.

### 4.2 transcript usage Variational Inference

Transcript usage Variational Inference (tuVI) is a variational autoencoder [28] that learns cell embeddings from single-cell AS-induced transcript usage data using deep neural networks. Given the transcript usage count matrix **X**^(TU)^, tuVI learns a generative model 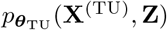 and an amortised variational posterior 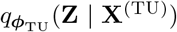, which maps the observed transcript usage profiles to a low-dimensional latent space of cell embeddings **Z** ∈ ℝ^*N ×L*^.

#### 4.2.1 Representation of transcript usage observations

In contrast to gene expression data, transcript usage data are compositional, because one gene *g* can encode several isoforms through alternative splicing (Supplementary Fig. 2). Following existing literature for unsupervised modelling of splicing [18], [30], the transcript usage profile 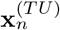 of cell *n* is comprised of *G* independent spliced gene vectors. Each spliced gene vector factorises into independent *I*_*g*_ isoform groups (also called intron groups). For a gene *g* with *I*_*g*_ isoform groups, the profile of isoform group *i* is quantified by the isoform group count vector 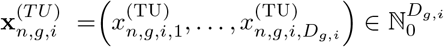 where *D*_*g,i*_ are the number of exon junctions. Each exon junction is quantified by read counts indicating the presence of a spliced isoform. Assuming conditional independence of the isoform group vectors 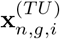, the data likelihood of of a single-cell transcript usage profile 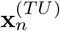 is obtained by

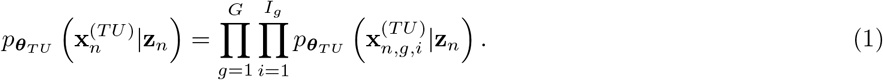

where 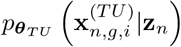 is the transcript usage observation model assessing the plausibility of 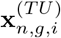 given the latent cell embedding **z**_*n*_ and the parameters ***θ***_*TU*_.

#### 4.2.2 Generative model

The generative model 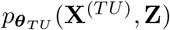 of tuVI factorises across *N* cells. For a cell *n*, it is described by the data likelihood 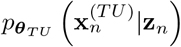 and a standard multivariate Gaussian prior *p*(**z**_*n*_) = *N* (**0, I**_*L*_) on the latent cell embedding **z**_*n*_ ∈ ℝ^*L*^ yielding

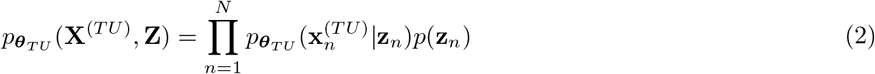

Hereby ***θ***_*TU*_ denote the parameters of the decoder that maps the cell embedding **z**_*n*_ to the parameters of the transcript usage observation model.

#### 4.2.3 Variational posterior

Since inference is generally intractable for a variational autoencoder, tuVI approximates the posterior over the cell embeddings using an amortised variational posterior

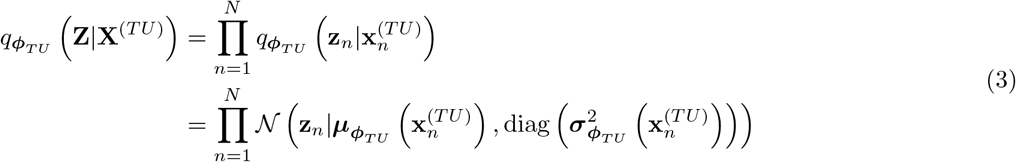

that factorises across cells. The encoder, parameterised by ***ϕ***_TU_, maps the observed transcript usage profile 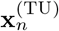 to the mean 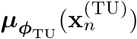 and diagonal variance 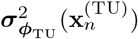 of the variational posterior.

#### 4.2.4 Training objective

To determine the optimal parameters ***θ***_*TU*_, ***ϕ***_*TU*_ of the deep neural networks of the encoder and decoder of tuVI, the evidence lower bound (ELBO) is optimised [28]. For a single-cell transcript usage profile 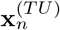 it is given by

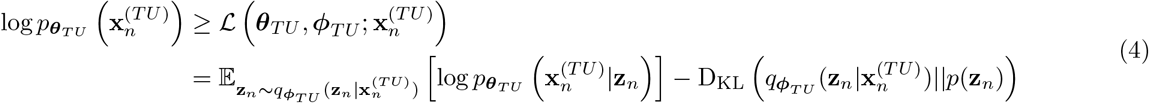

where the expected log-likelihood evaluates the data generation fidelity and the Kullback-Leibler regularises the variational posterior towards the standard Gaussian prior.

#### 4.2.5 Transcript usage observation models

To model an isoform group count vector 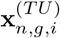, the decoder of tuVI uses a deep neural network 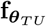 to compute the intron proportions 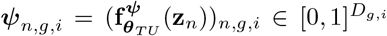, and additionally the excess-of-zeros parameter 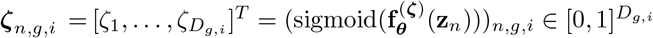, or a logit of the excess-of-zero parameters 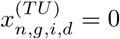 dependent on the observation model chosen. The intron proportion *ψ*_*n,g,i,d*_ quantifies the expected relative usage of exon junction *d* in the isoform group. The excess-of-zero parameter *ζ*_*n,g,i,d*_ quantifies the probability of a zero exon junction 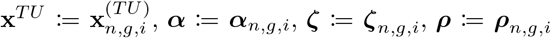.

The intron proportion vector ***ψ***_*n,g,i*_ is then used to define the concentration ***α***_*n,g,i*_ = ***γ***_*n,g,i*_ ⊙ ***ψ***_*n,g,i*_ where ***γ***_*n,g,i*_ is an additional feature precision parameter which we fixed to ***γ*** = **1** in all models. With that, tuVI enables then the application of three different transcript usage observation models: the Dirichlet-Multinomial (DM), the Zero-and-N-Inflated Dirichlet Multinomial (ZANIDM), and a heuristic surrogate called Zero-Inflated Dirichlet-Multinomial (ZIDM). All are designed w.r.t. a cell *n*, a spliced gene *g*, and its isform group *i*. Therefore, to simplify notation throughout this section, we adopt the following shorthands: 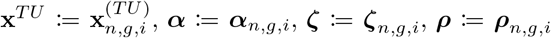, and *D*_*g,i*_ := *D*.

The DM distribution is the standard observation model of an isoform group vector **x**^(*TU*)^ [18, 30] and only requires the concentration ***α***. Its probability mass function is given by

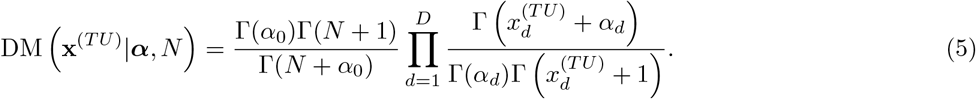

Here, 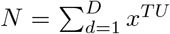 denotes the total number of exon-junction counts observed in the isoform group and 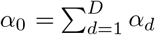 is the sum of the concentration. Each concentration parameter must satisfy *α*_*d*_ *>* 0. Because the DM distribution is conditional on the total number exon-junction counts *N*, an all-zero vector **x**^(*TU*)^ = **0** with *N* = 0 has probability one independently of ***α***. It therefore contributes a zero log-likelihood and provides no signal for learning the decoder parameters.

In contrast the ZANIDM, is a finite mixture distribution [33], that enables modelling the case **x**^*TU*^ = **0** directly by applying the excess-of-zeros parameter that denotes for each exon junction *d* the probability of no exon junction counts *ζ*_*d*_ ∈ [0, 1]. This excess-of-zeros parameter ***ζ*** is used to define the mixture weights 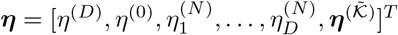 of ZANIDM as functions of the excess-of-zero parameters. A mixture weight quantifies the relative importance of a specific event scenario. Four different event scenarios can be defined for the ZANIDM. In case 1, all *D* exon junctions have counts greater than zero 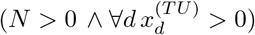 which is covered by the mixture weight

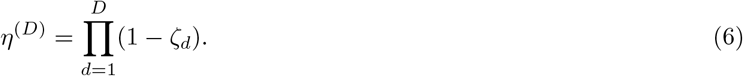

The opposite scenario is the event the event in which no exon-junction counts are observed (*N* = 0 and **x**^(*TU*)^ = **0**) constituting case 2 which has the mixture weight

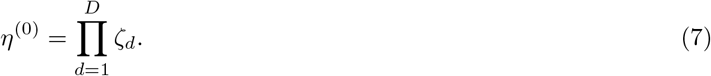

If *D* − 1 exon junctions are zero-inflated and *N >* 0, one exon junction will be *N*-inflated (case 3). This event scenario is weighted by

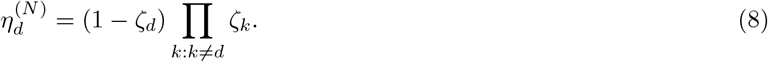

All other events are instances of the case where *N >* 0 and at most *D* − 2 exon junctions are zero-inflated (case 4). Therefore, 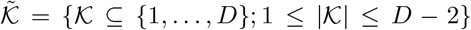 is the set of all subsets *K* of *{*1, …, *D}* with cardinality |*K*| ∈ *{*1, …, *D* − 2*}* where a subset *K* gathers all exon junctions with zero counts excluding the cases where exactly *D* and *D* − 1 exon junctions are zero-inflated. The mixture weight for a subset 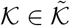 is then obtained by

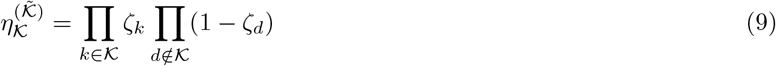

such that all mixture weights of these subsets are given by the vector 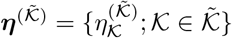. As additional constraint for each subset *K*, a truncated sum of concentration parameters is defined by 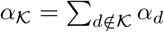. Using these four cases, the probability mass function (PMF) of the ZANIDM is given as mixture distribution by

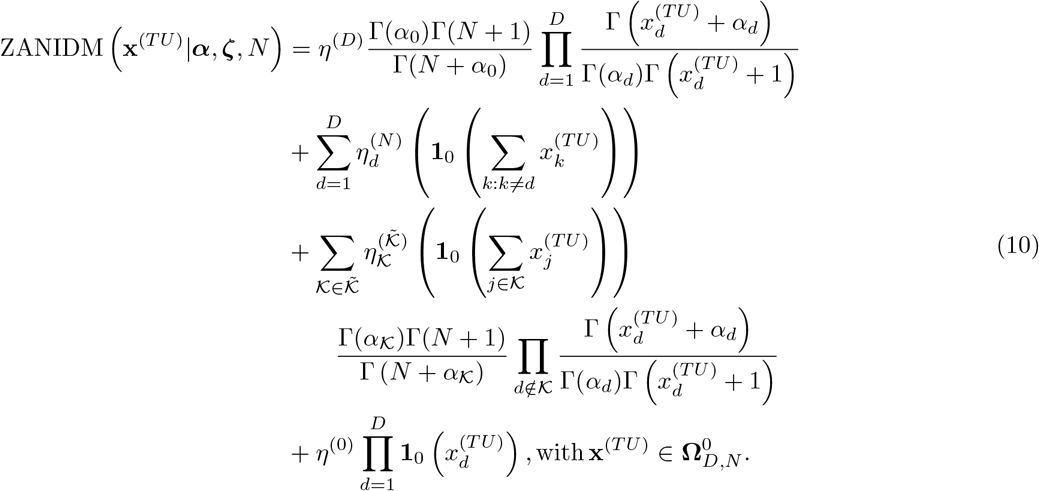

Hereby, 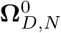 indicates the expansion of the support to the zero-inflation scenario. However, the computation of the ZANIDM observation model is hard to parallelise due to the burdensome construction of the mixture weights ***η*** and the four mixture scenarios. Therefore, tuVI provides for training an efficient heuristic of the log-likelihood that also takes the potential zero-inflation case into account while easing the computational burden and maintaining the latent space geometry (Supplementary Note 5). Therefore, tuVI is enabled to compute for an isoform group vector **x**^(*TU*)^, a logit of the excess-of-zero parameters for each exon junction

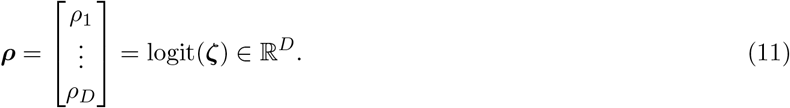

To account for the case of no exon junction counts **x**^(*TU*)^ = **0**, *N* = 0, we calculate for the isoform group for each mini-batch *B* of size |*B*| = *M* an average number of total exon junction

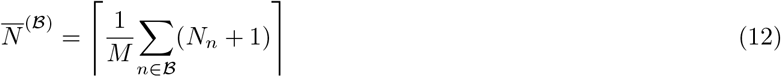

that is at least 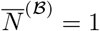. In the case of no exon junction counts **x**^(*TU*)^ = **0**, *N* = 0, this average count 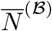 is used to compute the term

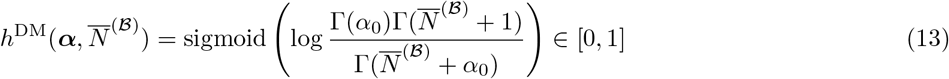

taking the parameters of the DM into account. With this term, we define a heuristic surrogate inspired by the ZANIDM log-likelihood as well as the general zero-inflation log-likelihood as

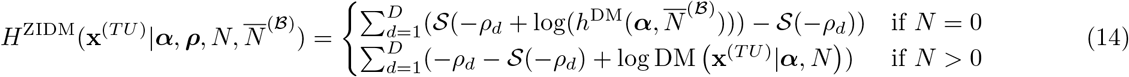

where *S*(·) is the softplus function. We refer to *H*^ZIDM^ as the heuristic ZIDM score and enable tuVI to use it in place of the expected log-likelihood to accelerate training. Because this score is not the logarithm of a normalised probability distribution, the resulting training criterion is a surrogate variational objective rather than a strict ELBO. In contrast to the DM log-likelihood, the heuristic ZIDM score permits gradients to propagate from all-zero intron-group observations. We compared the latent cell embeddings obtained by tuVI using the ZANIDM or the heuristic ZIDM as described in Supplementary Note 5.

### 4.3 Transcriptomic Regulation Variational Inference

Transcriptomic Regulation Variational Inference (TRVI) is a multimodal variational autoencoder that jointly learns cell embeddings from single-cell gene expression (GE) and alternative-splicing-induced transcript usage (TU) profiles. TRVI combines shared and modality-specific latent representations [53] with a cell-specific, modality-relevance-weighted mixture-of-experts variational posterior.

#### 4.3.1 Generative model with shared and modality-specific latent spaces

TRVI uses two types of latent variables: shared and private. The shared latent variable **Z** ∈ ℝ^*N ×L*^ denotes the joint gene expression and transcript usage (GE–TU) cell embeddings whereas the private cell embeddings **W** = {**W**^(*m*)^ ∈ ℝ^*N ×L*^}_*m*∈*M*={*GE,TU*}_ are introduced to capture patterns that are modality-specific (P–GE, P–TU). The priors over the shared and private latent variables are assumed to factorise independently. The generative model of TRVI is then given by

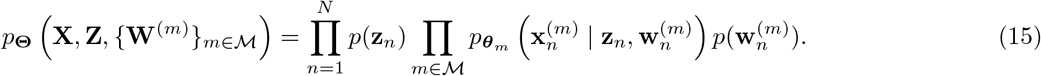

where *p*(**z**_*n*_) = *N* (**z**_*n*_ | **0, I**_*L*_), 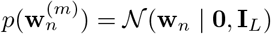 are the respective priors on the latent cell embeddings and 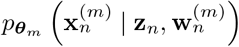 the data likelihood for modality *m* with ***θ***_*m*_ being the parameters of the deep neural network of the respective modality decoder. The set of all parameters of the generative model is denoted by **Θ** = *{****θ***_*GE*_, ***θ***_*TU*_ *}* For gene expression *m* = *GE*, the data likelihood factorises across *G*_GE_ genes as

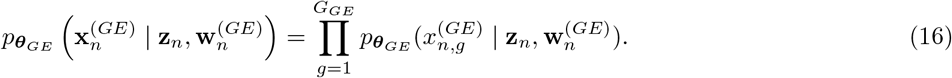

Here, 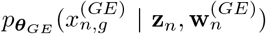 is the observation model of a gene expression 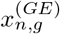 in cell *n* similar to scVI [29]. TRVI supports Gaussian, negative-binomial (NB), and zero-inflated negative-binomial (ZINB) observation models. All TRVI analyses reported here used the ZINB observation model. For transcript usage data *m* = *TU*, the data likelihood across spliced genes and their isoform groups

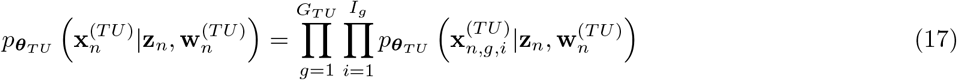

where 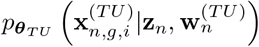 is the observation model of an isoform group vector 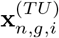. TRVI supports the same transcript usage observation models as tuVI (DM, ZANIDM) and the heuristic ZIDM. All TRVI analyses reported used the heuristic ZIDM objective.

#### 4.3.2 Modality-relevance-weighted mixture-of-experts variational posterior

TRVI uses a cell-specific modality-relevance-weighted mixture-of-experts variational posterior for the joint GE–TU cell embeddings and modality-specific Gaussian variational posteriors for the private P–GE and P–TU cell embeddding. Conventional mixture-of-experts variational posteriors combine the modality-specific experts using uniform mixture weights [38, 53] ([38, 53]. In contrast, TRVI infers the relative contribution of each modality separately for each cell through the modality-relevance weight 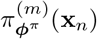. This modality-relevance-weighted variational posterior of a GE–TU cell embedding is then obtained by

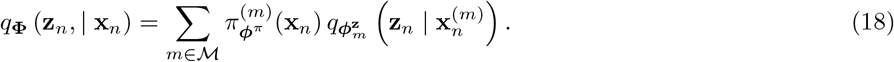

For a set of modalities ℳ = {GE, TU}, this therefore defines a two-component Gaussian mixture. The GE-derived and TU-derived components constitute the shared gene expression (S–GE) and shared transcript usage (S–TU) posterior components 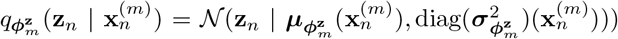, respectively. The modality-relevance weight is constrained by 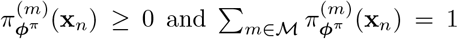. The modality-specific cell embeddings (P–GE, P–TU) use diagonal Gaussian posteriors 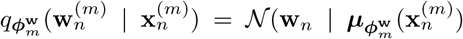, diag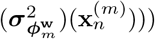. Under the corresponding mean-field factorisation, the full variational posterior of TRVI for a data point 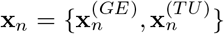 is obtained by

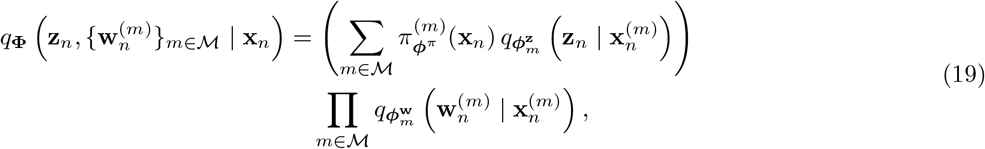

where 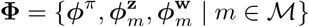 denotes the set of all variational parameters.

#### 4.3.3 TRVI training objective

The parameters (**Φ, Θ**) of TRVI are optimised using a cell-specific, modality-weighted extension of the *K*-sample MMVAE+ objective [53] (derivation of lower bound on log evidence (ELBO) is provided in Supplementary Note 3). For a single data point 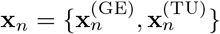, the objective of TRVI is given by

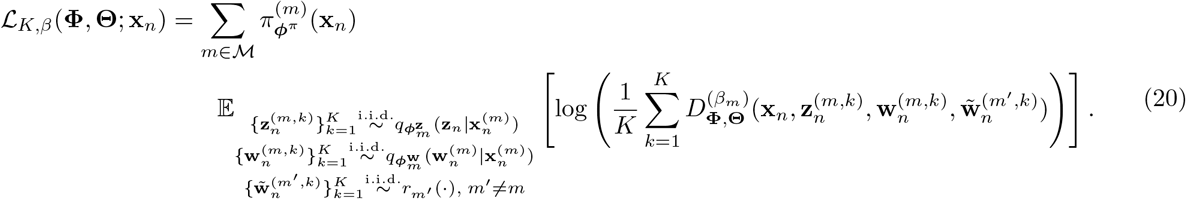

For each source modality *m*, the expectation is taken over *K* independently sampled latent-variable tuples. For sample *k*, the shared and private latent variables 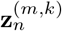 and 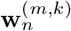 are sampled from the corresponding modality-specific variational posteriors. For the target modality *m*^*′*^ ≠ *m*, the private latent variable 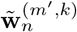 is sampled from the auxiliary distribution 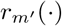 which is set to the prior. Consequently, cross-modal reconstruction of modality *m*^*′*^ cannot access the private representation inferred from 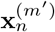 and must instead rely on information encoded in the shared latent variable. This construction discourages shared biological information from being encoded in the private latent spaces. The sample specific importance ratio is obtained by

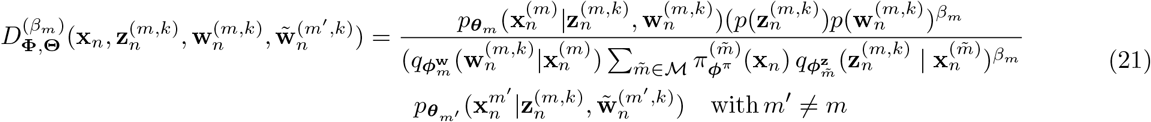

where the sum in the denominator is the probability density of the modality-relevance-weighted mixture-of-experts variational posterior evaluated at 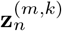. For *β*_*m*_ = 1 this results in a valid *K*-sample lower bound on the log-evidence.

During KL warm-up, *β*_*m*_ controls the strength of regularisation of the shared and private latent variables and is gradually increased to its final value *β*_*m*_ = 1. We used *K* = 10 Monte Carlo samples for all TRVI models.

#### 4.3.4 Cell-specific modality-relevance weights

To compute the cell-specific modality-relevance weights 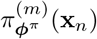 for the modality-relevance-weighted mixture-of-experts variational posterior 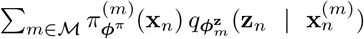, the deep neural network 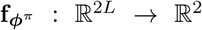 with parameters ***ϕ***^*π*^ receives the concatenated posterior means produced by the S–GE and S–TU encoders and returns two modality-specific logits. The cell-specific modality-relevance weights are then obtained by applying the softmax function yielding

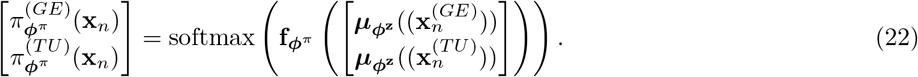

The softmax ensures that 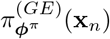 and 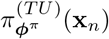 follow the constraints

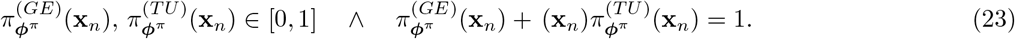

The modality-relevance weights can therefore be interpreted as cell-specific mixture probabilities that quantify the relative contributions of the S-GE and S-TU component for representing each cell in the joint GE–TU latent space. Thus, 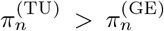 indicates that transcript usage contributes more strongly than gene expression to the joint GE-TU embedding of cell *n*.

For the Tabula Muris analyses, cell-type-, tissue- and atlas-level modality weights were calculated as averages of the cell-specific modality-relevance weights: e.g. for a cell type, this results in the average of the cell-specific modality modality-relevance weights across cells of that cell type.

#### 4.3.5 Likelihood scaling and modality-relevance weight regularisation

To ensure that neither modality dominates the optimisation of TRVI’s objective, log-likelihood scaling and regularisation of the average modality-relevance weights is applied. To adjust for differences in the magnitude of modalities the log-likelihood of the transcript usage modality is scaled during each training batch ℬ by the mean ratio of the gene expression log-likelihood to the transcript usage log-likelihood. Furthermore, to reduce the sensitivity of the inferred modality-relevance weights to stochastic optimisation and parameter initialisation, we penalise their aggregated average per batch via two competing regularisation techniques. Based on load balancing and entropy regularisation of mixture-of-experts variational autoencoders [54], we compute per mini-batch ℬ, the mean weight assigned to modality *m* as

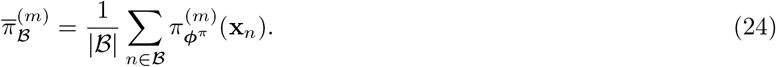

With that we first define the load balancing regularisation term

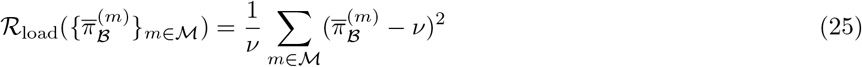

and set *ν* = 0.5 to prevent TRVI from collapsing an entire modality in the modality-relevance-weighted mixture-of-experts variational posterior and to encourage it to rely on both modalities. However, to prevent TRVI from learning an equal modality-relevance weights independent from the information content of the modalities, we also introduced the entropy regularisation

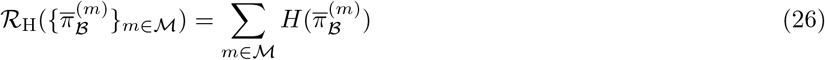

on the mini-batch mean modality-relevance weights. This entropy regularisation encourages sharper decisions between the modalities. Thus, the total regularisation of the aggregated modality-relevance weights is given by

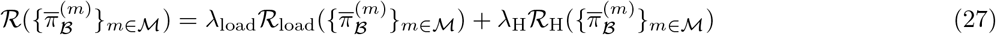

which is added to the negative TRVI objective during stochastic minimisation. Together, this regularisation constrains the batch-level of the modality-relevance weighting network and was intended to improve the stability and reproducibility of the inferred cell-specific modality-relevance weights.

### 4.4 Deep neural network architectures

The architecture of all deep neural networks applied for tuVI and TRVI is fully-connected. As building block for each neural network we designed a fully-connected module consisting of a linear layer, batch normalisation, a rectified linear unit (ReLU) as non-linearity, and a dropout layer which is based on the `scvi.nn.FCLayers` class implemented in scvi-tools (https://docs.scvi-tools.org/en/stable/; [6]). The output layer of each network comprised a linear transformation followed, where required, by an output-specific transformation, such as a softmax function for the cell-specific modality-relevance weights. The numbers of hidden layers and units, dropout rates, latent-space dimensions and other model-specific architectural hyperparameters are reported in Supplementary Tab. 1 and 3.

### 4.5 Model implementation and training of tuVI and TRVI

tuVI and TRVI were implemented using PyTorch ([55]). The weights of the linear layers were initialised using He initialisation ([56]). The parameters of the TRVI output layer that computes the modality-relevance weight logits were initialised to zero, resulting in equal initial modality-relevance weights, 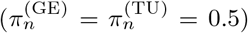 for every cell at the start of training.

The negative training objectives of tuVI and TRVI were minimised using the AdamW optimiser. (Training hyperparameters in Supplementary Tab. 1 and 3). To improve numerical stability during backpropagation, the global gradient norm was clipped to a maximum value of 1.0. Early stopping based on the validation loss was applied to limit overfitting. The optimiser settings, batch sizes, learning rates, maximum numbers of epochs and early-stopping parameters are reported in Supplementary Tab. 1 and 3.

TRVI training was divided into three phases. During the first 50 epochs, the KL-divergence contribution was gradually increased to its final value via KL-warm-up, while the modality-relevance weights were fixed at 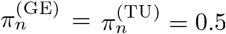. The modality-relevance weights were subsequently made learnable, and their regularisation was warmed up over a further 50 epochs. Then, training continued with the final objective until the early-stopping criterion was met or the maximum number of epochs reached.

### 4.6 Training and model specification of scVI

scVI was trained using its implementation in scvi-tools (https://scvi-tools.org/, [6]). The predefined training, validation and test partitions were specified by cell indices in a combined AnnData object. The training dataset was used to optimise the model parameters, the validation dataset was used for early stopping and model selection, and the test dataset was held out until final evaluation. The architecture and training hyperparameters applied are reported in Supplementary Tab. 2.

### 4.7 Model selection and random-seed evaluation

To select the model and assess the robustness to different random initialisations, tuVI and TRVI are evaluated using the mean per-cell conditional negative log-likelihood on the held-out test dataset. For tuVI, the AS-induced transcript usage test data 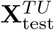 are used to infer the *N* cell embeddings **z**_*n*_. For tuVI trained with seed *s*, the NLL is then computed as

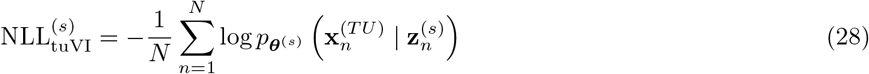

using the observation model (DM, ZANIDM, heuristic ZIDM) chosen. Because DM, ZANIDM and ZIDM define different log-likelihood functions, their absolute NLL values are not directly comparable. We therefore used NLL only to assess variation among seeds within the same observation model and to select one seed per model. Across seeds, the tuVI model with the lowest NLL score is selected for downstream tasks.

For TRVI, the conditional per-cell NLL is computed for the gene expression test data 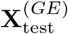 as well as for the AS-induced transcript usage data 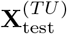. Thus, for every test cell *n*, the modality-relevance-weighted mixture-of-experts variational posterior inferred the joint GE–TU embedding 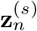, private gene expression (P-GE) embedding 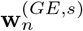 and private transcript usage (P-TU) embedding 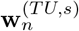 under seed *s*. Then, the gene expression NLL is computed as

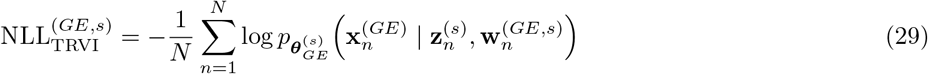

and for transcript usage data as

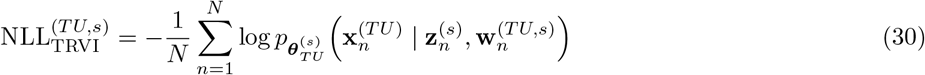

The TRVI model selected for visualisations and downstream tasks is the one with the strongest balanced performance across the two modalities. However, the modality-relevance weights (e.g. for cell types, or whole dataset) are always reported together with the distribution over all seeds.

### 4.8 Evaluation of learnt cell embeddings

To evaluate the recovery of annotated cell types in the cell embeddings learnt by scVI, tuVI and TRVI, we used the scIB library [57]. For each model, we performed Leiden clustering over resolutions ranging from 0.0 to 2.0 in increments of 0.05. The optimal resolution was defined as that yielding the highest normalised mutual information (NMI) between the inferred clusters and the reference cell-type annotations. NMI quantifies the agreement between these two partitions. Using the same NMI-optimised clustering, we calculated the adjusted Rand index (ARI), which measures agreement between the inferred clusters and reference annotations while correcting for agreement expected by chance. We additionally calculated the cell-type average silhouette width ASW_cell_ directly from the latent embeddings and reference annotations. This metric quantifies within-cell-type cohesion relative to separation from other cell types and was rescaled from ([-1,1]) to ([0,1]). Higher values of all three metrics indicate better conservation of the annotated biological structure. The average biological score was calculated as their arithmetic mean

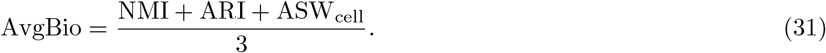

### 4.9 Evaluation of clustering and latent space geometry

To evaluate the clustering and the geometry in an inferred latent space of tuVI or TRVI, a Euclidean distance matrix 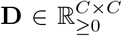 was calculated. Hereby, *C* can either be the number of cell type centroids or individual cells dependent on the setting. The Euclidean distance matrix for a pair (*i, j*) of latent cell embeddings or cell type centroids is then given by

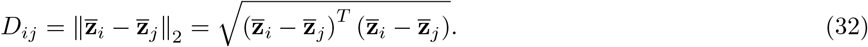

Distance matrices were then ordered according to cell types or lineages to inspect the clustering in the latent space and visualised as heatmaps.

### 4.10 Evaluation of joint latent space geometry robustness of TRVI

To evaluate the robustness of the joint gene expression and transcript usage (GE–TU) latent space of TRVI to random seed initialisations, we quantified global agreement in pairwise cell-type distances, constructed a consensus distance matrix, measured the variability of individual distances and evaluated the reproducibility of local nearest-neighbour relationships.

For seed *s*, let ℐ_*c*_ denote the set of cells annotated as cell type *c*, with *N*^(*c*)^ = |ℐ_*c*_|. The centroid of cell type *c* in the joint GE–TU representation was defined as

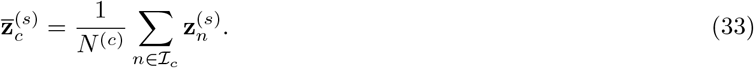

Given *C* cell types, the Euclidean distance matrix 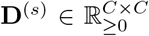 for seed *s* was then computed element-wise for each cell type centroid pair (*i, j*) as

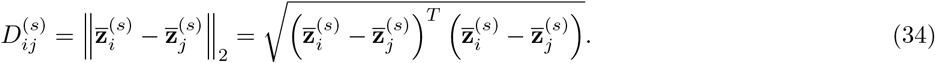

To determine whether different seeds recovered the same rank ordering of cell-type relationships, we vectorised the strict upper triangle of each distance matrix, 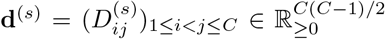, and constructed a seed-similarity matrix **R** *∈* [−1, 1]^*S×S*^. Each entry was the Spearman rank correlation between the distance vectors for a pair of seeds *s* and *t* obtained by

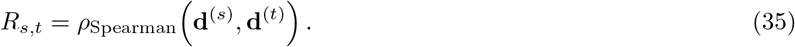

We also reported the median Spearman correlation and its range. The reproducibility of individual, scale-independent cell-type distances was assessed using a consensus distance matrix. To account for seed-specific differences in the overall scale of the latent space, each seed distance matrix **D**^(*s*)^ was divided by the median of its strict off-diagonal entries

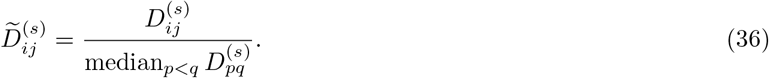

Then, the consensus distance matrix 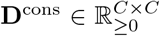 was defined element-wise for each cell type pair (*i, j*) as the median normalised distance across seeds for

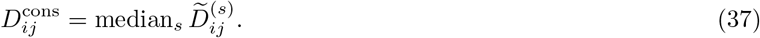

To identify cell-type relationships that were more sensitive to initialisation, we calculated an interquartile-range matrix 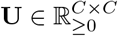 from the normalised distances computed element-wise for each cell type pair (*i, j*) as

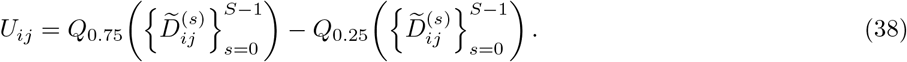

To assess the preservation of local cell-type neighbourhoods in the GE–TU latent space across seeds, we constructed a five-nearest-neighbour graph from the corresponding cell-type distance matrix and represented its edges by a binary adjacency matrix **A**^(*s*)^ for each seed *s*. The support of an edge between cell types *i* and *j* was defined as

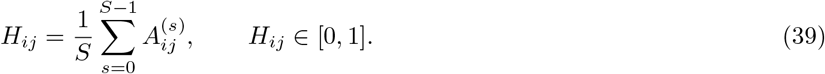

Thus, *H*_*ij*_ = 0.7 for example indicates that the neighbourhood edge was recovered in seven of the ten seed-specific graphs. For visualisation, nodes were positioned using the first two principal components of their consensus distance profiles and coloured by broad lineage.

### 4.11 Differential gene expression analysis

Differential gene expression analysis in *Crecerelle* was performed separately for each *Tabula Muris* tissue using Scanpy ([40]). A nearest-neighbour graph was constructed from either the gene expression latent cell embeddings inferred by scVI or the joint GE–TU latent cell embeddings inferred by TRVI using `sc.pp.neighbors`. The corresponding latent cell embedding was specified using `use_rep` and 15 nearest neighbours were chosen. Cells were subsequently partitioned using Leiden clustering `sc.tl.leiden` with the optimal resolution.

Differentially expressed genes were identified from the log-normalised gene expression matrix using `sc.tl.rank_genes_groups` with `method=“wilcoxon”`. For each Leiden cluster, gene expression was compared with that in all remaining cells using the default reference group. All other parameters were retained at their default values. Adjusted p-values *p*_*adj*_ were yielded using the Benjamini–Hochberg procedure, and genes with (*p*_adj_ *<* 0.05) were considered significantly differentially expressed.

### 4.12 Differential splicing analysis

Differential splicing analysis in *Crecerelle* was performed separately for each tissue in *Tabula Muris* using the approach implemented in scQuint [30], which employs the isoform group (intron group) representation introduced by LeafCutter [18]. Using Scanpy [40], a nearest-neighbour graph (`sc.pp.neighbors`) was constructed from the transcript usage cell embeddings inferred by tuVI or the joint GE–TU cell embeddings inferred by TRVI followed by Leiden clustering (`sc.tl.leiden`). When TRVI was used, the same Leiden clustering derived from the joint GE–TU cell embeddings was used for both differential gene expression and differential splicing analyses. Consequently, both analyses evaluated identical cell partitions whose definition incorporated information from both modalities. Leiden clusters that did not contain sufficient exon junction counts were excluded from downstream analysis. Each Leiden cluster was compared with the remaining clusters included in the corresponding tissue analysis.

For an exon junction *d* belonging to isoform group in a cell, the percent-spliced-in score

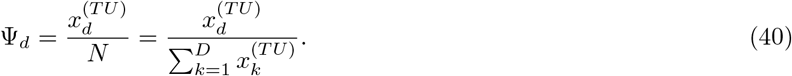

was defined as its proportion of the total exon junction counts in that isoform group. This quantity was defined only for a cell’s isoform group with present exon-junction counts *N >* 0. Isoform groups without any exon-junction counts in a cell were omitted from the calculation. For cluster *c*, the presence of a spliced isoform indicated by the change in exon-junction usage relative to all remaining cells was defined as

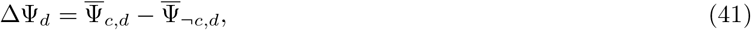

where 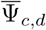 and 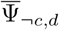 denote the mean PSI scores among cells in cluster *c* and among the remaining cells, respectively.

For each isoform group, scQuint fitted Dirichlet-Multinomial Generalised Linear Models under the null and alternative hypotheses. The alternative model included Leiden cluster membership as a predictor, whereas the null model excluded this term. Differential splicing was assessed at the isoform group level using a likelihood-ratio test. The analysis was restricted using `min_cells_per_intron_group=50` and `min_total_cells_per_intron=50`. *P*-values were adjusted for multiple testing, and an isoform group was considered differentially spliced when *p*_adj_ *<* 0.05 and at least one constituent exon junction had |ΔΨ| *>* 0.05.

Within significant isoform groups, spliced isoforms represented by exon junctions with ΔΨ *>* 0.05 were considered candidate cluster markers. For each cluster, the three highest-ranked isoform markers were then selected.

### 4.13 Functional enrichment analysis

Functional enrichment analysis within *Crecerelle* was performed separately for each *Tabula Muris* tissue using the sets of DEGs and DSGs determined for the individual cell type clusters in the tissue given. To construct the DEG set, the 100 highest-ranked genes for each cell type were considered, and genes with an adjusted *P*-value below *p*_*adj*_ *<* 0.05 and a log fold change greater than 0.5 were retained. The retained genes were pooled across all cell types within the tissue and duplicates removed. To construct the DSG set, genes associated with the significant differentially spliced intron groups identified for each cell type were pooled across the tissue and duplicates removed.

The DEG and DSG sets were analysed separately by over-representation analysis using the Python implementation of g (gprofiler-official, version 0.3.5; https://pypi.org/project/gprofiler-official/0.3.5/) through Scanpy [40, 58], with *Mus musculus* specified as the query organism. Queries were performed as unordered gene sets using the default g functional sources, annotated-gene background and g multiple-testing correction. Terms with *p*_*adj*_ *<* 0.05 were considered significantly enriched.

Significant DEG- and DSG-associated terms were compared using their database-specific term identifiers. Terms significantly enriched in both analyses were classified as shared, whereas terms significantly enriched only for the DSG set were classified as DSG-specific. DSG-specific and shared biological pathways were visualised using GO-Figure [59]. GO-Figure groups terms according to their semantic similarity and projects them into a two-dimensional semantic space, such that terms with related biological meanings are positioned closer together.

## 5 Data availability

The SmartSeq2 dataset of *Tabula Muris* containing the raw gene expression and transcript usage matrices as AnnData objects pre-processed by the bioinformatics pipeline of [30] is available at https://figshare.com/articles/dataset/scQuint_data_objects_-_Tabula_Muris/14471904. All AnnData objects processed with *Crecerelle* in this study are made available on Zenodo https://zenodo.org/records/22959716 [60].

## 6 Code availability

The source code for Crecerelle is publicly available on GitHub at https://github.com/Hollfelder-Lab/crecerelle. It can be installed via pip and git and will also be published as Python package. We provide a detailed documentation Website (https://hollfelder-lab.github.io/crecerelle/), with easily accessible notebooks and tutorials to assist with running the models and generalising these to new analyses. The processed data and trained models used in this study can be downloaded from from Zenodo https://zenodo.org/records/22959716 [60].

## 7 Acknowledgements

FMW acknowledges fellowship support from the Cambridge Trust, Evonik Stiftung, and Stiftung der deutschen Wirtschaft. IA was supported by the Bodossaki Foundation, IM was supported by the Wellcome Trust (203151/Z/16/Z) and the UKRI Medical Research Council (MC PC 17230). The work was supported by the Accelerated Adaptation programme within the Resilient Climate and Ecosystem / Engineering Ecosystem Resilience opportunity space, funded by the Advanced Research and Invention Agency (ARIA). The authors thank Chris Smith and Aishwarya Jacob for productive discussions and Clara Andiazabal, Timo Kohler, and Antonia Weberling for critical feedback on this manuscript. FMW acknowledges helpful input from the Scverse community at the Scverse Conference at Stanford 2025 (https://scverse.org/conference2025/). Some of the figures were generated using BioRender (https://biorender.com/).

## 8 Author information

### 8.1 Authors and affiliations

#### Department of Biochemistry, University of Cambridge, Cambridge, UK

Friedrich-Maximilian Weberling, Ioakeim Ampartzidis, Florian Hollfelder

#### Cambridge Stem Cell Institute, University of Cambridge, Cambridge, UK

Irina Mohorianu

#### Maastricht Center for Systems Biology and Bioinformatics, Maastricht University, Maastricht, Netherlands

Irina Mohorianu

### 8.2 Contribution

FMW conceptualised, designed, and implemented the models and downstream tasks. FMW and IA investigated the cell-type specific isoforms. IA and FMW interpreted functional features based on model outputs. FMW, IM, IA, and FH wrote the manuscript. IM and FH supervised and conceptualised the work.

### 8.3 Corresponding authors

Correspondence to Irina Mohorianu & Florian Hollfelder

## Appendix A Extended data figures

**Fig. A.1.**
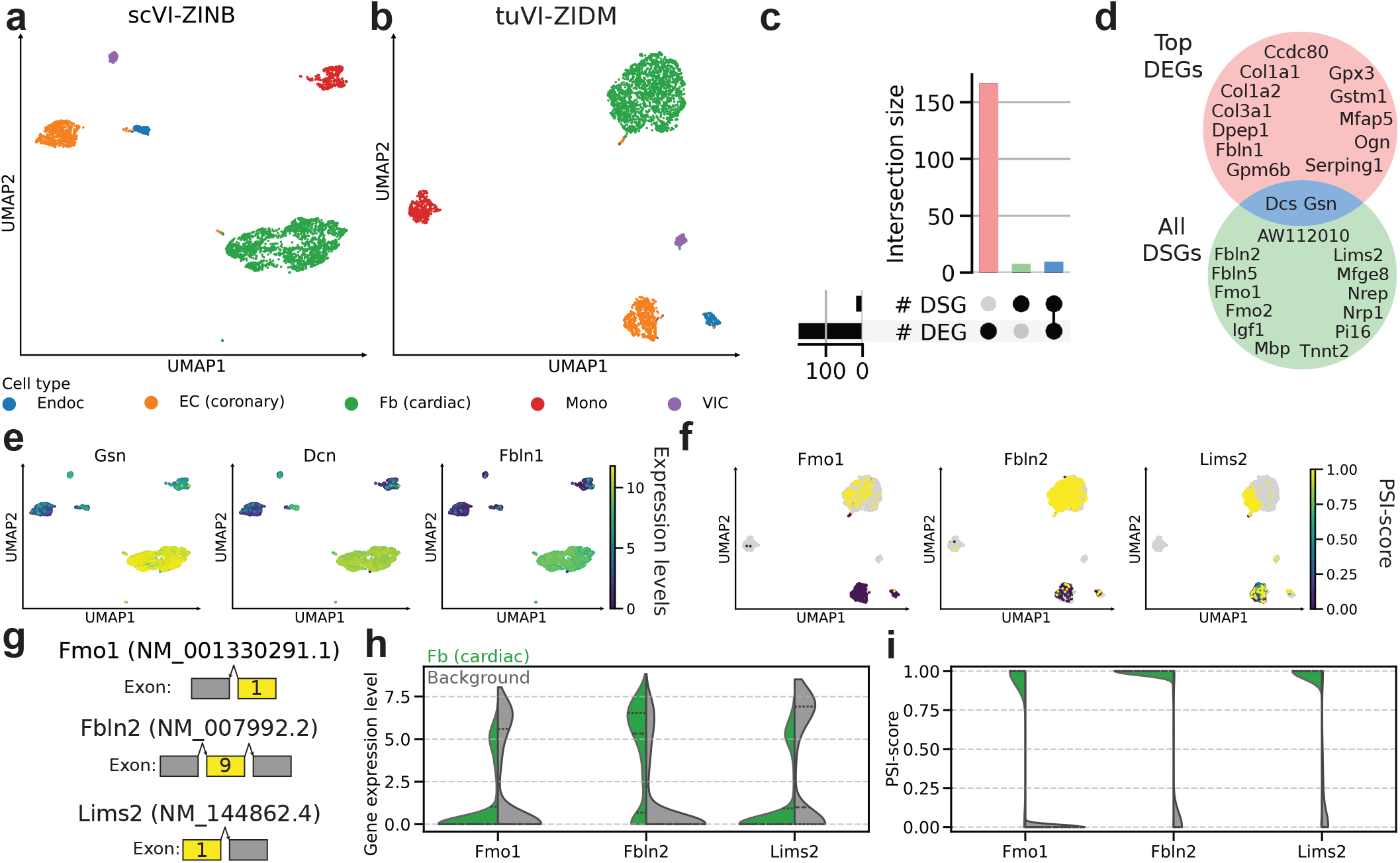
Identification of cell-type specific spliced isoform markers for cardiac fibroblasts in heart tissue, using *Crecerelle*. **a**,**b** UMAPs of gene expression cell embeddings learnt by scVI-ZINB (**a**) and transcript usage cell embeddings learnt by tuVI-ZIDM (**b**) from *Tabula Muris* (TM) heart cells, coloured by annotated cell type. Both embeddings recover the reference cell types but show modality-specific cluster structure. **c** Upsetplot comparing all significantly differentially spliced genes (DSGs) and differentially expressed genes (DEGs) for cardiac Fbs. **d** Venn diagram showcasing that only 2 of all DSGs (Dcn, Gsn) are among the top 14 ranked DEGs. **e** UMAP-projected gradient of expression for the top three DEGs (*Gsn, Dcn, Fbln1*) showcases their discriminative power as cardiac Fb cell type markers. **f** UMAP projected gradient of transcript usage for the top three DSG isoforms (*Fmo1, Fbln2, Lims2*) shows they are highly discriminative of cardiac Fbs as well. The colourbar summarises the PSI score. **g** Exon structure of the identified cell-type specific isoforms of *Fmo1, Fbln2*, and *Lims2*. **h** Violin plots displaying the gene expression level distribution of the top three DSGs across cardiac Fbs compared to all other cells shows that *Fmo1* and *Lims2* are not differentially expressed whereas *Fbln2* is upregulated. **i** Violin plots visualising the PSI-score distribution of the top three DSG isoforms across cardiac Fbs compared to all other cells indicate that the three spliced isoforms are significantly differentially spliced as well. *Fmo1* and *Lims2* are DSG only markers. Fbln2 is both a DEG and DSG.

**Fig. A.2.**
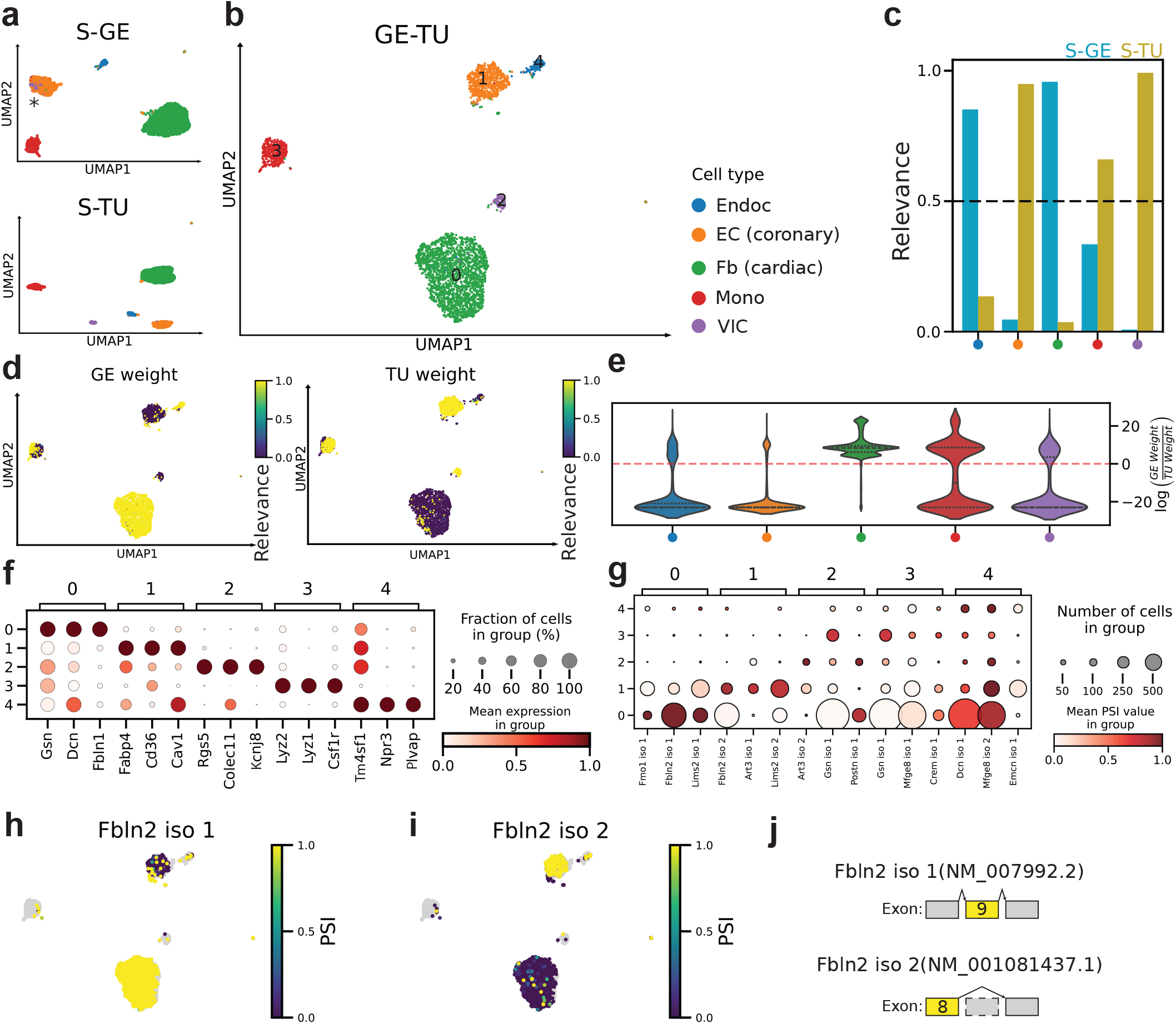
TRVI identifies cell-type specific AS programs and their interplay with expression levels in heart tissue. **a**,**b** UMAPs of the shared gene expression (S-GE) and transcript usage (S-TU) cell embeddings (**a**) and joint GE-TU cell embedding (**b**) of *Tabula Muris* heart cells, coloured by annotated cell type. Coronary endothelial cells (ECs) and valve cells (VICs), which overlap in S-GE (* in **a**), separate in S-TU and GE-TU. **c** Mean S-GE and S-TU modality-relevance weights by cell type. The dashed line indicates equal weighting. High S-TU weights for coronary ECs and VICs indicate a greater contribution of transcript usage to their joint representations. **d** GE-TU cell embedding UMAPs coloured by cell-level GE and TU modality-relevance weights. The coronary ECs and VICs have high TU weights. **e** To evaluate the robustness of the the modality-relevance weights, the violin plots depict the distribution of the log ratio of GE and TU modality-relevance weights per cell type across random seeds revealing GE (*>* 0) or TU (*<* 0) as driving factor for the inferred cell embeddings. For coronary ECs and VICs, transcript usage yields more cell-type discriminative patterns whereas for cardiac Fbs gene expression is more important. **f, g** Dot plots of the top three significantly differentially expressed genes (DEGs; **f**) and differentially spliced gene (DSG) isoforms (**g**) for Leiden clusters 0–4 (0, cardiac Fbs; 1, coronary ECs; 2, VICs; 3, monocytes; 4, endocardial cells). **h**,**i** The same DSG can be present in several cell types e.g. *Fbln2* in cardiac Fbs and coronary ECs but different isoforms of this DSG facilitate distinction as shown by UMAPs coloured by PSI for two *Fbln2* isoforms specific to cardiac Fbs (**h**) and coronary ECs (**i**). **j** Exon structures of the two mutually exclusive, cell-type-specific *Fbln2* isoforms.

**Fig. A.3.**
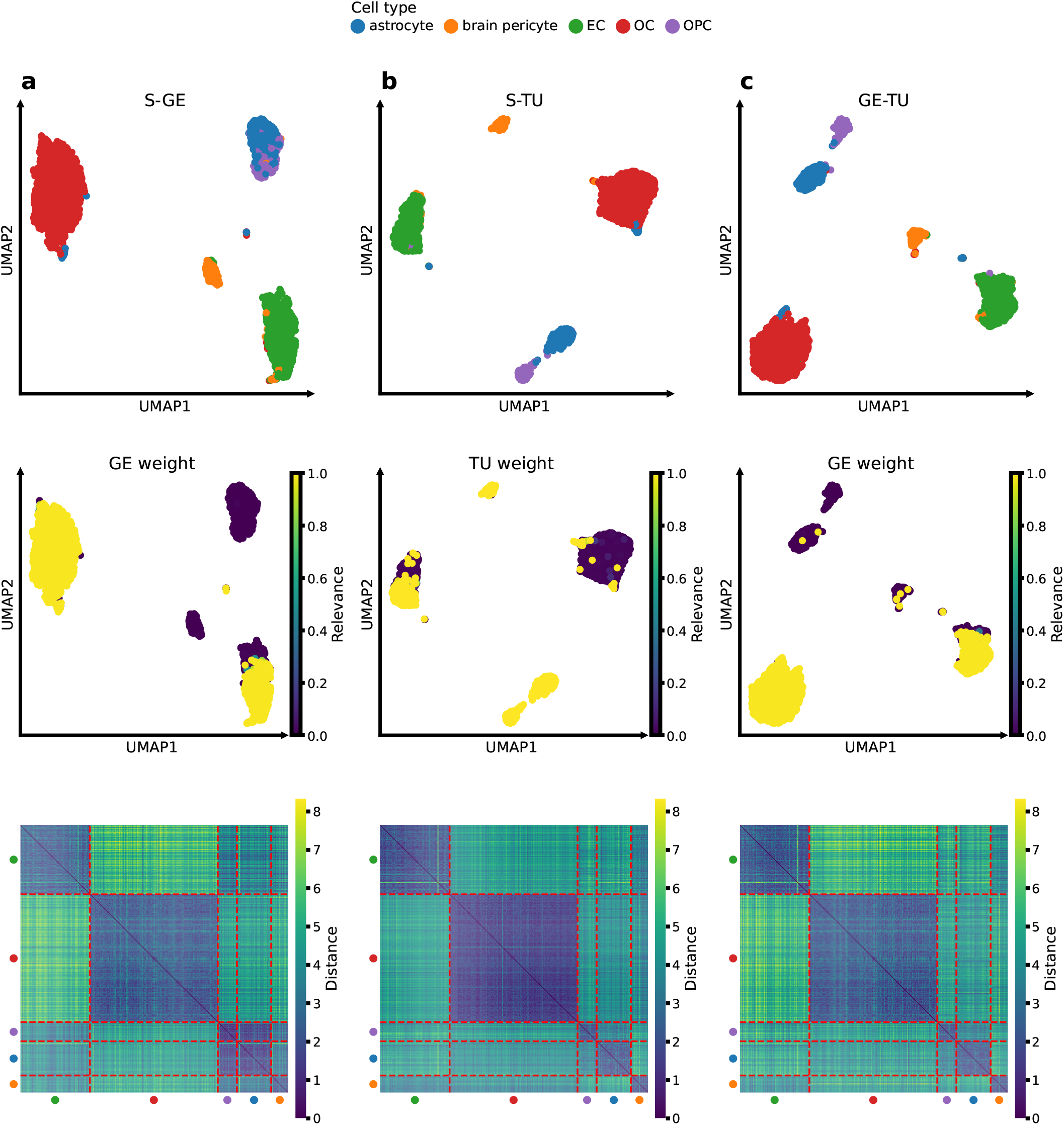
TRVI integrates complementary gene expression and transcript usage patterns to refine cell-type separation in Tabula Muris brain non-myeloid tissue. For each embedding, UMAPs show cell-type annotations (top) and learnt modality-relevance weights (middle), while heatmaps show relative Euclidean distances between cells ordered by cell type (bottom). **a** The S-GE cell embedding partially recapitulates the reference annotations, but astrocytes and oligodendrocyte precursor cells (OPCs) overlap and receive low gene expression (GE) modality-relevance weights. Their small pairwise distances further indicate weak separation. **b** The S-TU cell embedding separates astrocytes from OPCs more clearly. Both populations receive high transcript usage (TU) modality-relevance weights and form more distinct blocks in the distance matrix. **c** Modality-relevance-weighted integration of the S-GE and S-TU embeddings into the joint GE–TU embedding retains the TU-driven separation of astrocytes and OPCs and improves overall cell-type delineation.

**Fig. A.4.**
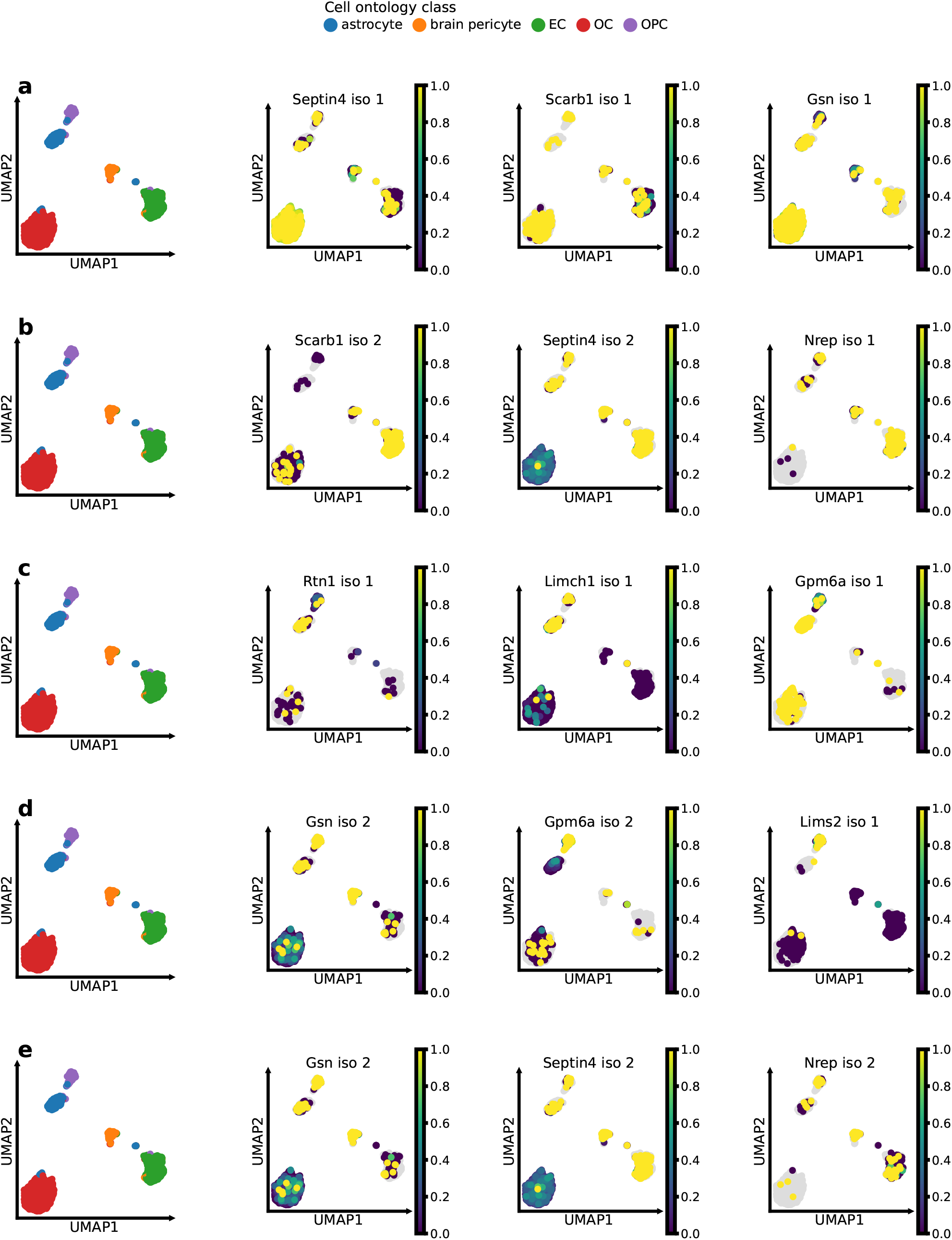
Cell-type specific sets of isoform markers can be determined using cell embeddings learnt by TRVI. In each row, the first UMAP shows the Tabula Muris reference cell-type annotation, followed by the PSI score of the three most significant differentially spliced gene isoforms for the indicated Leiden cluster. **a** OCs: (*Septin4* iso 1, *Scarb1* iso 1, *Gsn* iso 1). **b** ECs: (*Scarb1* iso 2, *Septin4* iso 2, *Nrep* iso 1). **c** astrocytes:(*Rtn* iso 1, *Limch1* iso 1, *Gpm6a* iso 1). **d** OPCs: (*Gsn* iso 2, *Gpm6a* iso 2, *Lims2* iso 1). **e** brain pericytes: (*Gsn* iso 2, *Septin4* iso 2, *Nrep* iso 2)

## Appendix B Supplementary data

**Supplementary Data Tab. 1** `trvi_degs_brain_nonmyeloid.xlsx`

Tables of all significant differentially expressed genes determined per Leiden cluster of TRVI joint GE–TU cell embeddings in *Tabula Muris* brain non-myeloid tissue.

**Supplementary Data Tab. 2** `trvi_degs_gat.xlsx`

Tables of all significant differentially expressed genes determined per Leiden cluster of TRVI joint GE–TU cell embeddings in *Tabula Muris* gonadal adipose tissue.

**Supplementary Data Tab. 3** `trvi_degs_heart.xlsx`

Tables of all significant differentially expressed genes determined per Leiden cluster of TRVI joint GE–TU cell embeddings in *Tabula Muris* heart tissue.

**Supplementary Data Tab. 4** `trvi_degs_marrow.xlsx`

Tables of all significant differentially expressed genes determined per Leiden cluster of TRVI joint GE–TU cell embeddings in *Tabula Muris* bone marrow tissue.

**Supplementary Data Tab. 5** `trvi_dsgs_brain_nonmyeloid.xlsx`

Tables of all significant differentially spliced gene isoforms determined per Leiden cluster of TRVI joint GE–TU cell embeddings in *Tabula Muris* brain non-myeloid tissue.

**Supplementary Data Tab. 6** `trvi_dsgs_gat.xlsx`

Tables of all significant differentially spliced gene isoforms determined per Leiden cluster of TRVI joint GE–TU cell embeddings in *Tabula Muris* gonadal adipose tissue.

**Supplementary Data Tab. 7** `trvi_dsgs_heart.xlsx`

Tables of all significant differentially spliced gene isoforms determined per Leiden cluster of TRVI joint GE–TU cell embeddings in *Tabula Muris* heart tissue.

**Supplementary Data Tab. 8** `trvi_dsgs_marrow.xlsx`

Tables of all significant differentially spliced gene isoforms determined per Leiden cluster of TRVI joint GE–TU cell embeddings in *Tabula Muris* bone marrow tissue.

**Supplementary Data Tab. 9** `deg_all_enriched_terms.xlsx`

Tables of all significant enrichment terms and biological pathways determined using the sets of differentially expressed genes in brain non-myeloid, gonadal adipose, heart, and bone marrow tissue of *Tabula Muris*.

**Supplementary Data Tab. 10** `dsg_all_enriched_terms.xlsx`

Tables of all significant enrichment terms and biological pathways determined using the sets of differentially spliced genes in brain non-myeloid, gonadal adipose, heart, and bone marrow tissue of *Tabula Muris*.

**Supplementary Data Tab. 11** `deg_dsg_all_enriched_terms.xlsx`

Tables of all significant enrichment terms and biological pathways that are both individually determined by the sets of differentially expressed and spliced genes in brain non-myeloid, gonadal adipose, heart, and bone marrow tissue of *Tabula Muris*.

## Footnotes

1 French word for kestrel. The acronym is derived from “**C**ellula**R** development **E**xploration of single-**C**ell gene **E**xpression and t**R**anscript usage data enabled through multimoda**L** deep generative **L**atent space**E**s”.

