## Supplementary Information for "Deep generative embeddings of gene expression and splicing reposition the interpretation of single-cell transcriptomic signatures"

### Contents

**Supplementary Tables** **3**

**Supplementary Figures** **8**

**Supplementary Note 1: Training objective of transcript usage Variational Inference (tuVI)** **42**

**Supplementary Note 2: Observation models of alternative splicing-induced transcript usage data** **43**

**Supplementary Note 3: Derivation of multi-modal Transcriptomic Regulation Variational Inference (TRVI)** **46**

**Supplementary Note 4: Sparsity and compositionality of single-cell splicing data complicate their probabilistic modelling** **49**

**Supplementary Note 5: Comparison of single-cell alternative splicing-induced transcript usage embeddings learnt by tuVI with a ZANIDM or ZIDM observation model** **50**

**Supplementary Note 6: Model evaluation of tuVI and benchmarking of its cell embeddings against scVI** **51**
6.1 Robustness to random initialisation and seed selection . . . 51
6.2 The alternative splicing-induced transcript usage cell embeddings learnt by tuVI are competitive with gene expression cell embeddings learnt by scVI . . . 51

**Supplementary Note 7: tuVI cell embeddings identify cell-type-specific isoforms that complement gene expression markers in *Tabula Muris* brain non-myeloid tissue** **52**
7.1 Clustering and differential analysis of gene expression and transcript usage embeddings . . . 52
7.2 Gene expression and transcript usage provide complementary cell-type markers . . . 52

**Supplementary Note 8: tuVI cell embeddings identify cell-type-specific isoforms that complement gene expression markers in *Tabula Muris* heart tissue** **54**

**Supplementary References** **60**

### Supplementary Tables

**Supplementary Table 1** — Training and model hyperparameters for tuVI trained on *Tabula Muris* (TM).

| Hyperparameter | Value for tuVI trained on TM |
|---|---|
| Latent space dimensionality | 10 |
| Beta | 1.0 |
| # hidden units | 256 |
| # hidden layers | 2 |
| Dropout rate | 0.1 |
| # epochs | 300 |
| Patience for early stopping | 20 |
| Initial learning rate | 0.001 |
| Batch size | 256 |
| Random seeds | 0, 1, 2, 3, 4, 5, 6, 7, 8, 9 |

**Supplementary Table 2** — Training and model hyperparameters for scVI trained on *Tabula Muris* (TM).

| Hyperparameter | Value for scVI trained on TM |
|---|---|
| Latent space dimensionality | 10 |
| # hidden units | 256 |
| # hidden layers | 2 |
| Dropout rate | 0.1 |
| # epochs | 300 |
| Patience for early stopping | 20 |
| Initial learning rate | 0.001 |
| Batch size | 128 |

**Supplementary Table 3** — Training and model hyperparameters for TRVI trained on *Tabula Muris* (TM).

| Hyperparameter | Value for TRVI trained on TM |
|---|---|
| Latent space dimensionality | 10 |
| Final Beta | 1.0 |
| # hidden units | 256 |
| # hidden layers | 256 |
| Dropout rate | 0.1 |
| # epochs | 1000 |
| # epochs KL warm-up | 50 |
| # epochs modality-relevance weight warm-up | 50 |
| Patience for early stopping | 20 |
| Initial learning rate | 0.0001 |
| Batch size | 256 |
| Random seeds | 0, 1, 2, 3, 4, 5, 6, 7, 8, 9 |

**Supplementary Table 4** — Robustness of tuVI with Dirichlet–multinomial (DM), zero-and-$N$-inflated Dirichlet–multinomial (ZANIDM) and zero-inflated Dirichlet–multinomial (ZIDM) observation models to random initialisation benchmarked on *Tabula Muris* (TM). Values represent the mean ± standard deviation of the per-cell conditional negative log-likelihood (NLL) across the held-out test set. Bold indicates the lowest mean NLL within each observation model. Because the likelihood definitions differ, NLL values should not be compared across observation models.

| | Observation model | | |
|---|---|---|---|
| Random seed | DM | ZANIDM | ZIDM (heuristic) |
| 0 | 65.379 ± 54.629 | 174.894 ± 78.314 | 225.665 ± 87.295 |
| 1 | 65.531 ± 54.587 | 174.813 ± 78.240 | **225.327 ± 87.399** |
| 2 | 65.836 ± 54.611 | 174.707 ± 77.819 | 225.975 ± 87.449 |
| 3 | 65.435 ± 54.498 | 175.227 ± 78.196 | 225.933 ± 87.493 |
| 4 | 65.523 ± 54.569 | 175.023 ± 77.798 | 226.087 ± 87.533 |
| 5 | **65.095 ± 54.505** | 174.689 ± 78.259 | 225.520 ± 87.378 |
| 6 | 65.178 ± 54.593 | **174.495 ± 77.852** | 226.051 ± 87.683 |
| 7 | 65.439 ± 54.423 | 174.816 ± 78.005 | 226.911 ± 86.761 |
| 8 | 65.264 ± 54.498 | 174.692 ± 77.745 | 225.779 ± 87.473 |
| 9 | 65.288 ± 54.576 | 174.687 ± 77.902 | 225.486 ± 87.310 |

**Supplementary Table 5** — Test-set reconstruction performance of TRVI across random-seed initialisations benchmarked on *Tabula Muris* (TM). Model performance was assessed separately for gene expression (GE) and transcript usage (TU) using the conditional negative log-likelihood (NLL) on held-out test cells. Values denote the mean ± standard deviation of the per-cell NLL distribution. Bold type indicates the seed selected for representative visualisations and downstream analyses.

| | Test-set NLL | |
|---|---|---|
| Random seed | Gene expression (GE) | Transcript usage (TU) |
| 0 | 1094.949 ± 533.816 | 294.349 ± 177.340 |
| 1 | 1094.659 ± 518.792 | 338.748 ± 210.181 |
| 2 | 1059.807 ± 485.853 | 279.720 ± 172.274 |
| 3 | 1077.182 ± 460.032 | 285.875 ± 205.508 |
| 4 | 1077.083 ± 461.200 | 259.769 ± 142.033 |
| 5 | 1104.435 ± 506.281 | 329.804 ± 268.041 |
| 6 | 1050.539 ± 443.835 | 229.408 ± 102.988 |
| 7 | 1140.276 ± 1200.166 | 258.110 ± 122.580 |
| 8 | **1044.347 ± 429.198** | **229.593 ± 104.584** |
| 9 | 1070.998 ± 466.193 | 287.122 ± 156.485 |

**Supplementary Table 6** — Evaluation of the cell embeddings learnt by scVI and tuVI with different observation models. Metrics were calculated relative to the *Tabula Muris* cell-type annotations. AvgBio denotes the mean of the normalised mutual information (NMI), adjusted Rand index (ARI) and average silhouette width (ASW). The highest value for each metric is shown in bold.

| Evaluation metric | scVI-ZINB | tuVI-DM | tuVI-ZANIDM | tuVI-ZIDM |
|---|---|---|---|---|
| NMI | **0.879** | 0.323 | 0.871 | 0.864 |
| ARI | 0.685 | 0.128 | 0.715 | **0.724** |
| ASW | **0.619** | 0.456 | 0.606 | 0.576 |
| Avg Bio | 0.727 | 0.302 | **0.730** | 0.722 |

**Supplementary Table 7** — Evaluation of the cell embeddings learnt by TRVI. Metrics were calculated relative to the *Tabula Muris* cell-type annotations. AvgBio denotes the mean of the normalised mutual information (NMI), adjusted Rand index (ARI) and average silhouette width (ASW). The highest score for each evaluation metric is shown in bold.

| Evaluation metric | S-GE | S-TU | GE-TU |
|---|---|---|---|
| NMI | 0.804 | 0.846 | **0.850** |
| ARI | 0.680 | 0.759 | **0.761** |
| ASW | 0.548 | **0.573** | 0.569 |
| Avg Bio | 0.677 | **0.726** | **0.726** |

**Supplementary Table 8** — Atlas-level modality-relevance weights learnt from *Tabula Muris*.

| Atlas-level GE weight | Atlas-level TU weight |
|---|---|
| $0.558 \pm 0.049$ | $0.442 \pm 0.049$ |

**Supplementary Table 9** — Tissue-level GE and TU mean weights learnt from *Tabula Muris*.

| Tissue | Mean GE weight | Mean TU weight |
|---|---|---|
| BAT | 0.299 | 0.701 |
| GAT | 0.699 | 0.301 |
| MAT | 0.653 | 0.347 |
| SCAT | 0.570 | 0.430 |
| Brain myeloid | 0.250 | 0.750 |
| Brain non-myeloid | 0.645 | 0.355 |
| Aorta | 0.525 | 0.475 |
| Heart | 0.662 | 0.338 |
| Lung | 0.670 | 0.330 |
| Trachea | 0.595 | 0.405 |
| Large intestine | 0.971 | 0.029 |
| Liver | 0.004 | 0.996 |
| Pancreas | 0.818 | 0.182 |
| Mammary gland | 0.394 | 0.606 |
| Skin | 0.554 | 0.446 |
| Bladder | 0.982 | 0.018 |
| Marrow | 0.542 | 0.458 |
| Spleen | 0.168 | 0.832 |
| Thymus | 0.812 | 0.188 |
| Diaphragm | 0.174 | 0.826 |
| Limb muscle | 0.212 | 0.788 |
| Tongue | 0.976 | 0.024 |

**Supplementary Table 10** — Cell-type-specific GE and TU mean weights learnt from *Tabula Muris.*

| Cell type | Organ system | Mean GE weight | Mean TU weight |
|---|---|---|---|
| MSC (adipose) | Adipose | $0.893 \pm 0.270$ | $0.107 \pm 0.270$ |
| myeloid cell | Adipose | $0.408 \pm 0.273$ | $0.592 \pm 0.273$ |
| T cell* | Adipose | $0.616 \pm 0.244$ | $0.384 \pm 0.244$ |
| OC | Brain | $0.963 \pm 0.077$ | $0.037 \pm 0.077$ |
| OPC | Brain | $0.771 \pm 0.407$ | $0.229 \pm 0.407$ |
| microglial cell | Brain | $0.279 \pm 0.293$ | $0.721 \pm 0.293$ |
| brain pericyte | Brain | $0.312 \pm 0.366$ | $0.688 \pm 0.366$ |
| astrocyte | Brain | $0.784 \pm 0.376$ | $0.216 \pm 0.376$ |
| EC* | Brain | $0.157 \pm 0.263$ | $0.843 \pm 0.263$ |
| VIC | Cardiorespiratory | $0.258 \pm 0.388$ | $0.742 \pm 0.388$ |
| Fb (cardiac) | Cardiorespiratory | $0.956 \pm 0.056$ | $0.044 \pm 0.056$ |
| Fb | Cardiorespiratory | $0.896 \pm 0.260$ | $0.104 \pm 0.260$ |
| EC (aortic) | Cardiorespiratory | $0.137 \pm 0.309$ | $0.863 \pm 0.309$ |
| Mono | Cardiorespiratory | $0.463 \pm 0.404$ | $0.537 \pm 0.404$ |
| Endoc | Cardiorespiratory | $0.185 \pm 0.384$ | $0.815 \pm 0.384$ |
| chondrocyte | Cardiorespiratory | $0.623 \pm 0.412$ | $0.377 \pm 0.412$ |
| EC (coronary) | Cardiorespiratory | $0.106 \pm 0.277$ | $0.894 \pm 0.277$ |
| SMC (bronchial) | Cardiorespiratory | $0.159 \pm 0.317$ | $0.841 \pm 0.317$ |
| adventitial cell | Cardiorespiratory | $0.947 \pm 0.117$ | $0.053 \pm 0.117$ |
| EC (hepatic sinusoid) | Digestive & metabolic | $0.015 \pm 0.019$ | $0.985 \pm 0.019$ |
| Enterocyte | Digestive & metabolic | $0.596 \pm 0.513$ | $0.404 \pm 0.513$ |
| pancreatic B cell | Digestive & metabolic | $0.890 \pm 0.311$ | $0.110 \pm 0.311$ |
| pancreatic D cell | Digestive & metabolic | $0.893 \pm 0.311$ | $0.107 \pm 0.311$ |
| pancreatic ductal cell | Digestive & metabolic | $0.404 \pm 0.497$ | $0.596 \pm 0.497$ |
| pancreatic A cell | Digestive & metabolic | $0.893 \pm 0.312$ | $0.107 \pm 0.312$ |
| Goblet cell | Digestive & metabolic | $0.779 \pm 0.409$ | $0.221 \pm 0.409$ |
| secretory cell | Digestive & metabolic | $0.774 \pm 0.406$ | $0.226 \pm 0.406$ |
| hepatocyte | Digestive & metabolic | $0.400 \pm 0.511$ | $0.600 \pm 0.511$ |
| intestinal crypt stem cell | Digestive & metabolic | $0.592 \pm 0.499$ | $0.408 \pm 0.499$ |
| stromal cell | Excretory & integumentary | $0.866 \pm 0.304$ | $0.134 \pm 0.304$ |
| Ep (luminal) | Excretory & integumentary | $0.676 \pm 0.389$ | $0.324 \pm 0.389$ |
| basal cell | Excretory & integumentary | $0.826 \pm 0.315$ | $0.174 \pm 0.315$ |
| bulge keratinocyte | Excretory & integumentary | $0.754 \pm 0.228$ | $0.246 \pm 0.228$ |
| epidermal cell | Excretory & integumentary | $0.810 \pm 0.284$ | $0.190 \pm 0.284$ |
| Mono (pro) | Immune & hematopoietic | $0.531 \pm 0.371$ | $0.469 \pm 0.371$ |
| B cell (pre) | Immune & hematopoietic | $0.236 \pm 0.321$ | $0.764 \pm 0.321$ |
| B cell (naive) | Immune & hematopoietic | $0.160 \pm 0.173$ | $0.840 \pm 0.173$ |
| B cell* | Immune & hematopoietic | $0.183 \pm 0.159$ | $0.817 \pm 0.159$ |
| macrophage* | Immune & hematopoietic | $0.422 \pm 0.279$ | $0.578 \pm 0.279$ |
| T cell (CD4+)* | Immune & hematopoietic | $0.284 \pm 0.204$ | $0.716 \pm 0.204$ |
| T cell (CD8+) | Immune & hematopoietic | $0.380 \pm 0.306$ | $0.620 \pm 0.306$ |
| DN4 thymocyte | Immune & hematopoietic | $0.473 \pm 0.253$ | $0.527 \pm 0.253$ |
| NK cell | Immune & hematopoietic | $0.347 \pm 0.267$ | $0.653 \pm 0.267$ |
| bladder urothelial cell | Immune & hematopoietic | $0.697 \pm 0.476$ | $0.303 \pm 0.476$ |
| bladder cell | Immune & hematopoietic | $0.892 \pm 0.312$ | $0.108 \pm 0.312$ |
| granulocytopoietic cell | Immune & hematopoietic | $0.701 \pm 0.442$ | $0.299 \pm 0.442$ |
| thymocyte | Immune & hematopoietic | $0.764 \pm 0.259$ | $0.236 \pm 0.259$ |
| B cell (immature) | Immune & hematopoietic | $0.314 \pm 0.256$ | $0.686 \pm 0.256$ |
| B cell (late pro) | Immune & hematopoietic | $0.678 \pm 0.265$ | $0.322 \pm 0.265$ |
| granulocyte | Immune & hematopoietic | $0.520 \pm 0.305$ | $0.480 \pm 0.305$ |
| HSC | Immune & hematopoietic | $0.363 \pm 0.309$ | $0.637 \pm 0.309$ |
| MSC | Musculoskeletal | $0.877 \pm 0.227$ | $0.123 \pm 0.227$ |
| basal cell (epidermis)* | Musculoskeletal | $0.727 \pm 0.278$ | $0.273 \pm 0.278$ |
| keratinocyte | Musculoskeletal | $0.687 \pm 0.471$ | $0.313 \pm 0.471$ |
| SC (muscle) | Musculoskeletal | $0.378 \pm 0.239$ | $0.622 \pm 0.239$ |

**Supplementary Table 11** — Mean gene expression (GE) and transcript usage (TU) modality-relevance weights by cell type in brain non-myeloid tissue of *Tabula Muris* for the best-performing TRVI model.

| Cell type | Mean GE weight | Mean TU weight |
|---|---|---|
| Astrocyte | 0.059 | 0.941 |
| Brain pericyte | 0.101 | 0.899 |
| EC | 0.628 | 0.372 |
| OC | 0.975 | 0.025 |
| OPC | 0.000 | 1.000 |

**Supplementary Table 12** — Differentially expressed genes (DEGs), differentially spliced genes (DSGs), and candidate transcript usage (TU) markers across *Tabula Muris* brain non-myeloid cell types determined from scVI and tuVIs cell embeddings respectively.

| Cell type | # DEGs only | # DSGs only | # DEGs and DSGs | Main DSG isoform candidates |
|---|---|---|---|---|
| OC (Leiden 0) | 90 | 6 | 5 | *Scarb1* (general OC), *Lims2* (subpopulation specific) |
| EC | 176 | 12 | 5 | *Scarb1*, *Septin4*, *Nrep* |
| Astrocytes | 143 | 5 | 9 | *Rtn1*, *Limch1*, *Gpm6a* |
| OC (Leiden 3) | 89 | 5 | 6 | *Scarb1* and *Septin4* for general OC |
| OPC | 182 | 6 | 11 | *Gsn*, *Gpm6a* |
| Brain pericytes | 123 | 1 | 6 | inconclusive due to sparsity |

**Supplementary Table 13** — Differentially expressed genes (DEGs), differentially spliced genes (DSGs), and candidate transcript usage (TU) markers across *Tabula Muris* heart cell types determined from scVI and tuVIs cell embeddings respectively.

| Cell type | # DEGs only | # DSGs only | # DEGs and DSGs | Main DSG isoform candidates |
|---|---|---|---|---|
| Cardiac Fbs | 166 | 6 | 8 | *Fmo1*, *Fbln2*, *Lims2* |
| Coronary EC | 149 | 5 | 11 | *Fbln2*, *Lims2*, *Art3* |
| VIC | 78 | 1 | 3 | *Art3*, *Postn* |
| Mono | 180 | 5 | 3 | *Gsn* |
| Endoc | 108 | 1 | 2 | inconclusive due to sparsity |

### Supplementary Figures

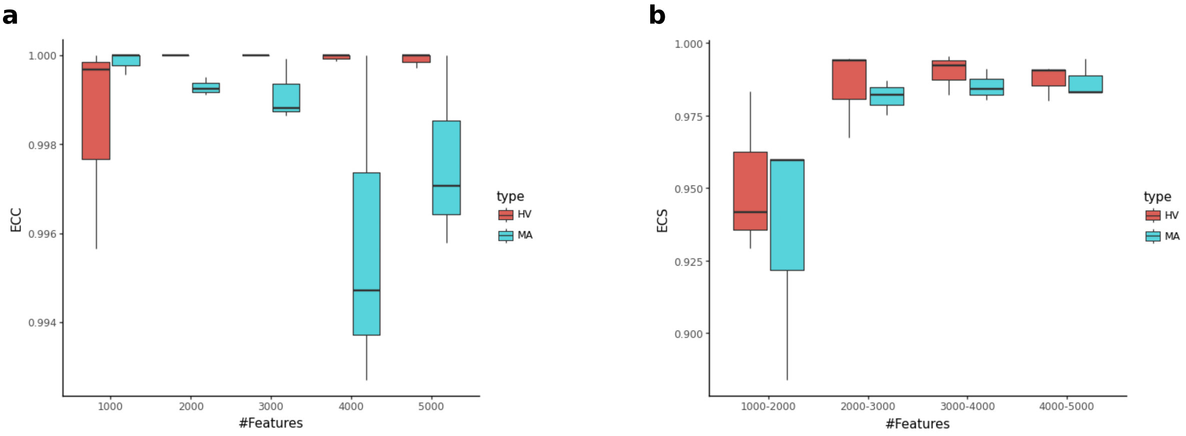

**Supplementary Fig. 1 — A stable number of highly variable genes is determined using ClustAssess on the *Tabula Muris* dataset. a** Feature stability of different sets of highly variable (HV) genes or most abundant (MA) genes are evaluated by the element-centric consistency (ECC) score using sets 1000, 2000, 3000, 4000, and 5000 sets of genes. The set of 3000 HV genes yields the best ECC score on *Tabula Muris* dataset. **b** Feature stability of different sets of HV genes or MA genes are evaluated by the element-centric similarity (ECS) score using sets 1000, 2000, 3000, 4000, and 5000 sets of genes. The best ECS is yielded by 3000 HV genes.

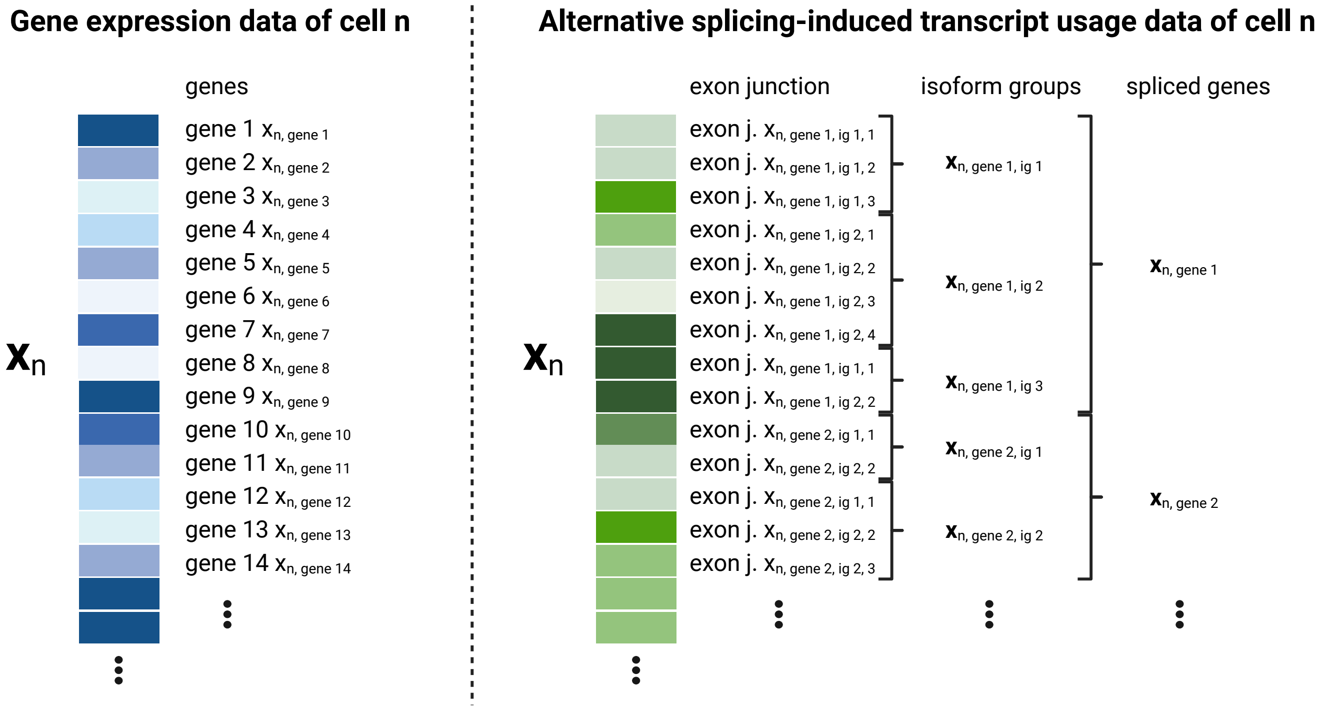

**Supplementary Fig. 2 — Single-cell alternative splicing-induced transcript usage data are compositional in contrast to gene expression data.** The gene expression vector $\mathbf{x}_n$ of cell $n$ uses genes as features and treats them independently. In contrast, an according single-cell transcript usage vector $\mathbf{x}_n$ has exon junctions (introns) as elements to quantify spliced isoforms. These exon junctions cannot be treated independently but need to be considered with reference to their corresponding isform group (intron group) which again belongs to a spliced gene. Each isoform group and each spliced gene are treated independently but since the isoform group sizes can vary a compositional structure is present in the transcript usage vector.

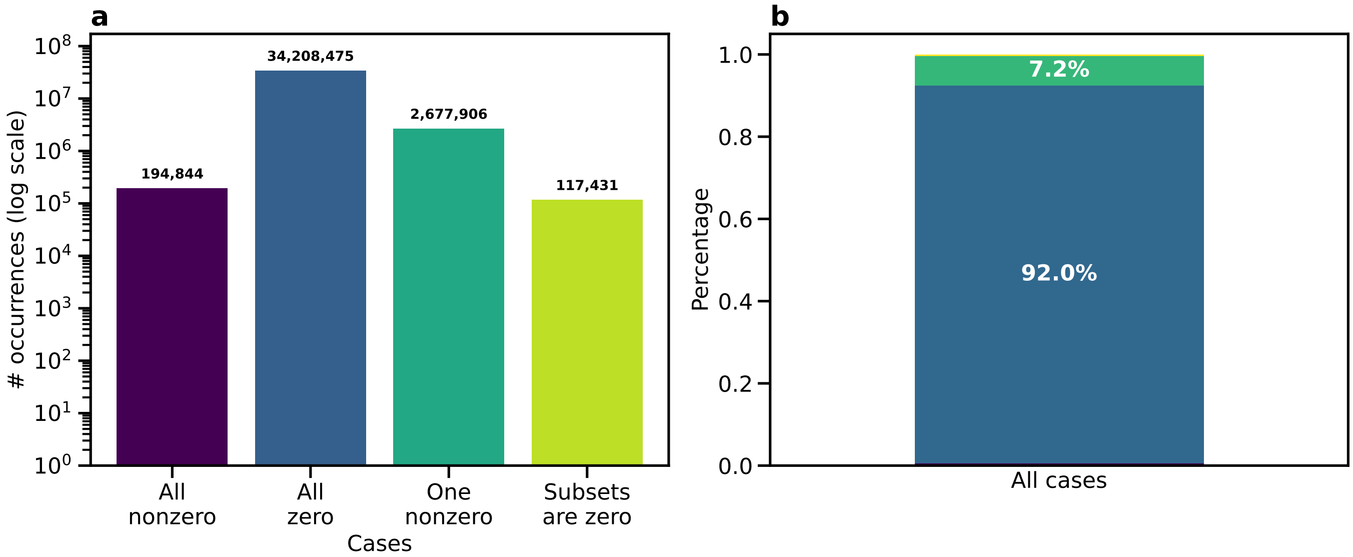

**Supplementary Fig. 3 — Ignoring the inherent sparsity of single-cell splicing data results in the loss of the majority of transcript usage profiles. a** Each isoform group count profile for each cell in the *Tabula Muris* atlas is sorted in its corresponding case. The bar chart displays the absolute number of occurrences of each case in the dataset on a log scale. The case "all zero" dominates followed by the case "one nonzero". Both cases are hard to model with a Dirichlet-Multinomial observation model. **b** The stacked bar chart displays the relative occurrence of each case in the dataset underlining that the case "all zero" which is outside of the event space of the Dirichlet-Multinomial represents the majority of transcript usage profiles.

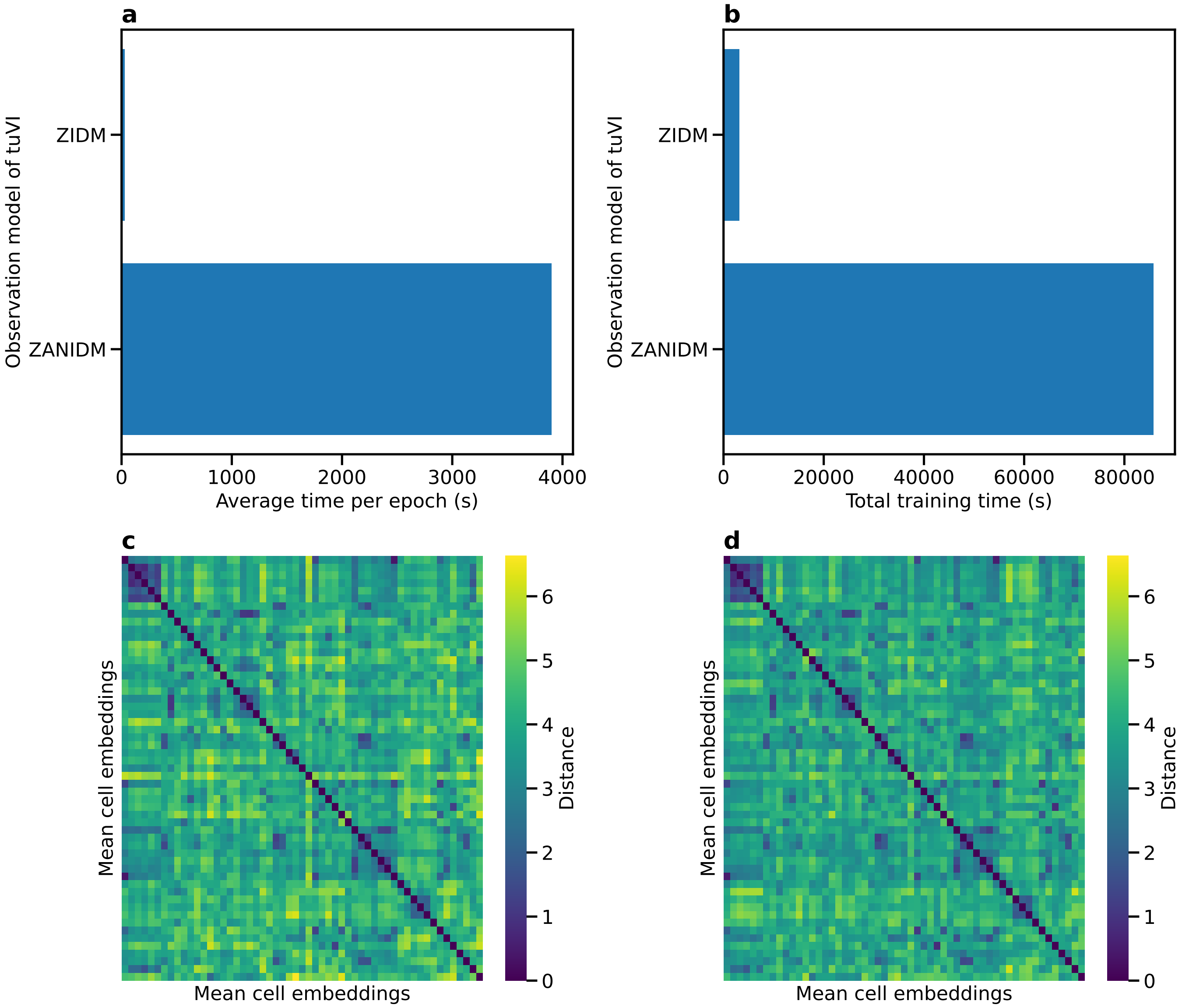

**Supplementary Fig. 4 — The heuristic ZIDM is a fast surrogate of the ZANIDM observation model log-likelihood yielding a highly similar latent space of cell embeddings. a** The bar chart displays the average time in seconds to complete a training epoch required by tuVI with a ZANIDM observation model or by tuVI with the heuristic ZIDM, a surrogate model of the ZANIDM log-likelihood. The ZIDM is 22 times faster. **b**. The bar chart depicts the total training time of tuVI-ZANIDM and tuVI-ZIDM. The total duration of the optimisation is 17 times faster for tuVI-ZIDM than for tuVI-ZANIDM. **c** The heatmap displays for tuVI-ZANIDM the Euclidean distance between the mean cell embeddings of cell type clusters. The mean cell embedding is computed by averaging all cell embeddings of a particular cell type. **d** The according heatmap is generated for tuVI-ZIDM. Comparing both heatmaps shows the relative distances between cell type clusters are preserved displaying visually that the latent space of cell embeddings of tuVI-ZIDM is highly representative of the latent space of tuVI-ZANIDM although the heursitic ZIDM is a surrogate model.

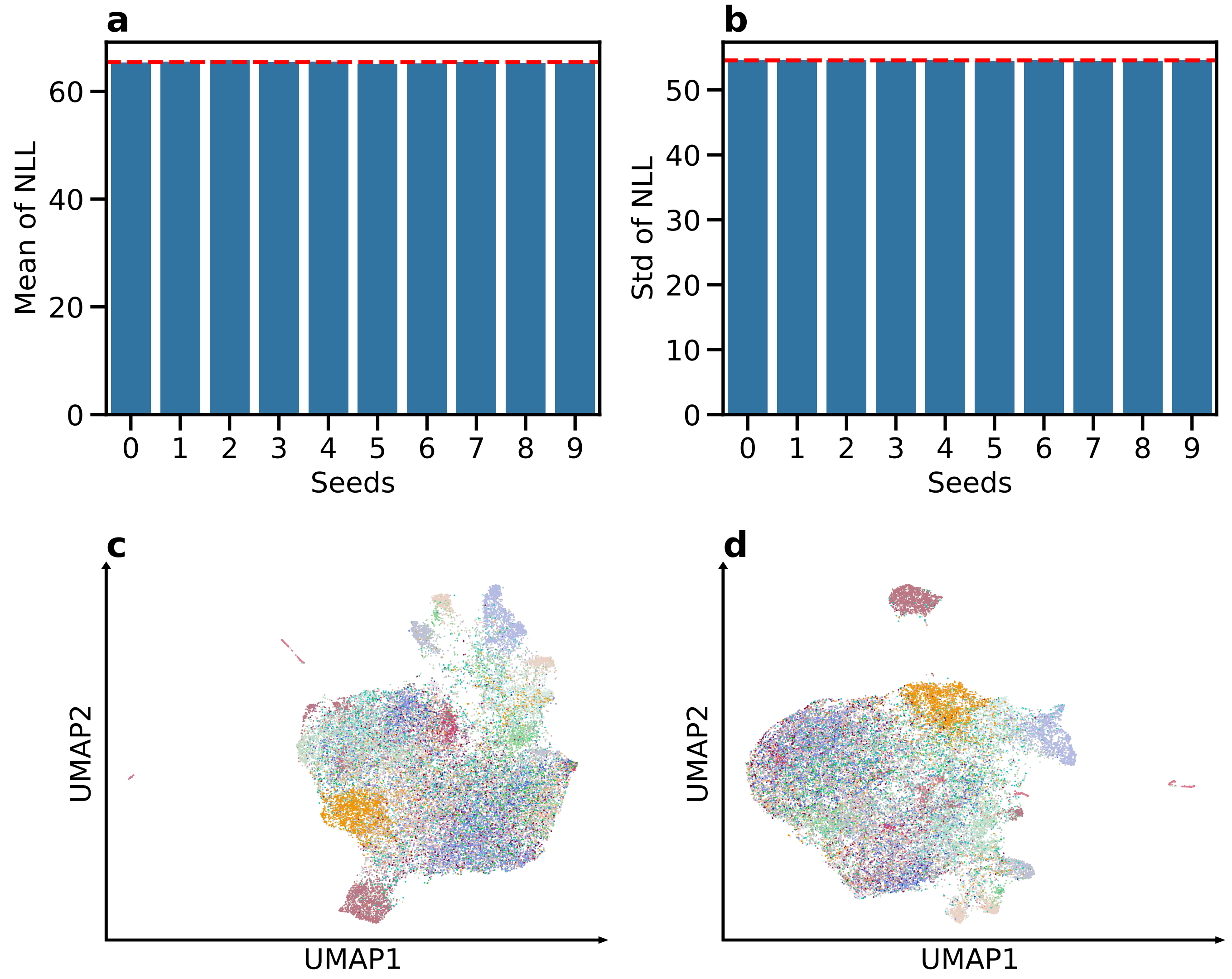

**Supplementary Fig. 5 — tuVI-DM is robust to random weight initialisation but resolves limited cell-type structure. a,b** Mean (**a**) and standard deviation (**b**) of the per-cell negative log-likelihood (NLL) on the held-out *Tabula Muris* test set for tuVI-DM models trained with ten random seeds. Red dashed lines indicate the corresponding averages across seeds. The small between-seed differences demonstrate that model performance is robust to random weight initialisation. **c,d** UMAPs of the *Tabula Muris* cell embeddings learnt by the models with the lowest (seed 5; **c**) and highest (seed 2; **d**) mean test-set NLL, coloured by annotated cell type. Both latent spaces show limited separation of cell populations, suggesting that the Dirichlet Multinomial observation model does not capture sufficient variation in alternative splicing-induced transcript usage to determine cell type clusters.

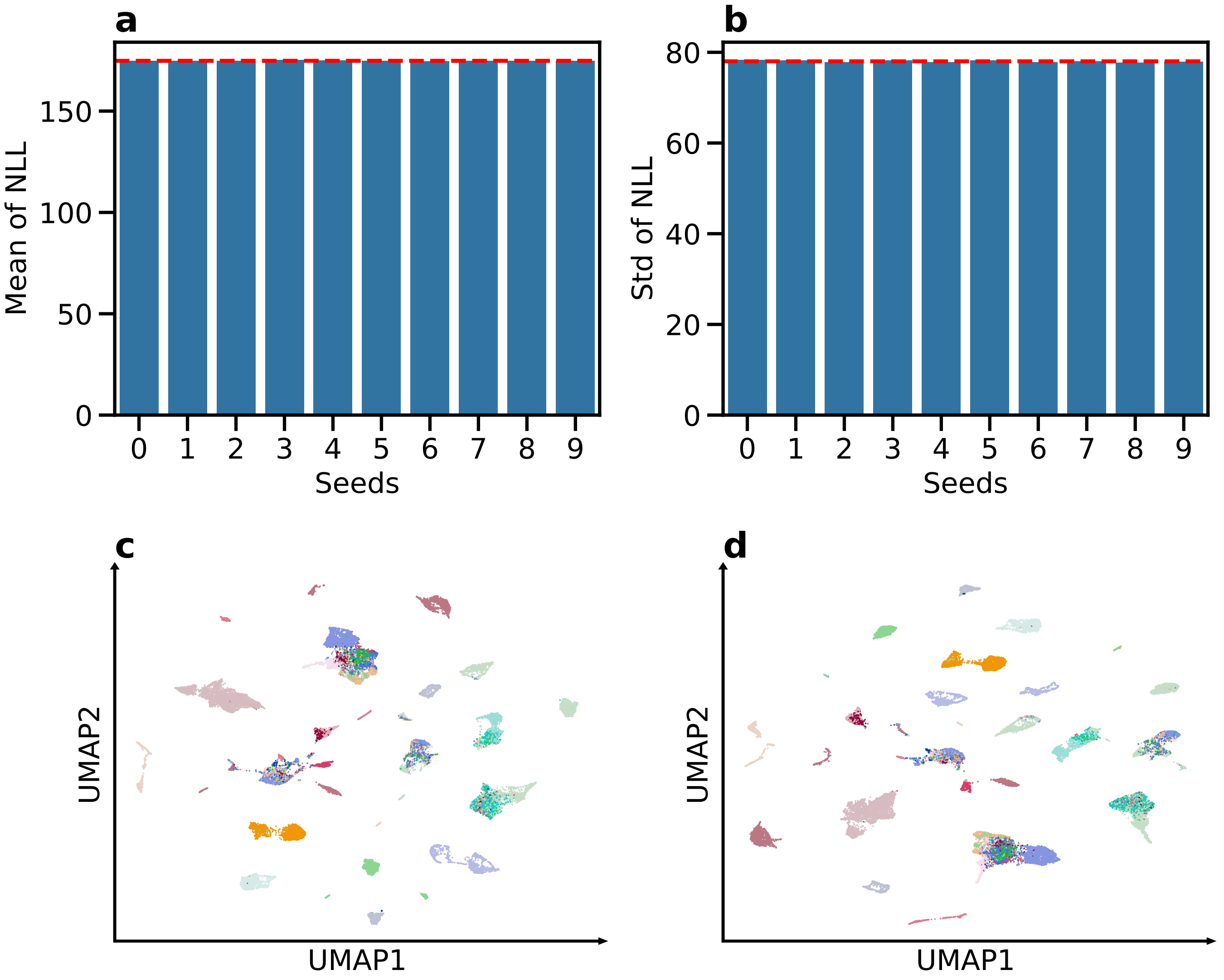

**Supplementary Fig. 6 — tuVI-ZANIDM is robust to random weight initialisation and learns a cell-type-informative latent space. a,b** Mean (**a**) and standard deviation (**b**) of the per-cell negative log-likelihood (NLL) on the held-out *Tabula Muris* test set for tuVI-ZANIDM models trained with ten random seeds. Red dashed lines indicate the corresponding averages across seeds. The small between-seed differences demonstrate that model performance is robust to random weight initialisation. **c,d** UMAPs of the *Tabula Muris* cell embeddings learnt by the models with the lowest (seed 6; **c**) and highest (seed 3; **d**) mean test-set NLL, coloured by annotated cell type. Both embeddings contain distinct, cell-type-associated clusters and show comparable qualitative structure despite differences in their global arrangement. These results indicate that tuVI with a ZANIDM observation model robustly captures cell-type-informative variation in alternative splicing-induced transcript usage.

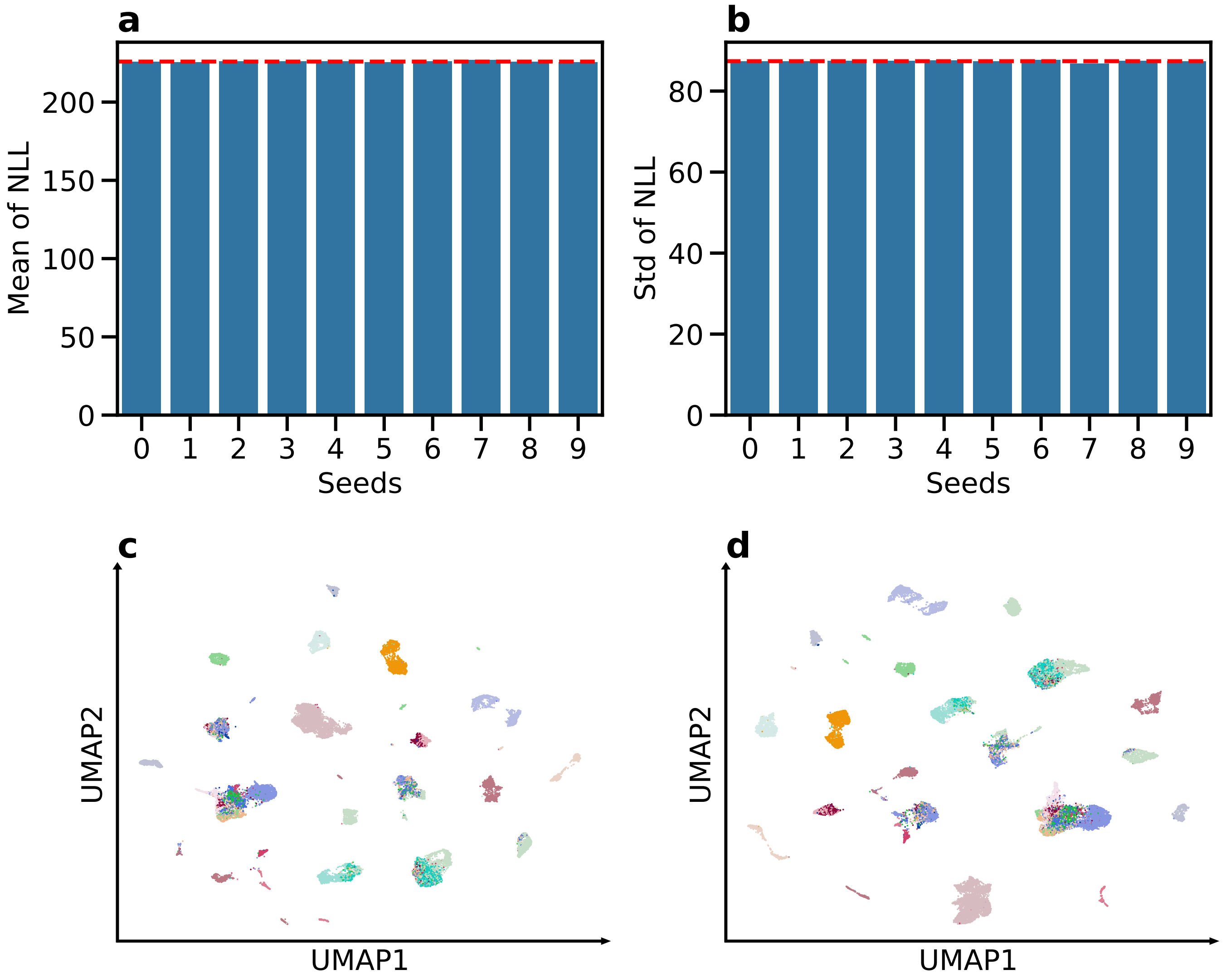

**Supplementary Fig. 7 — tuVI-ZIDM is robust to random weight initialisation and learns a cell-type-informative latent space. a,b** Mean (**a**) and standard deviation (**b**) of the per-cell negative log-likelihood (NLL) on the held-out *Tabula Muris* test set for tuVI-ZIDM models trained with ten random seeds. Red dashed lines indicate the corresponding averages across seeds. The small between-seed differences demonstrate that model performance is robust to random weight initialisation. **c,d** UMAPs of the *Tabula Muris* cell embeddings learnt by the models with the lowest (seed 1; **c**) and highest (seed 7; **d**) mean test-set NLL, coloured by annotated cell type. Both embeddings contain distinct, cell-type-associated clusters and show comparable qualitative structure despite differences in their global arrangement. These results indicate that tuVI with the heuristic ZIDM surrogate robustly captures cell-type-informative variation in alternative splicing-induced transcript usage.

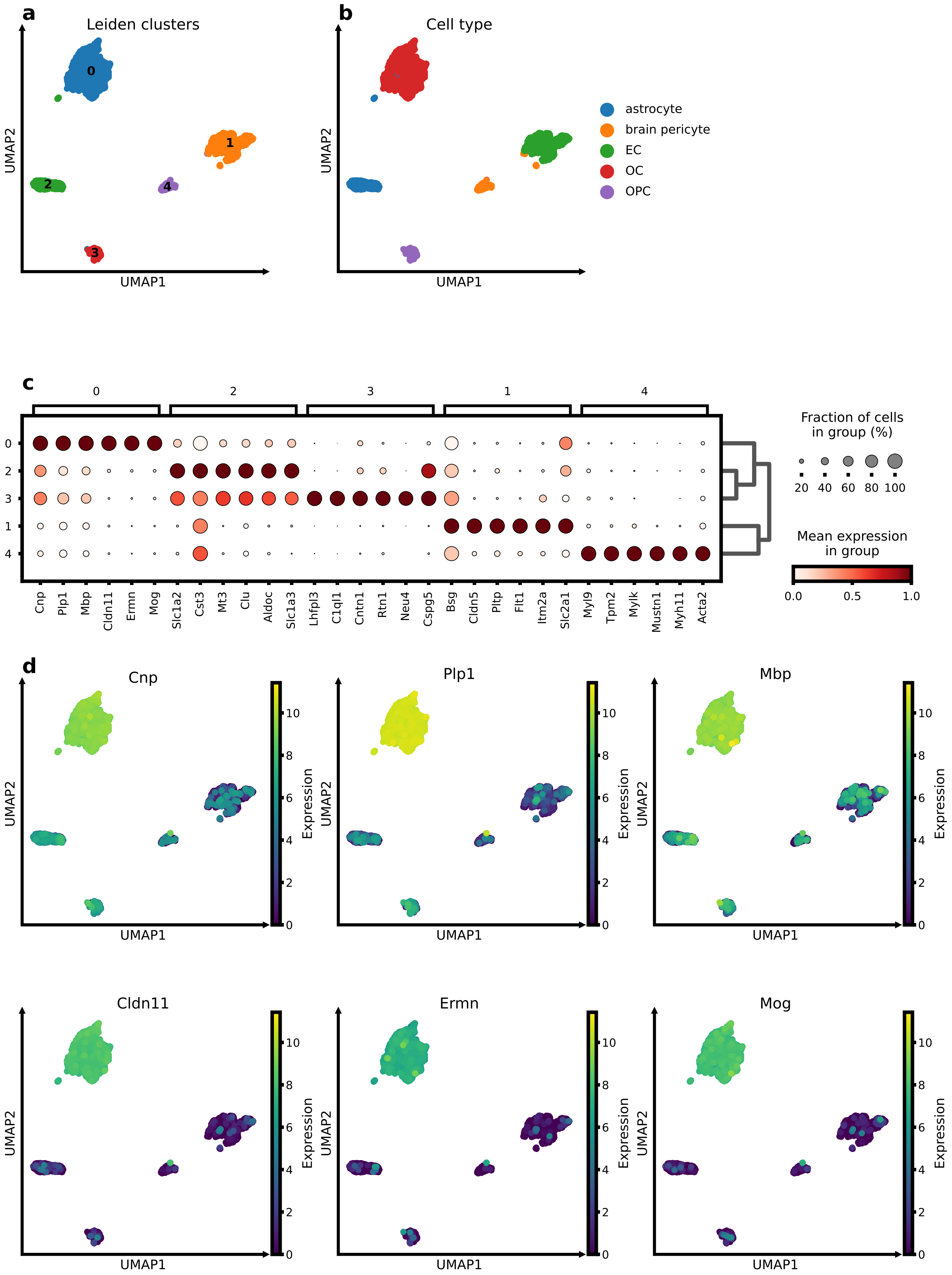
a
Leiden clusters
0
1
2
3
4
UMAP1
UMAP2
b
Cell type
astrocyte
brain pericyte
EC
OC
OPC
UMAP1
UMAP2
c
0
2
3
1
4
Cnp
Plp1
Mbp
Cldn11
Ermn
Mog
Slc1a2
Cst3
Mt3
Clu
Aldoc
Slc1a3
Lhfpl3
C1ql1
Cntn1
Rtn1
Neu4
Cspg5
Bsg
Cldn5
Pltp
Flt1
Itm2a
Slc2a1
Myl9
Tpm2
Mylk
Mustn1
Myh11
Acta2
Fraction of cells in group (%)
20 40 60 80 100
Mean expression in group
0.0 0.5 1.0
d
Cnp
Plp1
Mbp
Cldn11
Ermn
Mog
UMAP1
UMAP2
Expression
0 2 4 6 8 10

**Supplementary Fig. 8 — Cell annotation and differential expression analysis of scVI gene expression cell embeddings in brain non-myeloid tissue of *Tabula Muris*. a,b** UMAP of the scVI embeddings coloured by Leiden cluster (**a**) and *Tabula Muris* reference cell type (**b**), showing close correspondence between the annotations. **c** Mean expression of the top six differentially expressed genes (DEGs) per Leiden cluster identifies cell-type-specific markers. **d** UMAP expression profiles of the top six DEGs for endothelial cells demonstrate their cell-type specificity.

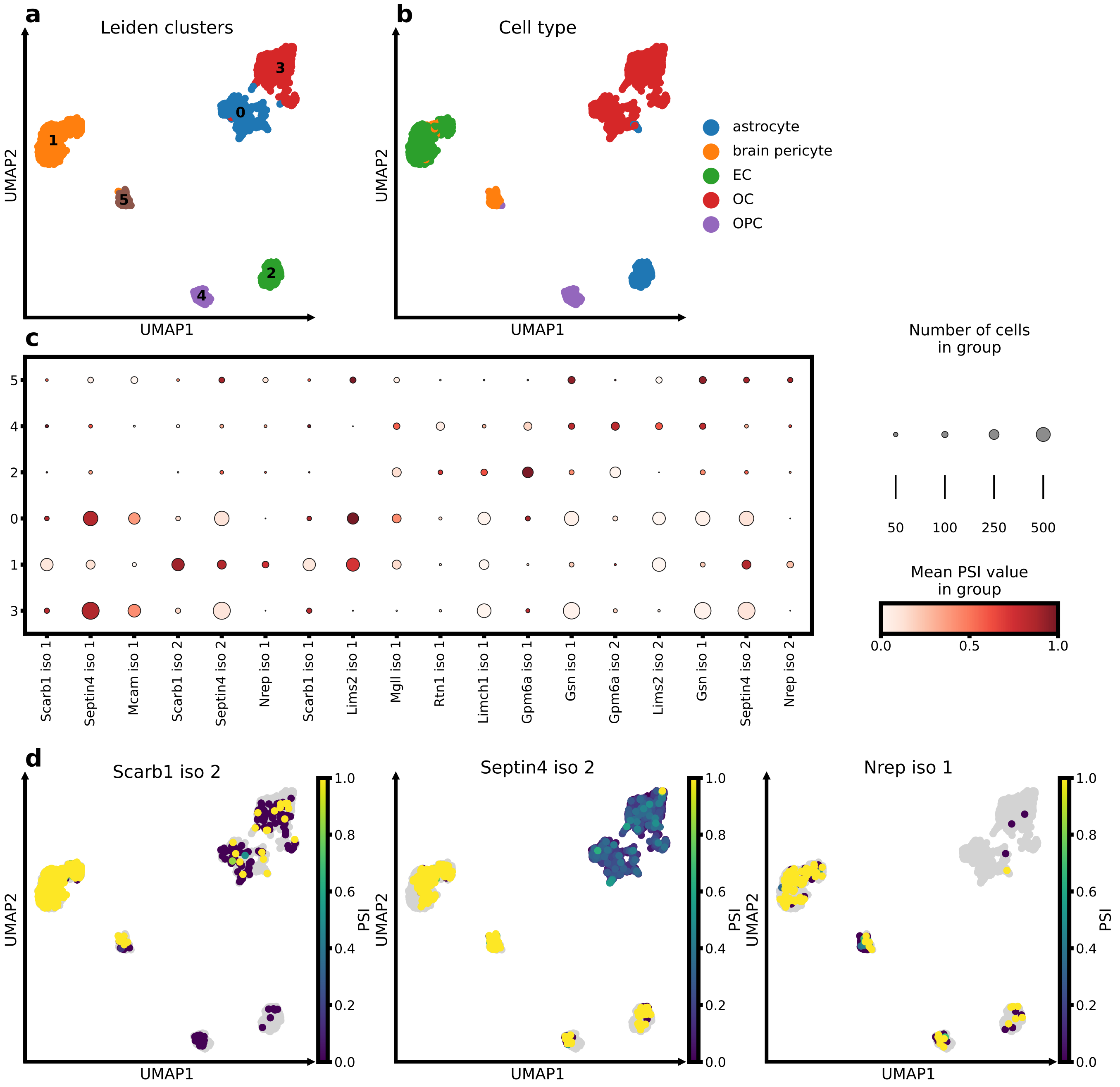

**Supplementary Fig. 9 — Cell annotation and differential splicing analysis of tuVI alternative splicing-induced transcript usage cell embeddings in brain non-myeloid tissue of *Tabula Muris*. a,b** UMAP of the tuVI embeddings coloured by Leiden cluster (**a**) and *Tabula Muris* reference cell type (**b**), showing close correspondence between the annotations but the tuVI cell embeddings also indicate two subpopulations of OCs (Leiden cluster 0 and 3) that were not detected by scVI gene expression embeddings. **c** Mean PSI values of the top three differentially spliced genes (DSG) isoforms per cluster identify candidate cell-type-specific isoform markers. **d** UMAP PSI profiles of the top three DSG isoforms for ECs shows a varied image of cell-type specificity. Jointly this set of isoforms could indicate ECs.

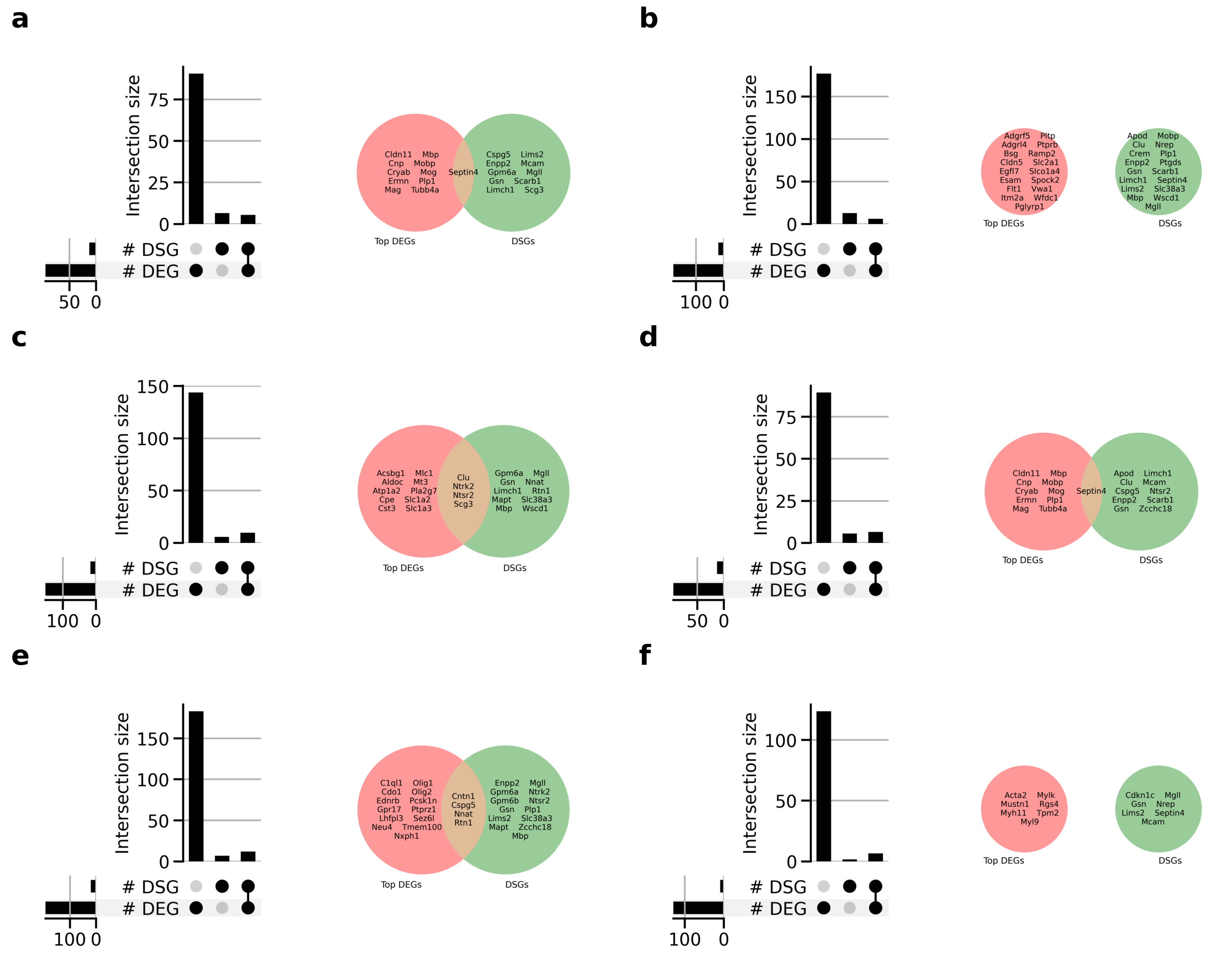

**Supplementary Fig. 10 — Differentially expressed and spliced genes show limited cell-type-specific overlap in brain non-myeloid tissue of *Tabula Muris*.** UpSet plots compare all significant differentially expressed genes (DEGs) and differentially spliced genes (DSGs), whereas Venn diagrams compare all $n$ DSGs with the top $n$ ranked DEGs. **a** OCs (Leiden cluster 0): 95 DEGs and eleven DSGs, with five genes shared; *Septin4* is the only DSG among the top eleven DEGs. **b** ECs: 181 DEGs and 17 DSGs, with five genes shared; no DSG was among the top 17 DEGs. **c** Astrocytes: 152 DEGs and 14 DSGs, with nine genes shared; *Clu*, *Ntrk2*, *Ntsr2*, *Scg3* were among the top 14 DEGs. **d** OCs (Leiden cluster 0): 95 DEGs and eleven DSGs, with five genes shared; *Septin4* is the only DSG among the top eleven DEGs. **e** OPCs: 193 DEGs and 17 DSGs, with eleven shared; *Cntn1*, *Cspg5*, *Nnat*, *Rtn1* were among the top 17 DEGs. **f** Brain pericytes: 128 DEGs and 7 DSGs, with 6 shared; no DSG was among the top six DEGs.

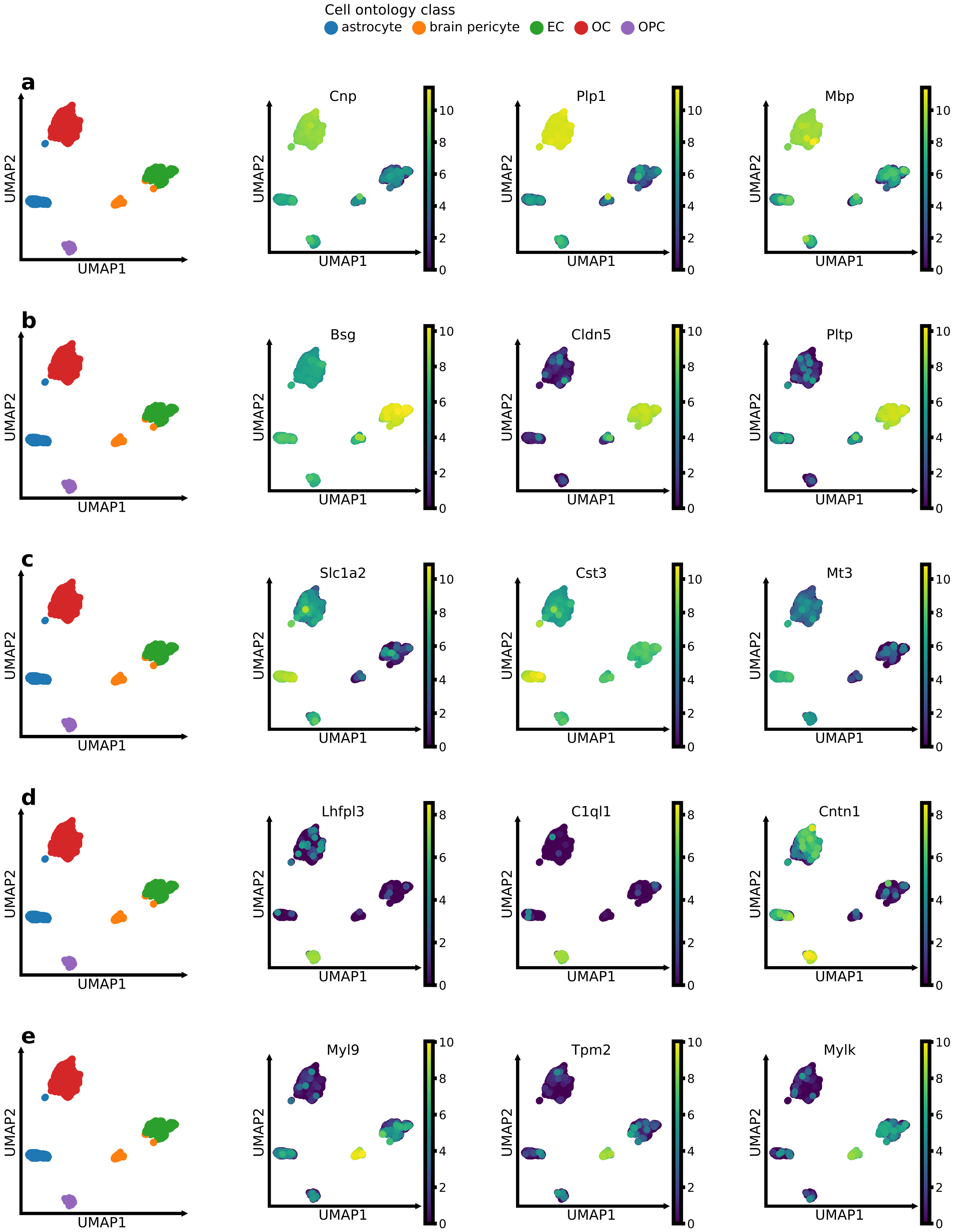

**Supplementary Fig. 11 — Differential analysis of scVI gene expression cell embeddings identifies cell-type-specific marker genes in brain non-myeloid tissue of *Tabula Muris*.** In each row, the first identical UMAP shows the reference cell-type annotation, followed by the expression of the three most significant differentially expressed genes for the indicated Leiden cluster. **a** Cluster 0: OCs (*Cnp*, *Plp1*, *Mbp*). **b** Cluster 1: ECs (*Bsg*, *Cldn5*, *Pltp*). **c** Cluster 2: astrocytes (*Slc1a2*, *Cst3*, *Mt3*). **d** Cluster 3: OPCs (*Lhfpl3*, *C1ql1*, *Cntn1*). **e** Cluster 4: brain pericytes (*Myl9*, *Tpm2*, *Mylk*).

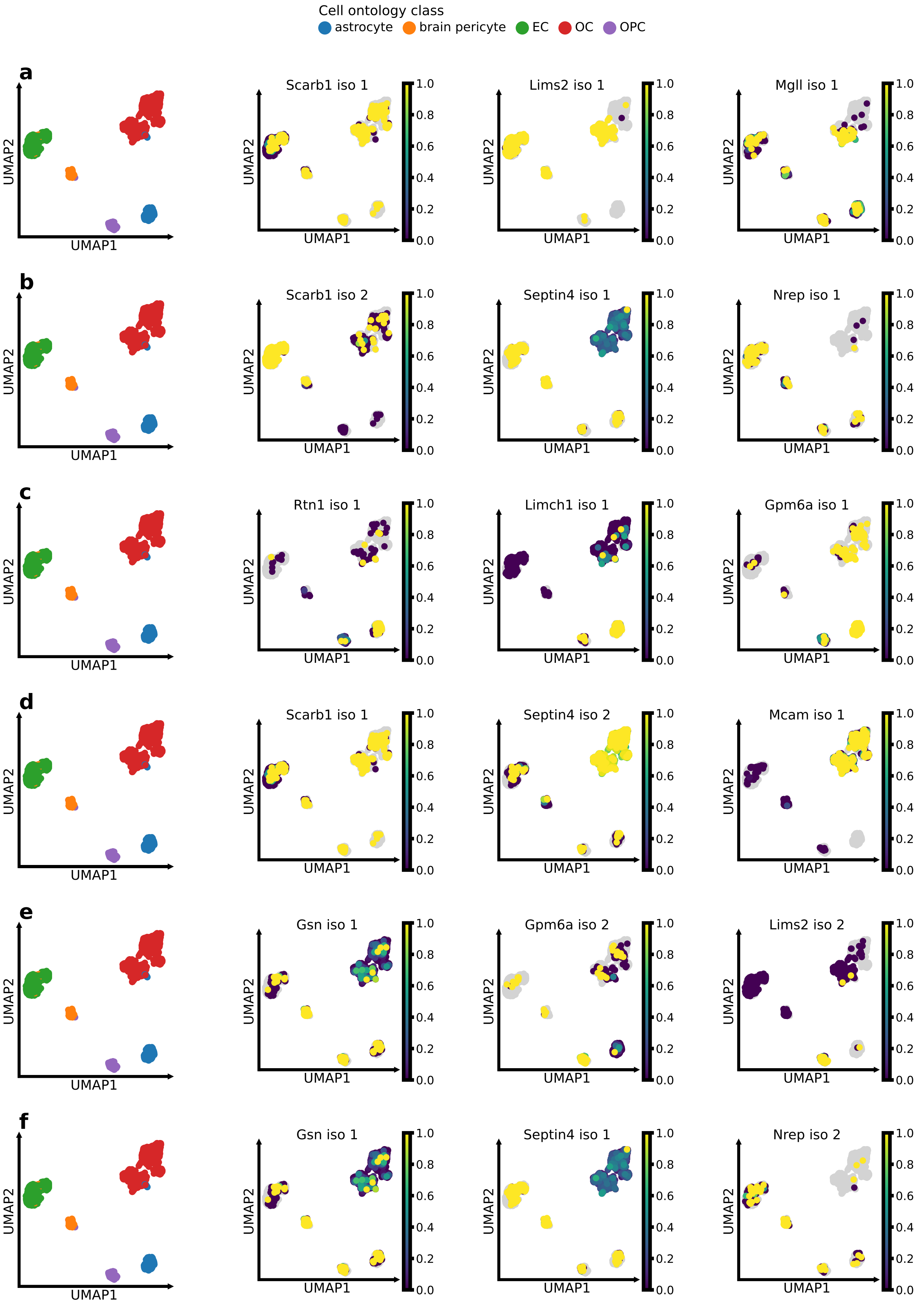
Cell ontology class
astrocyte
brain pericyte
EC
OC
OPC
a
b
c
d
e
f
UMAP1
UMAP2
Scarb1 iso 1
Lims2 iso 1
Mgll iso 1
Scarb1 iso 2
Septin4 iso 1
Nrep iso 1
Rtn1 iso 1
Limch1 iso 1
Gpm6a iso 1
Scarb1 iso 1
Septin4 iso 2
Mcam iso 1
Gsn iso 1
Gpm6a iso 2
Lims2 iso 2
Gsn iso 1
Septin4 iso 1
Nrep iso 2
0.0
0.2
0.4
0.6
0.8
1.0

**Supplementary Fig. 12 — Differential analysis of tuVI alternative splicing-induced transcript usage embeddings identifies cell-type-specific isoforms as potential markers in brain non-myeloid tissue of *Tabula Muris*.** In each row, the first UMAP shows the *Tabula Muris* reference cell-type annotation, followed by the PSI score of the three most significant differentially spliced gene isoforms for the indicated Leiden cluster. **a** Cluster 0: OCs (*Scarb* iso 1, *Lims2* iso 1, *Mgll* iso 1). **b** Cluster 1: ECs (*Scarb* iso 2, *Septin4* iso 1, *Nrep* iso 1). **c** Cluster 2: astrocytes (*Rtn* iso 1, *Limch1* iso 1, *Gpm6a* iso 1). **d** Cluster 3: OCs (*Scarb* iso 1, *Septin4* iso2, *Mcam* iso 1). **e** Cluster 4: OPCs (*Gsn* iso 1, *Gpm6a* iso 2, *Lims2* iso 2). **f** Cluster 5: brain pericytes (*Gsn* iso 1, *Septin4* iso 1, *Nrep* iso 2)

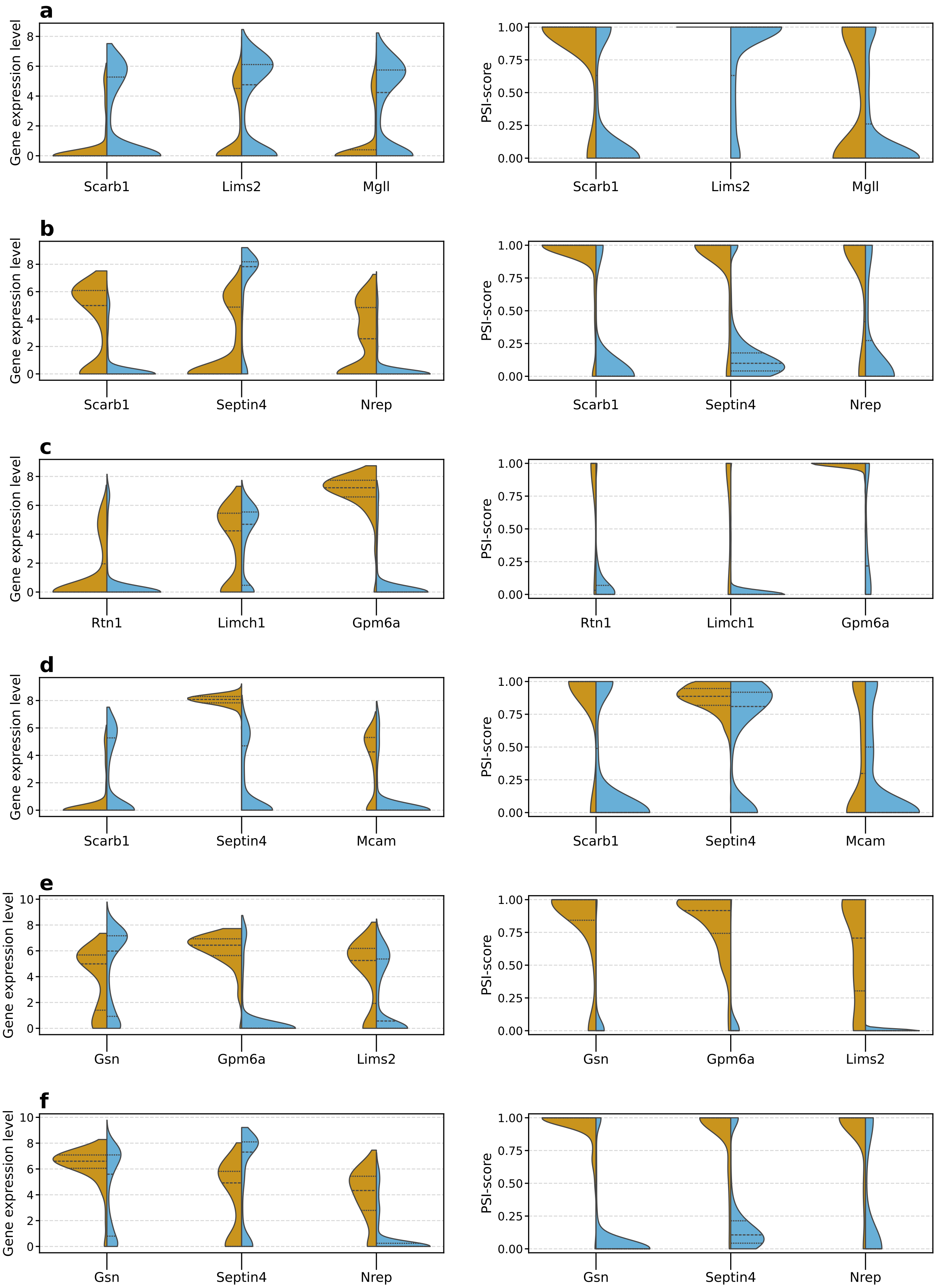
a
Gene expression level
Scarb1
Lims2
Mgll
PSI-score
Scarb1
Lims2
Mgll
b
Gene expression level
Scarb1
Septin4
Nrep
PSI-score
Scarb1
Septin4
Nrep
c
Gene expression level
Rtn1
Limch1
Gpm6a
PSI-score
Rtn1
Limch1
Gpm6a
d
Gene expression level
Scarb1
Septin4
Mcam
PSI-score
Scarb1
Septin4
Mcam
e
Gene expression level
Gsn
Gpm6a
Lims2
PSI-score
Gsn
Gpm6a
Lims2
f
Gene expression level
Gsn
Septin4
Nrep
PSI-score
Gsn
Septin4
Nrep

**Supplementary Fig. 13 — PSI distributions reveal cell-type-specific isoform markers beyond gene expression in brain non-myeloid tissue of *Tabula Muris*.** Left violins compare gene expression in the indicated cell type with all other cells; right violins compare the corresponding isoform PSI distributions. **a** In OCs (cluster 0), *Scarb1*, *Lims2*, and *Mgll* are not upregulated, but their isoforms show OC-specific usage. Enrichment of the *Lims2* and *Mgll* isoforms in cluster 0 relative to OC cluster 3 suggests an OC subpopulation. **b** In ECs, *Scarb1* and *Nrep*, but not *Septin4*, are upregulated; all three isoforms show greater EC specificity than gene expression. **c** In astrocytes, *Gpm6a*, but not *Rtn1* or *Limch1*, is upregulated, whereas all three isoforms are cell-type-specific. **d** In OCs (cluster 3), *Septin4* and *Mcam*, but not *Scarb1*, are upregulated. The *Scarb1* and *Septin4* isoforms are OC-specific, whereas the *Mcam* isoform is not; *Septin4* also marks OC cluster 0. **e** In OPCs, *Gpm6a* and *Lims2*, but not *Gsn*, are upregulated. The *Gsn* and *Gpm6a* isoforms are OPC-specific, whereas *Lims2* is not. **f** In brain pericytes, *Gsn* and *Nrep*, but not *Septin4*, show elevated expression. Their isoforms show cell-type-specific PSI distributions, although sparse splicing measurements warrant caution.

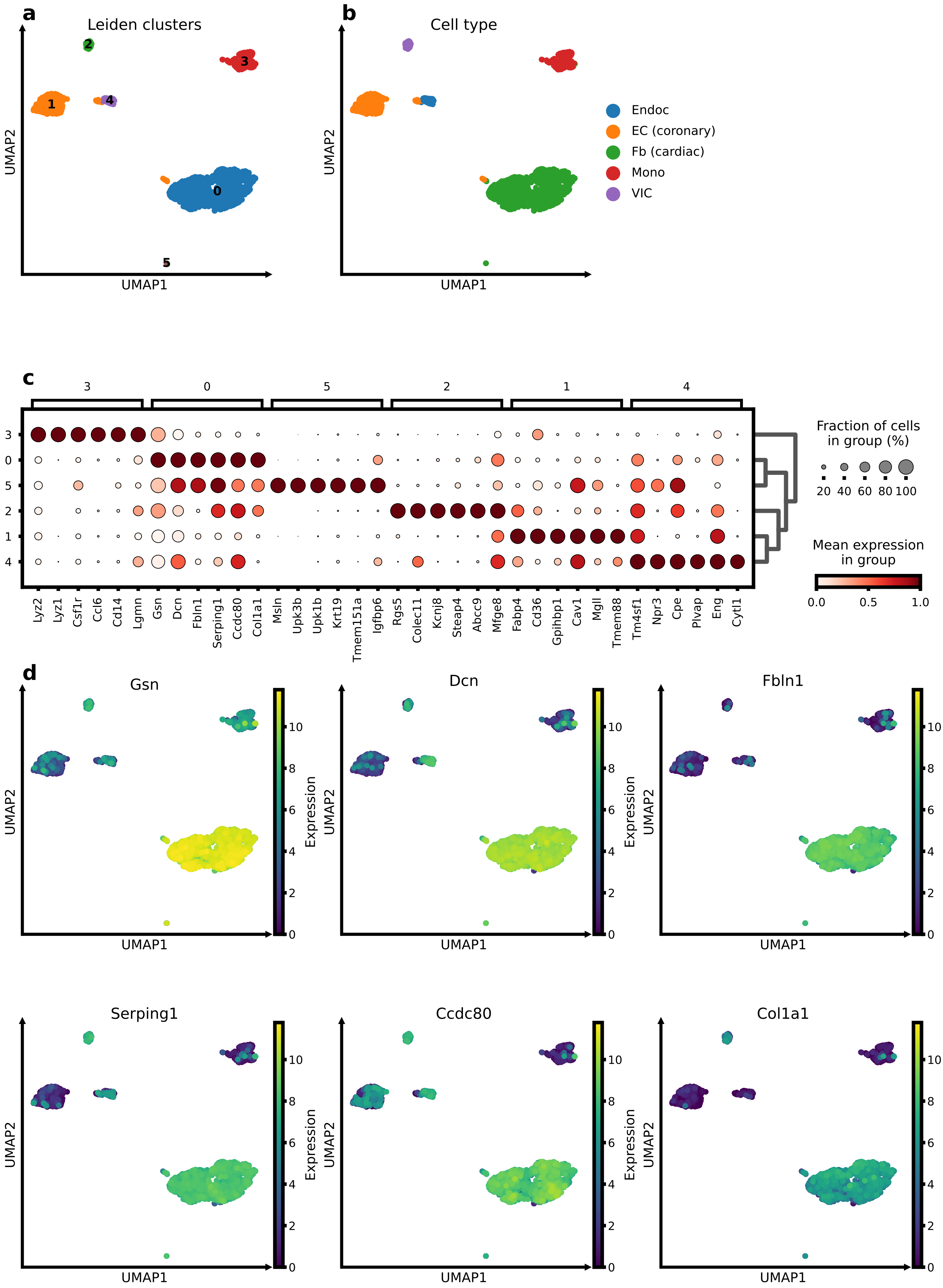
a
Leiden clusters
0
1
2
3
4
5
UMAP1
UMAP2
b
Cell type
Endoc
EC (coronary)
Fb (cardiac)
Mono
VIC
UMAP1
UMAP2
c
3
0
5
2
1
4
Lyz2
Lyz1
Csf1r
Ccl6
Cd14
Lgmn
Gsn
Dcn
Fbln1
Serping1
Ccdc80
Col1a1
Msln
Upk3b
Upk1b
Krt19
Tmem151a
Igfbp6
Rgs5
Colec11
Kcnj8
Steap4
Abcc9
Mfge8
Fabp4
Cd36
Gpihbp1
Cav1
Mgll
Tmem88
Tm4sf1
Npr3
Cpe
Plvap
Eng
Cytl1
Fraction of cells in group (%)
20 40 60 80 100
Mean expression in group
0.0 0.5 1.0
d
Gsn
Dcn
Fbln1
Serping1
Ccdc80
Col1a1
UMAP1
UMAP2
Expression
0 2 4 6 8 10

**Supplementary Fig. 14 — Cell annotation and differential gene expression analysis scVI gene expression cell embeddings in heart tissue of *Tabula Muris*.** **a,b** UMAP of the scVI embeddings coloured by Leiden cluster (**a**) and *Tabula Muris* reference cell type (**b**), showing close correspondence between the annotations. **c** Mean expression of the top six differentially expressed genes (DEGs) per Leiden cluster identifies cell-type-specific markers. **d** UMAP expression profiles of the top six DEGs for cardiac fibroblasts (Fb (cardiac) )demonstrate their cell-type specificity.

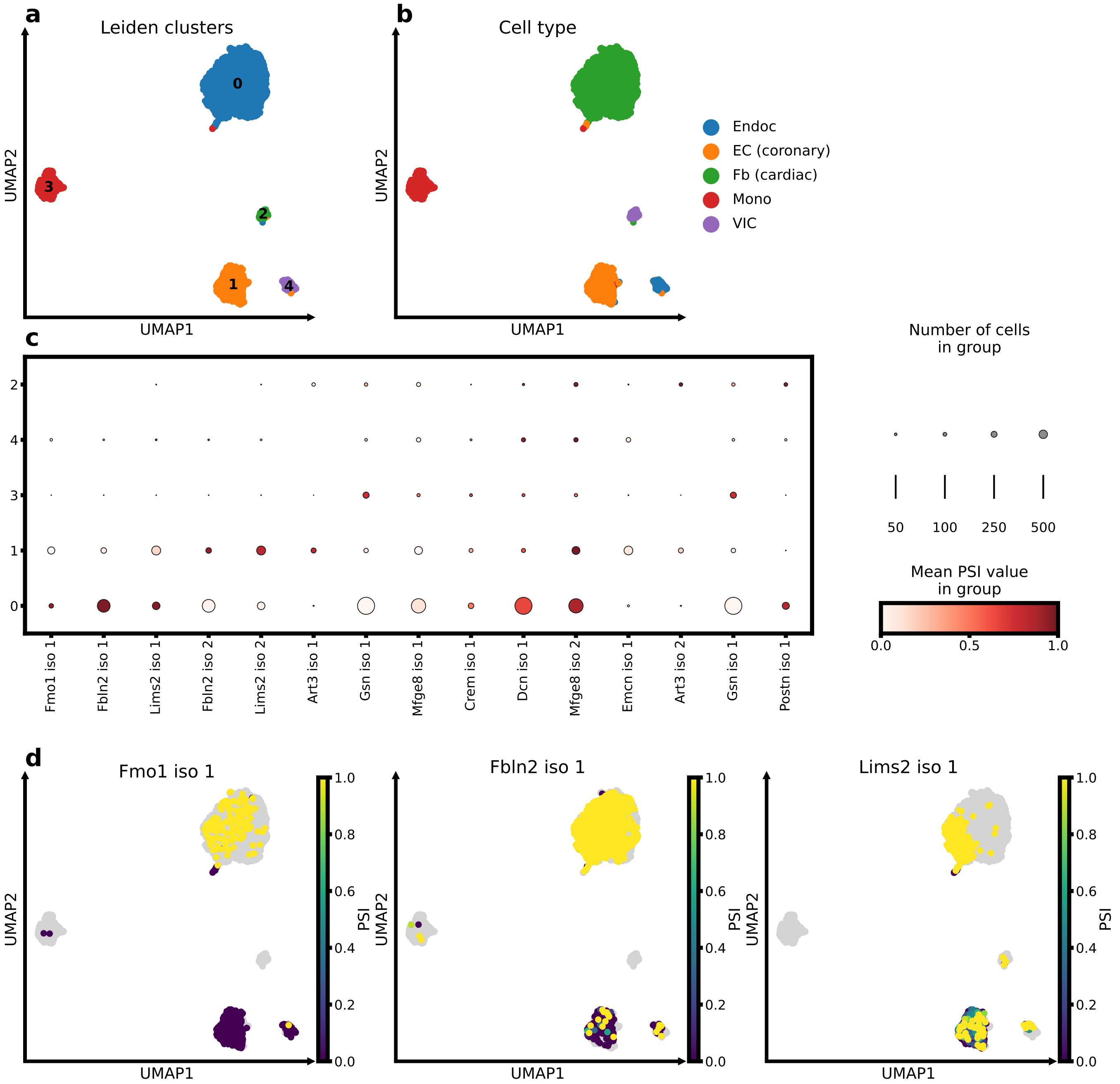

**Supplementary Fig. 15 — Cell annotation and differential splicing analysis of tuVI alternative splicing-induced transcript usage cell embeddings in heart tissue of *Tabula Muris*. a,b** UMAP of the tuVI embeddings coloured by Leiden cluster (**a**) and *Tabula Muris* reference cell type (**b**), showing close correspondence between the annotations. **c** Mean PSI values of the top three differentially spliced gene (DSG) isoforms per cluster identify candidate cell-type-specific isoform markers. **d** UMAP PSI profiles of the top three DSG isoforms for cardiac Fbs demonstrate their cell-type specificity.

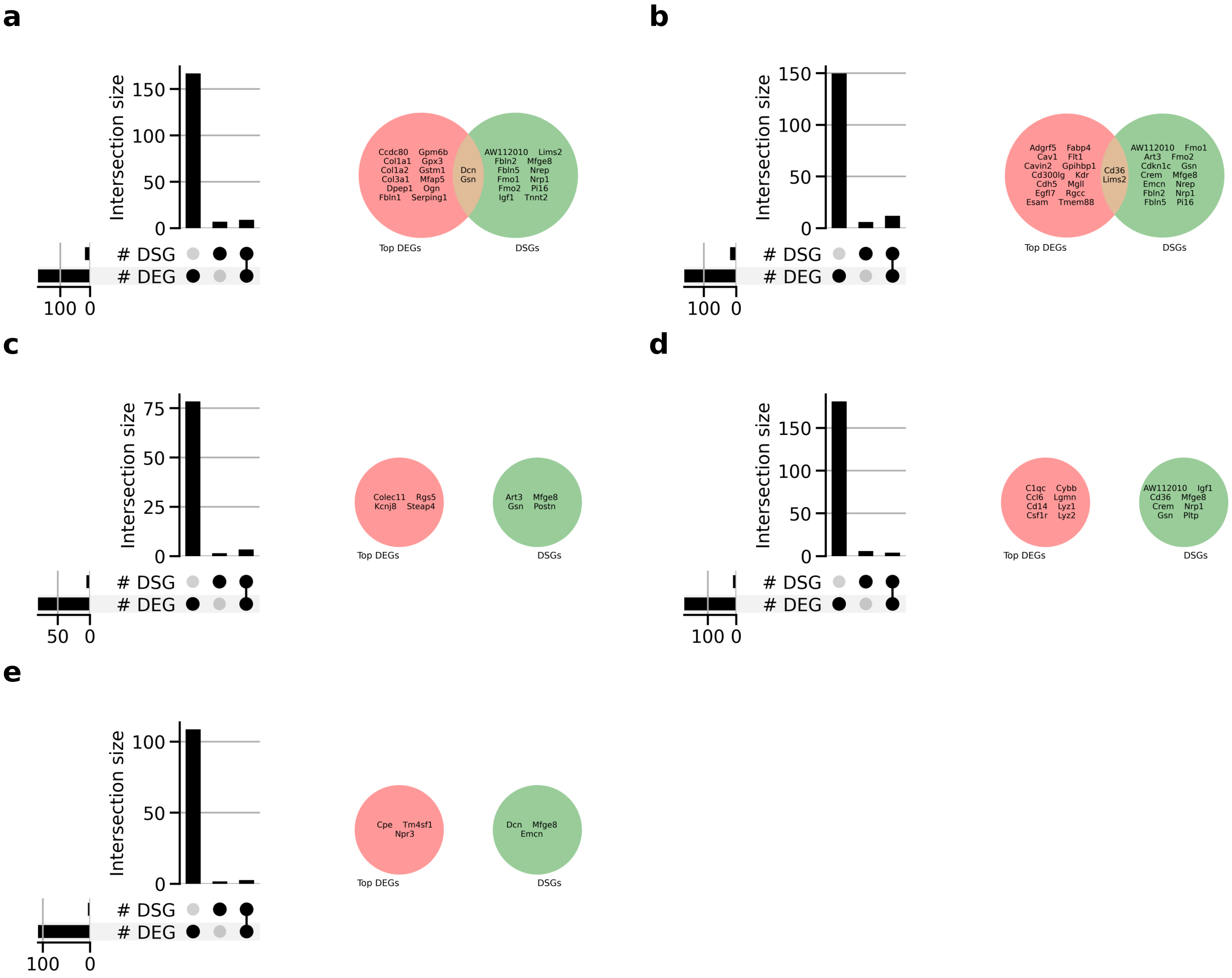

**Supplementary Fig. 16 — Differentially expressed and spliced genes show limited cell-type-specific overlap in heart tissue of *Tabula Muris*.** UpSet plots compare all significant differentially expressed genes (DEGs) and differentially spliced genes (DSGs), whereas Venn diagrams compare all $n$ DSGs with the top $n$ ranked DEGs. **a** Cardiac Fbs: 174 DEGs and 14 DSGs, with eight genes shared; *Dcn* and *Gsn* were among the top 14 DEGs. **b** Coronary ECs: 160 DEGs and 16 DSGs, with 11 genes shared; *Cd36* and *Lims2* were among the top 16 DEGs. **c** VICs: 81 DEGs and four DSGs, with three genes shared; no DSG was among the top four DEGs. **d** Monos: 183 DEGs and eight DSGs, with three genes shared; no DSG was among the top eight DEGs. **e** Endocs: 110 DEGs and three DSGs, with two genes shared; no DSG was among the top three DEGs.

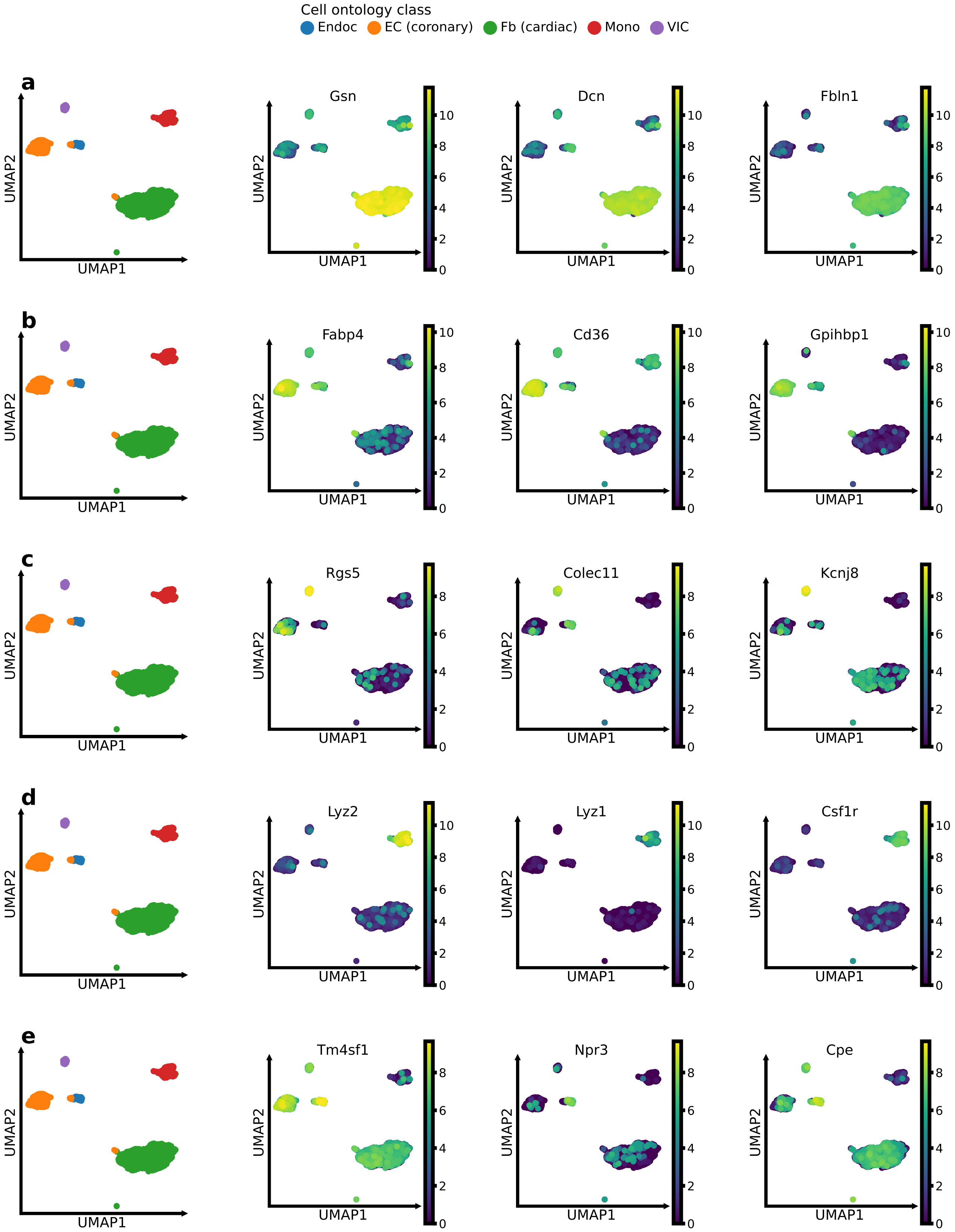

**Supplementary Fig. 17 — Differential analysis of scVI gene expression cell embeddings identifies cell-type-specific marker genes in heart tissue of *Tabula Muris*.** In each row, the first UMAP shows the *Tabula Muris* reference cell-type annotation, followed by the expression of the three most significant differentially expressed genes for the indicated

Leiden cluster. **a** Cluster 0: cardiac Fbs (*Gsn*, *Dcn*, *Fbln1*). **b** Cluster 1: coronary ECs (*Fabp4*, *Cd36*, *Gpihbp1*). **c** Cluster 2: VICs (*Rgs5*, *Colec11*, *Kcnj8*). **d** Cluster 3: Monos (*Lyz2*, *Lyz1*, *Csf1r*). **e** Cluster 4: Endocs (*Tm4sf1*, *Npr3*, *Cpe*).

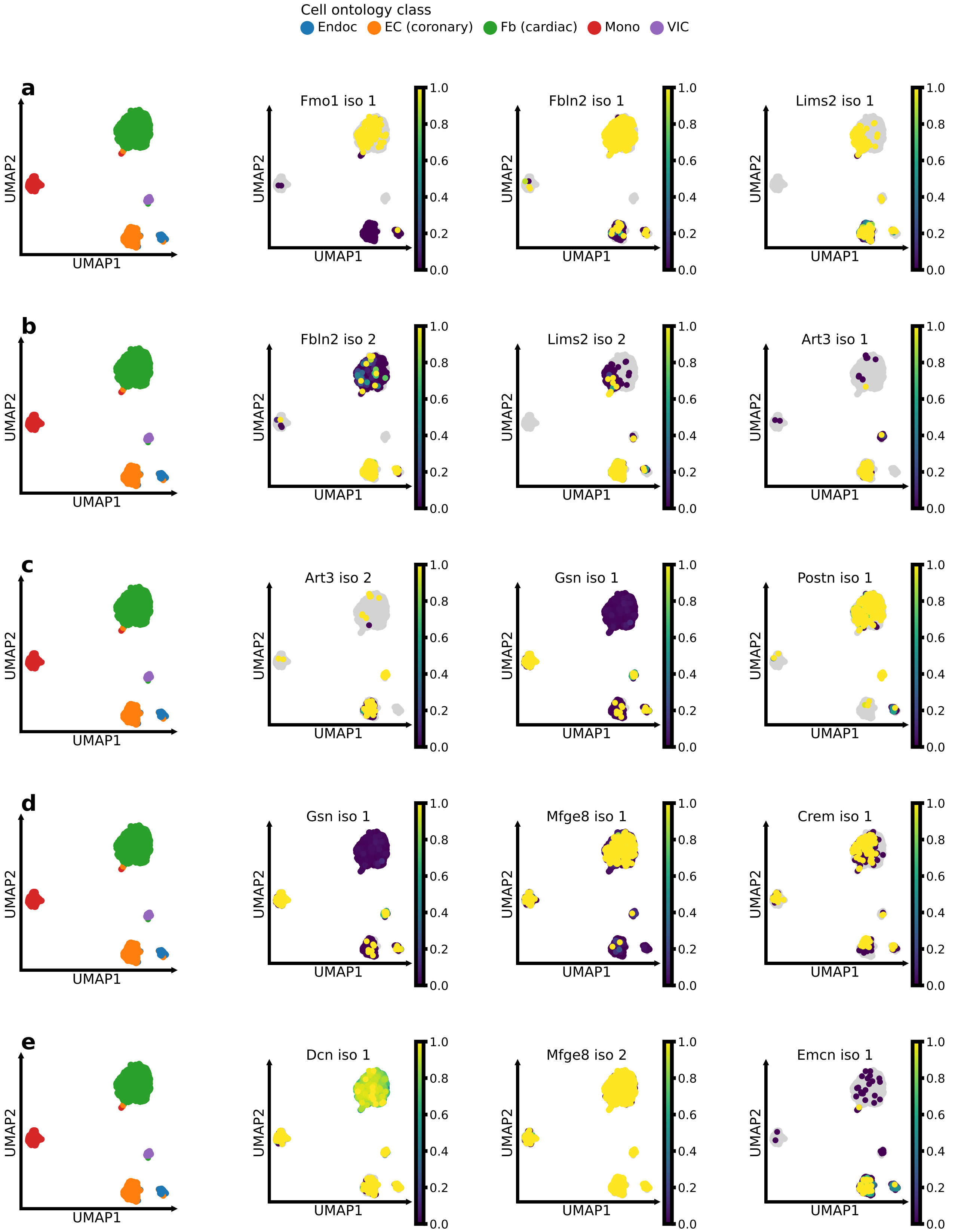

**Supplementary Fig. 18 — Differential analysis of tuVI AS-induced transcript usage embeddings identifies cell-type-specific isoforms as potential markers in heart tissue of *Tabula Muris*.** In each row, the first UMAP shows the *Tabula Muris* reference cell-type annotation, followed by the PSI score of the three most significant differentially spliced gene isoforms for the indicated Leiden cluster. **a** Cluster 0: cardiac Fbs (*Fmo1* iso 1, *Fbln2* iso 1, *Lims2* iso 1). **b** Cluster 1: coronary

ECs (*Fbln2* iso 2, *Lims2* iso 2, *Art3* iso 1). **c** Cluster 2: VICs (*Art3* iso 2, *Gsn* iso 1, *Postn* iso 1). **d** Cluster 3: Monos (*Gsn* iso 1, *Mfge8* iso 1, *Crem* iso 1). **e** Cluster 4: Endocs (*Dcn* iso 1, *Mfge8* iso 2, *Emcn* iso 1).

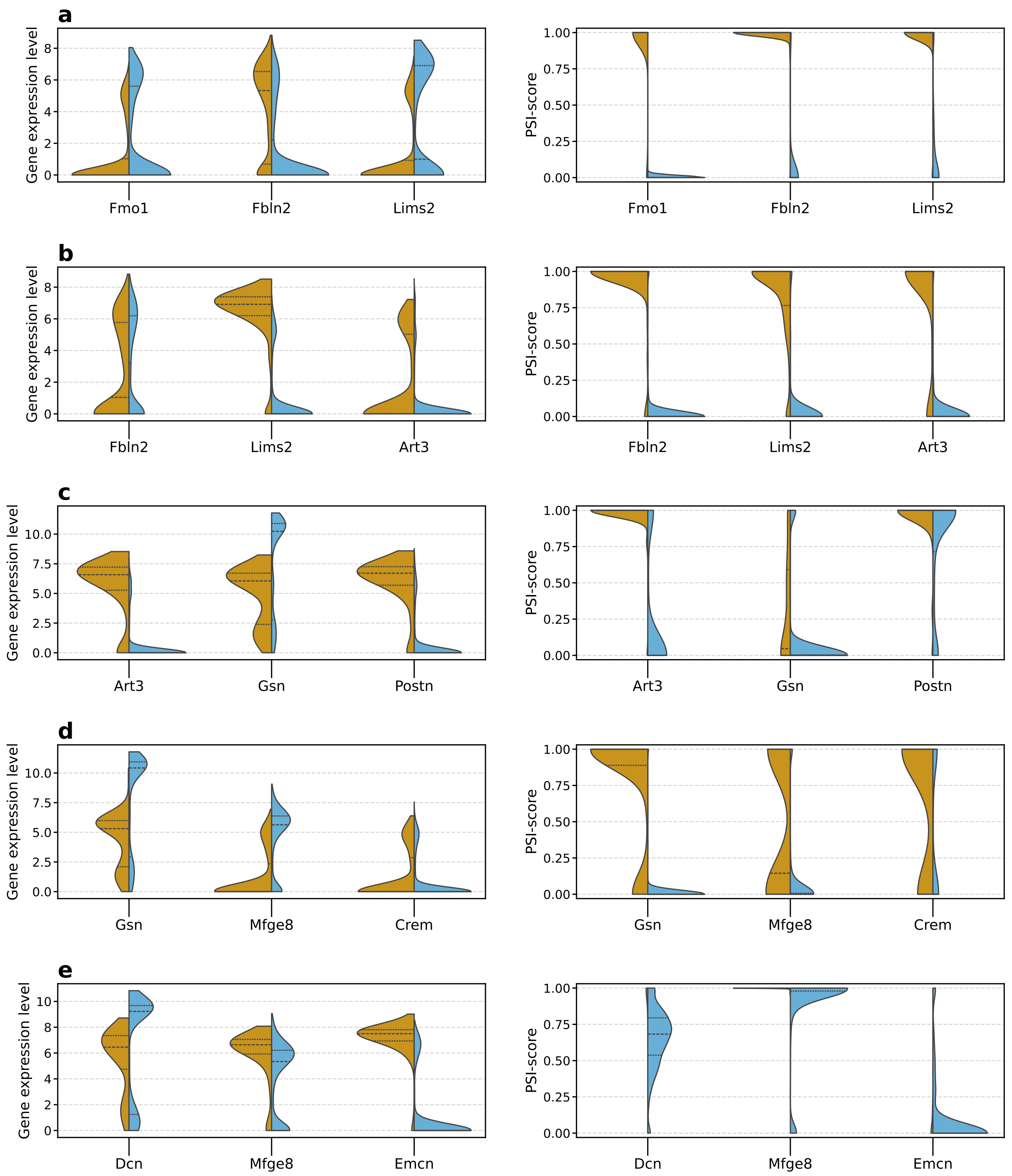

**Supplementary Fig. 19 — PSI distributions identify cell-type-specific isoform markers more clearly than gene expression levels in heart tissue of *Tabula Muris.*** In each row, the left violin plots compare gene expression between the indicated cell type and all other cells, while the right plots compare the corresponding isoform PSI distributions. **a** In cardiac Fbs, *Fmo1*, *Fbln2*, and *Lims2* show no expression level upregulation but highly cell-type-specific PSI distributions. **b** In coronary ECs, *Lims2* and *Art3*, but not *Fbln2*, are upregulated, whereas all three isoforms show cell-type-specific PSI distributions. **c** In VICs, *Art3* and *Postn* show elevated expression and cell-type-specific PSI distributions, whereas *Gsn* remains inconclusive. **d** In Monos, none of the three genes is upregulated; however, the *Gsn* isoform is cell-type-specific, while *Mfge8* and *Crem* are ambiguous. **e** In Endocs, *Mfge8* and *Emcn* are upregulated, but their isoforms are not cell-type-specific, potentially reflecting limited alternative splicing-induced transcript usage observations in this sparsely represented cell type.

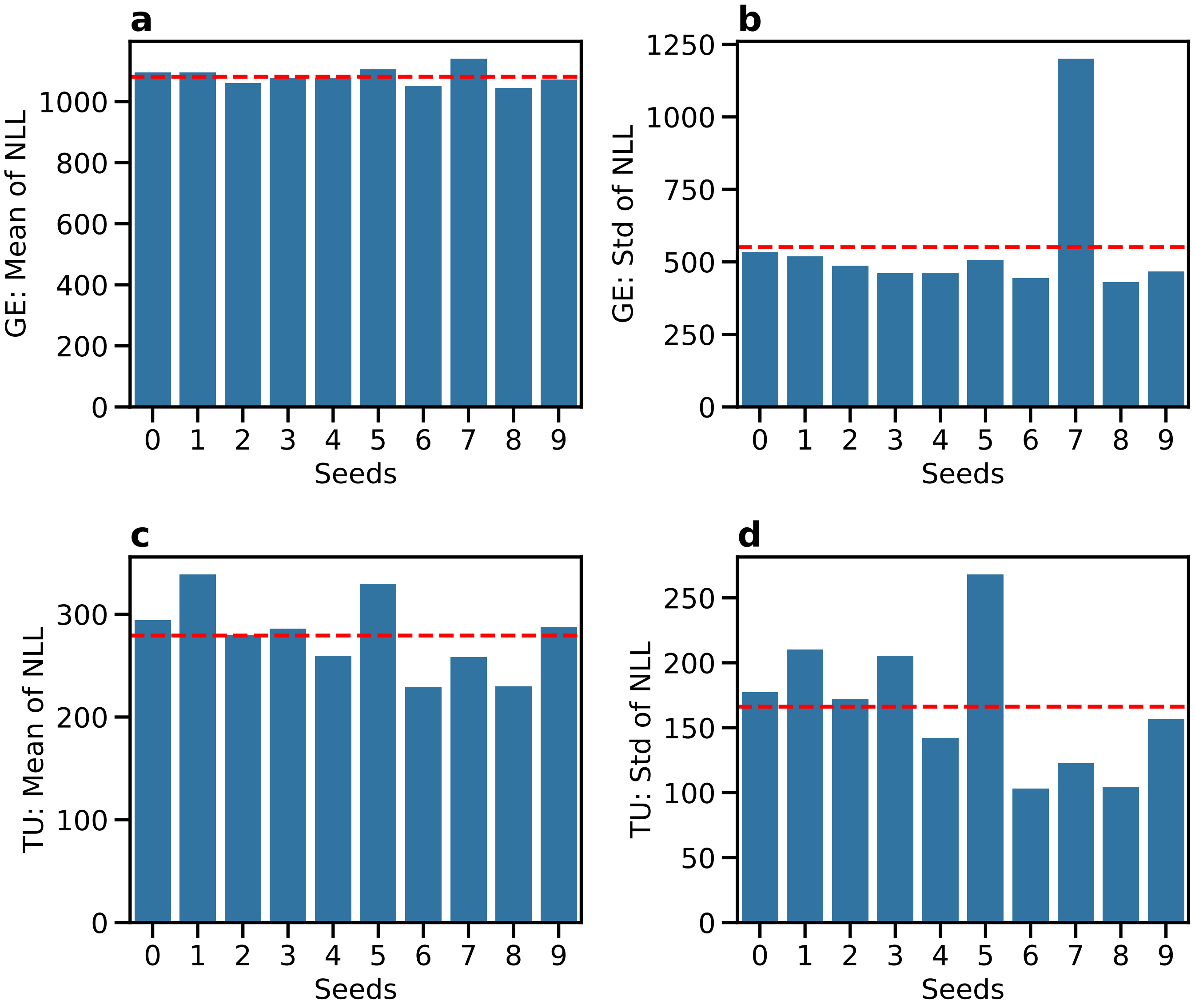

**Supplementary Fig. 20 — Model selection and robustness of TRVI to random weight initialisation benchmarked on *Tabula Muris*. a** Mean per-cell negative log-likelihood (NLL) for the gene expression (GE) modality on the held-out test set for TRVI-ZINB-ZIDM models trained using ten random seeds. **b** Standard deviation of the per-cell GE NLL on the held-out test set. Seed 7 shows substantially greater dispersion than the other seeds. **c** Mean per-cell NLL for the alternative-splicing-induced transcript usage (TU) modality on the held-out test set. **d** Standard deviation of the per-cell TU NLL on the held-out test set. Seed 5 shows the greatest dispersion. In all panels, red dashed lines indicate the corresponding mean across seeds.

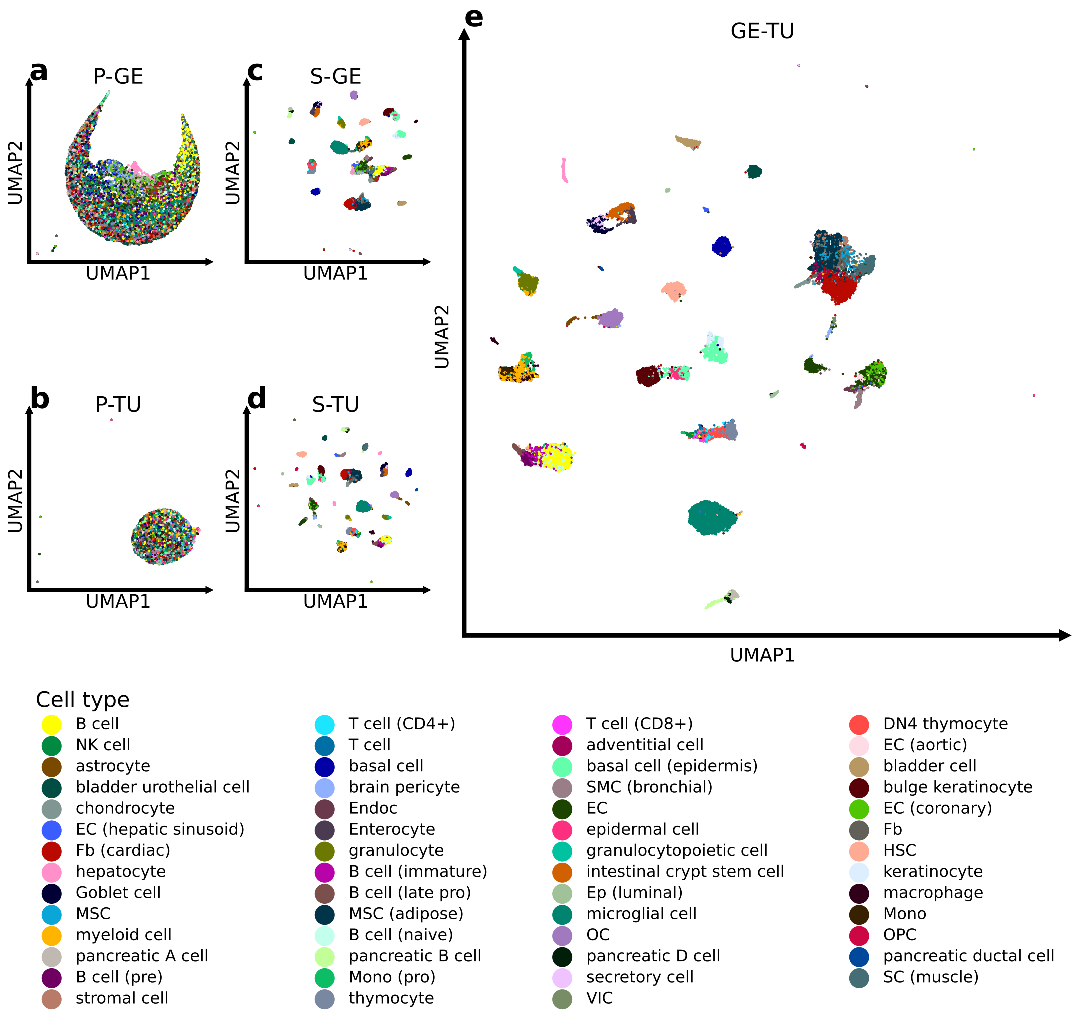

**Supplementary Fig. 21 — TRVI concentrates cell-type-associated structure in its joint gene expression and transcript usage latent representations.** The TRVI model (seed 8) trained on the *Tabula Muris* atlas was used to infer cell embeddings from single-cell gene expression (GE) and alternative splicing-induced transcript usage (TU). **a**–**e** UMAP projections of the private GE (P-GE; **a**), private TU (P-TU; **b**), shared GE (S-GE; **c**), shared TU (S-TU; **d**) and joint GE–TU (GE-TU; **e**) latent cell embeddings. Each point represents a cell and is coloured by its annotated cell type. The private representations show limited separation by cell type, consistent with their role in capturing modality-specific variation. By contrast, the S-GE, S-TU and joint GE–TU representations form discrete clusters aligned with the cell-type annotations, indicating that cell-type-associated information is concentrated in the shared and integrated representations.

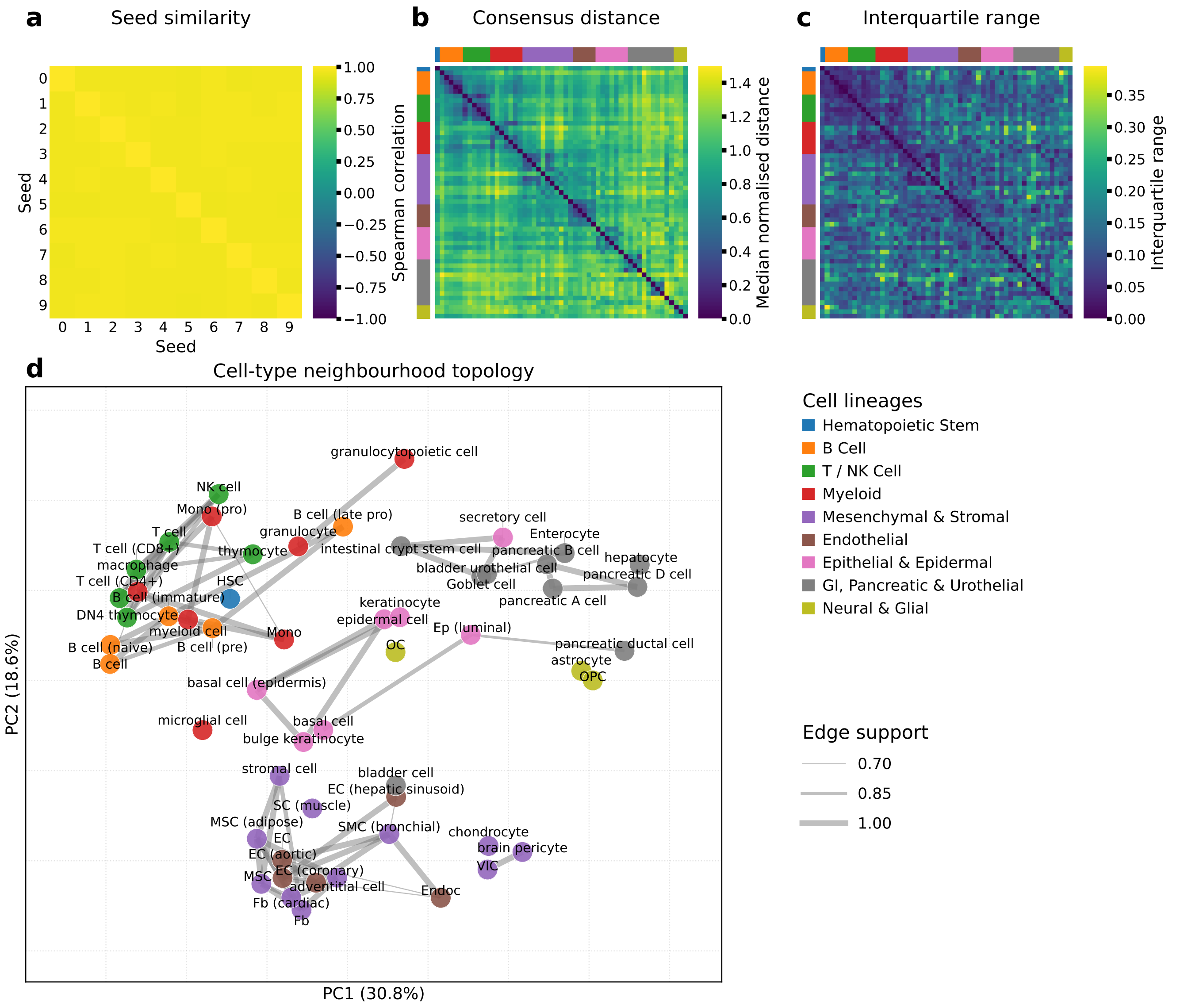

**Supplementary Fig. 22 — The geometry of the joint gene expression–transcript usage latent space of TRVI is stable across random weight initialisations. a** Global geometric stability was assessed by comparing Euclidean distance matrices computed from cell type centroids in the joint gene expression–transcript usage (GE–TU) embeddings across ten random seeds (0–9) learnt on *Tabula Muris*. Spearman correlations between the vectorised pairwise distances were consistently high (median across unique off-diagonal seed pairs, $\rho = 0.958$; range, 0.949–0.968). **b** The consensus distance between each pair of the 55 cell-type centroids was defined as the median of the normalised distances across the ten seeds. Cell types are ordered by broad lineage. The resulting consensus matrix reveals lineage-associated blocks of short distances along the diagonal, indicating that related cell types consistently occupy nearby regions of the latent space. **c** The interquartile range (25th–75th percentiles) of each pairwise normalised distance across seeds quantifies local variability. The predominantly low interquartile ranges indicate that most pairwise relationships are reproducible across initialisations. **d** Stability of the local cell-type topology was assessed by constructing a five-nearest-neighbour graph for each seed. Nodes represent cell types and are coloured by broad lineage; edge support denotes the fraction of seeds in which a given neighbourhood relationship was recovered. Only edges supported by at least seven of the ten seeds are shown, with line width indicating support from 0.70 (7/10 seeds) to 1.00 (10/10 seeds). The resulting stable neighbourhood structure forms distinct groups that broadly recapitulate cell-lineage organisation showcasing the preserved cell type geometry of the latent space across differing seeds.

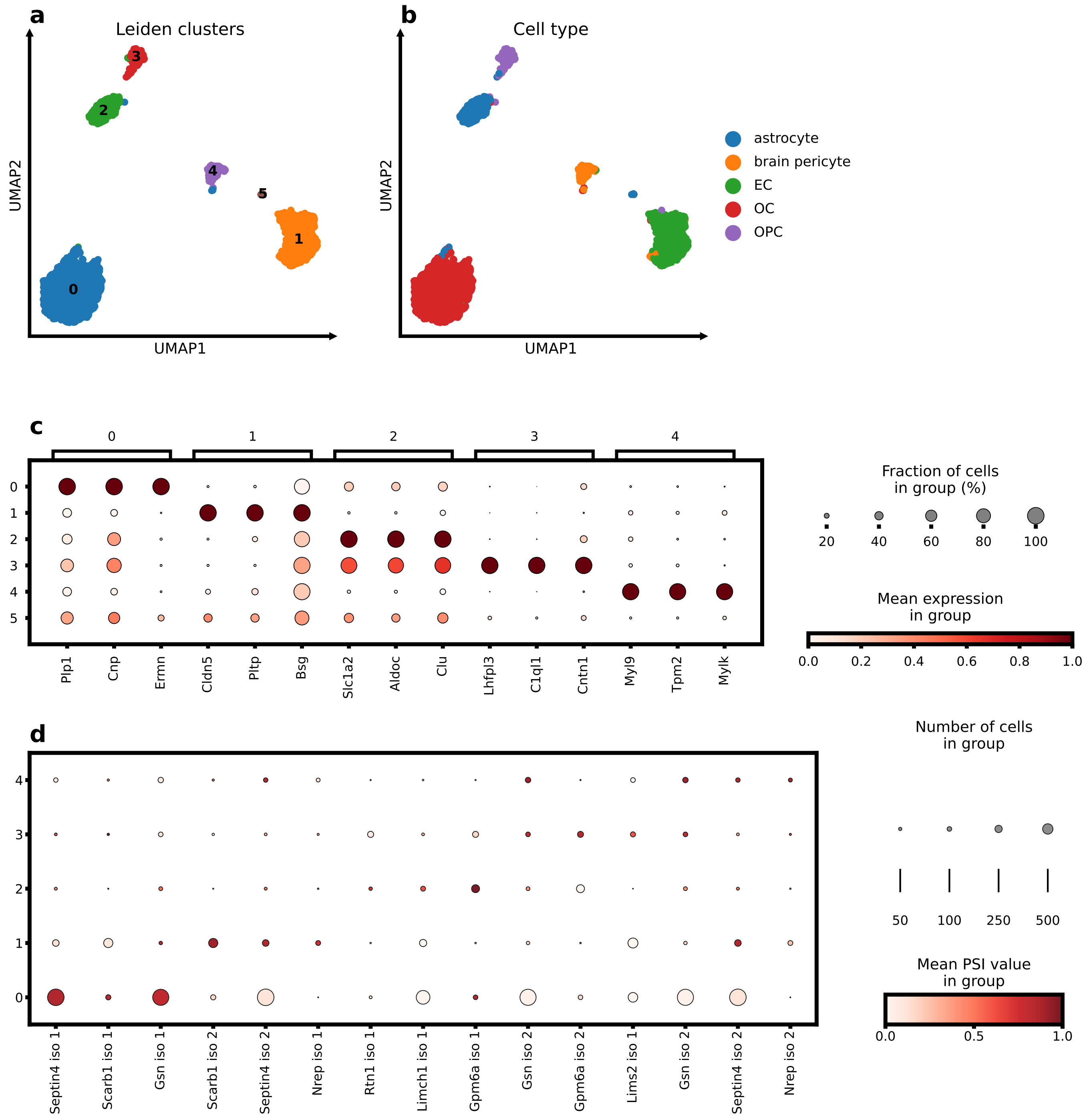

**Supplementary Fig. 23 — TRVI's GE-TU cell embeddings enable cell annotation with joint differential gene expression and splicing analysis in brain non-myeloid tissue of *Tabula Muris*. a,b** UMAP of the joint GE-TU cell embeddings coloured by Leiden cluster (**a**) and *Tabula Muris* reference cell type (**b**), showing close correspondence between the annotations and the inferred Leiden clusters (0: OCs, 1: ECs, 2: astrocytes, 3: OPCs, 4: brain pericytes, 5: astrocytes). Leiden cluster 5 is a small cluster of three astrocytes that we omit from the analysis. **c** Mean gene expression values of the top three differentially expressed genes (DEGs) per cluster identify cell-type marker genes. **d** Mean PSI values of the top three differentially spliced gene (DSG) isoforms per cluster identify candidate cell-type-specific isoform markers.

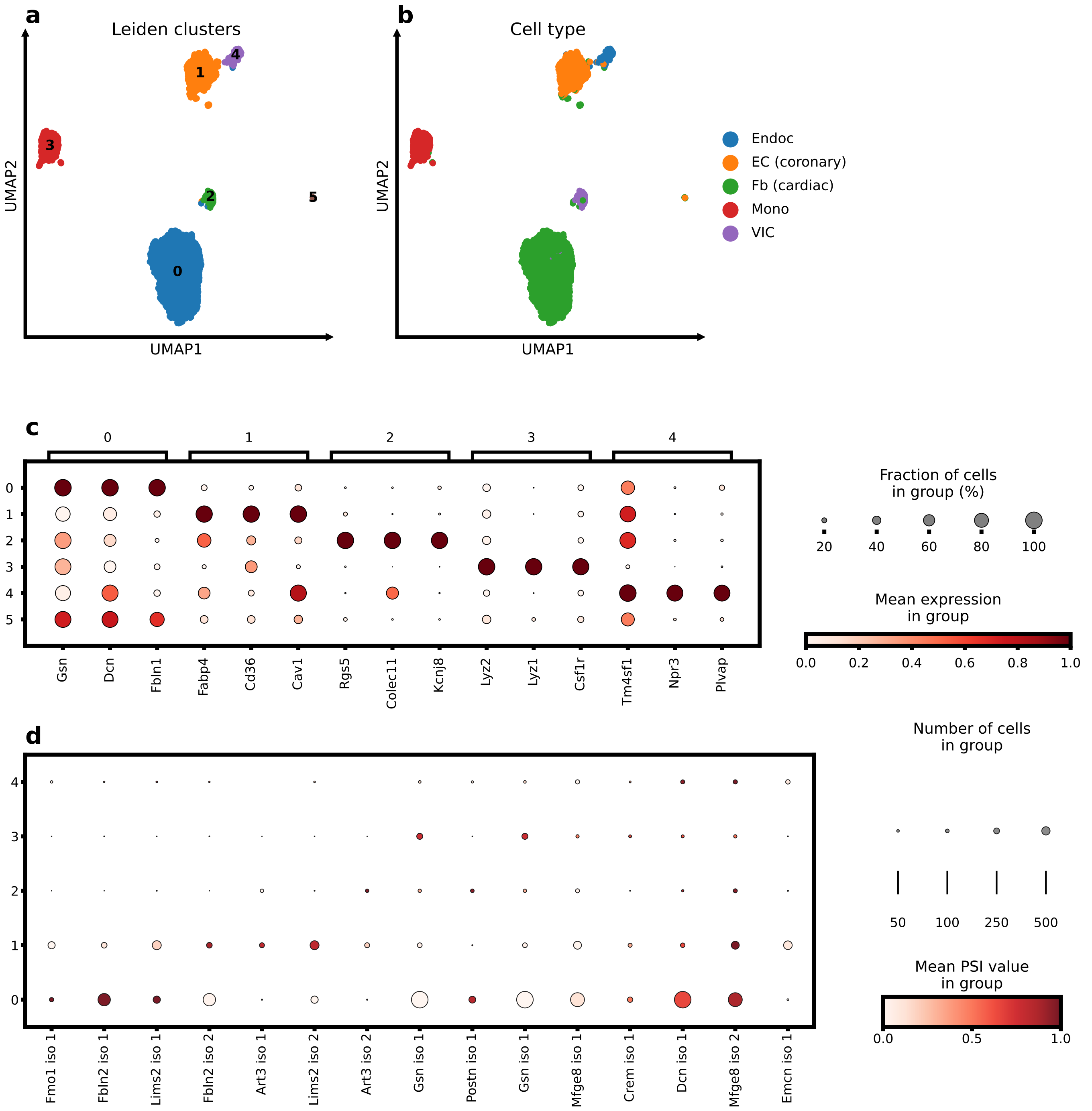

**Supplementary Fig. 24 — TRVI's GE-TU cell embeddings enable cell annotation with joint differential gene expression and splicing analysis in heart tissue of *Tabula Muris*. a,b** UMAP of the joint GE-TU cell embeddings coloured by Leiden cluster (**a**) and *Tabula Muris* reference cell type (**b**), showing close correspondence between the annotations and the inferred Leiden clusters (0: Fbs (cardiac), 1: ECs (coronary), 2: VICs, 3: Monos, 4: Endocs, 5: EC (coronary) and Fb (cardiac). Leiden cluster 5 is a small cluster of one EC (coronary) and one Fb (cardiac) cell that we omit from the analysis. **c** Mean gene expression values of the top three differentially expressed genes (DEGs) per cluster identify cell-type marker genes. **d** Mean PSI values of the top three differentially spliced gene (DSG) isoforms per cluster identify candidate cell-type-specific isoform markers.

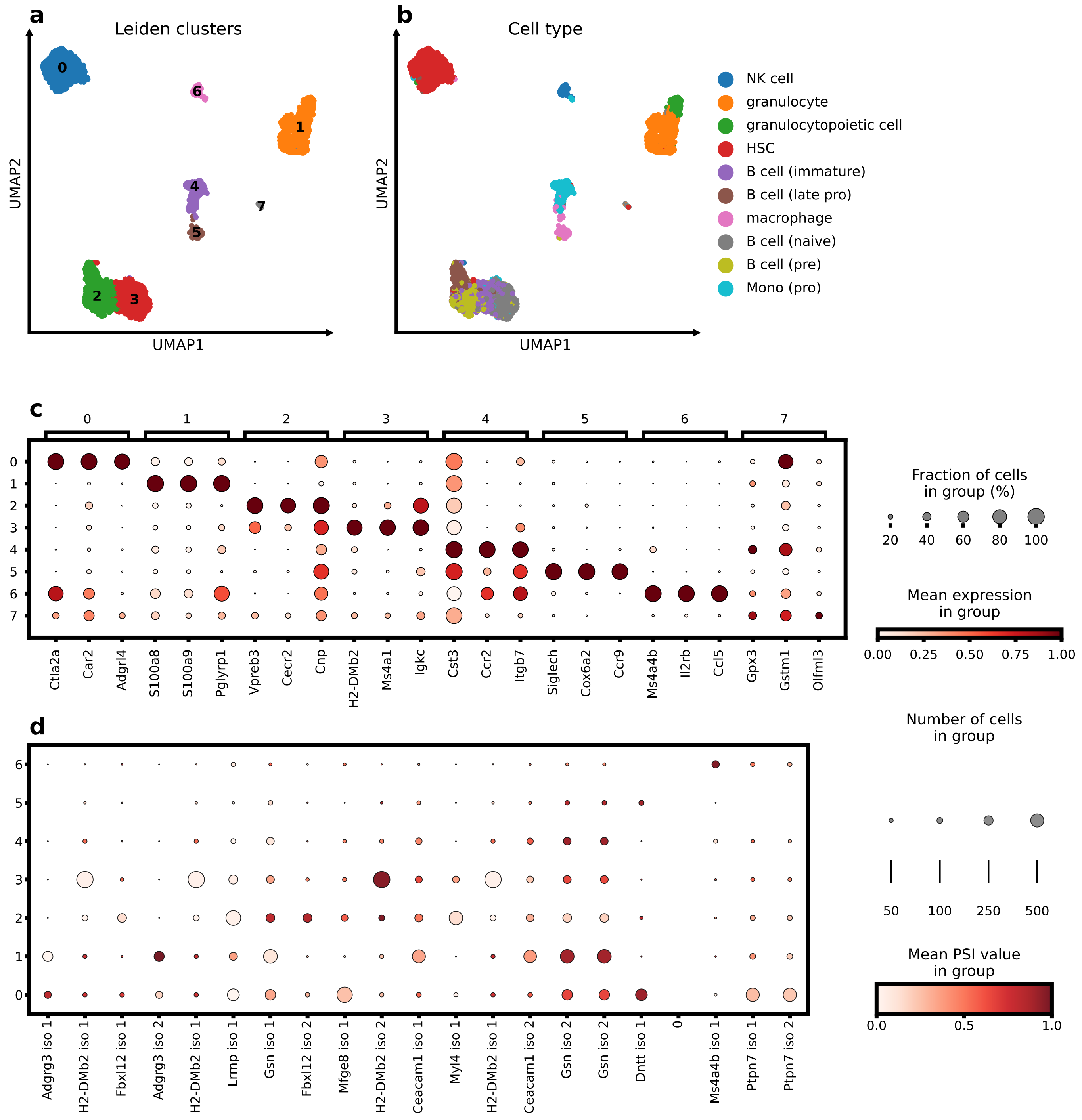

**Supplementary Fig. 25 — TRVI's GE-TU cell embeddings enable cell annotation with joint differential gene expression and splicing analysis in bone marrow tissue of *Tabula Muris*. a,b** UMAP of the joint GE-TU cell embeddings coloured by Leiden cluster (**a**) and *Tabula Muris* reference cell type (**b**), demonstrating close correspondence between the annotations and the inferred Leiden clusters (0: HSC, 1: granulocytes, granulocytopoietic cells, 2: B cells (late pro), B cells (pre), 3: B cells (immature), B cells (naïve), 4: Monos (pro), 5: macrophages, 6: NK cells, 7: B cells (naïve)). Leiden cluster 5 is a small cluster of naïve B cells and HSC (coronary). The four annotated B cell subtypes are captured by two structured clusters (2: B cells (late pro), B cells (pre)); 3: B cells (immature), B cells (naïve). However, the two Leiden clusters are structured into the respective sub-cell types. **c** Mean gene expression values of the top three differentially expressed genes (DEGs) per cluster identify cell-type marker genes. **d** Mean PSI values of the top three differentially spliced gene (DSG) isoforms per cluster identify candidate cell-type-specific isoform markers. For Leiden cluster 5 (macrophages) only two isoforms were statistically significant (*Gsn* iso2, *Dtn* iso 1) as indicated by the empty column 0.

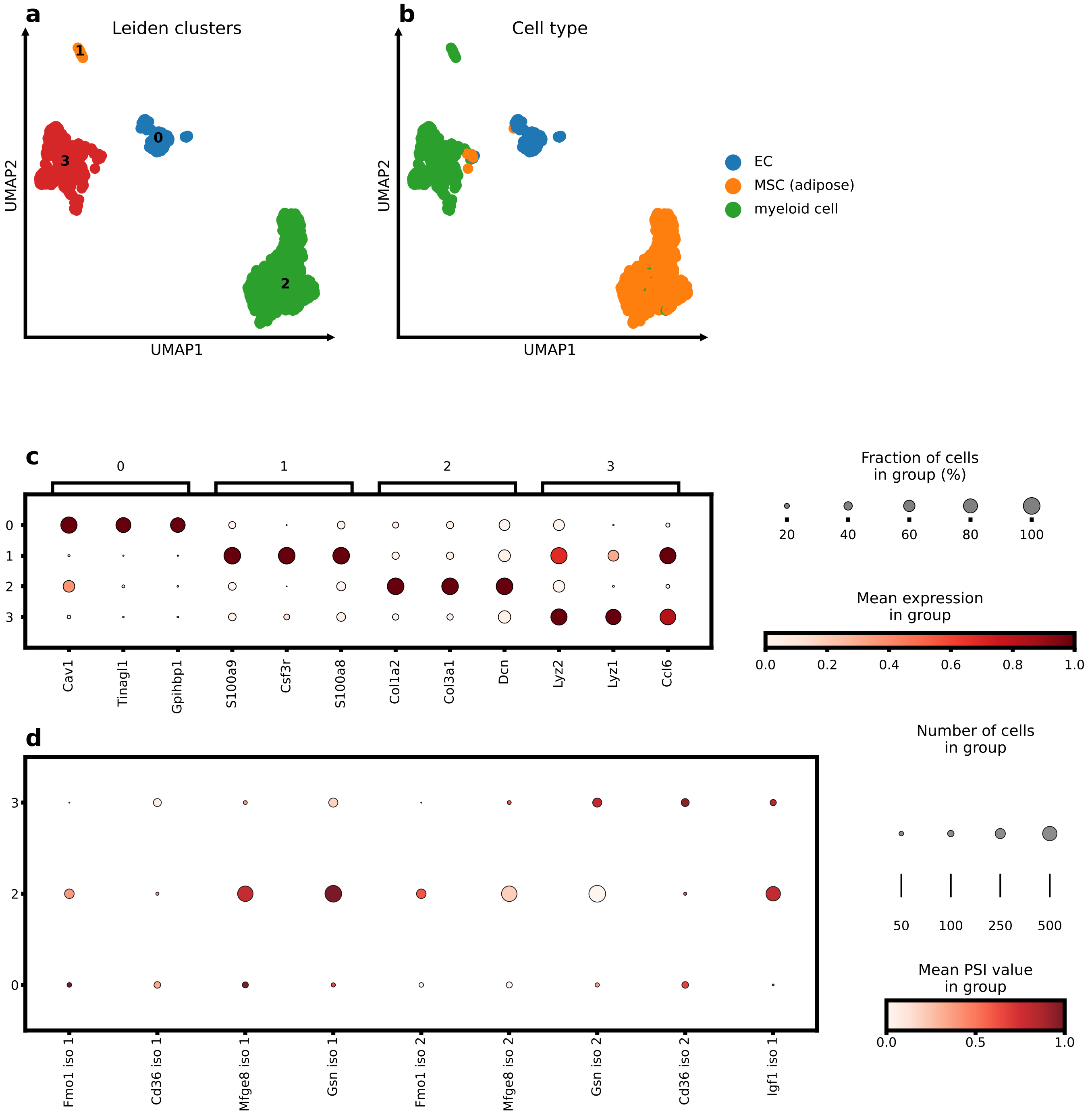

**Supplementary Fig. 26 — TRVI's GE-TU cell embeddings enable cell annotation with joint differential gene expression and splicing analysis in gonadal adipose tissue of *Tabula Muris*. a,b** UMAP of the joint GE-TU cell embeddings coloured by Leiden cluster (**a**) and *Tabula Muris* reference cell type (**b**), demonstrating close correspondence between the annotations and the inferred Leiden clusters (0: EC, 1: myeloid cells, granulocytopoietic cells, 2: MSC (adipose), 3: myeloid cells). Leiden cluster 1 is a small cluster of myeloid cells too small for DSG analysis and is therefore omitted. **c** Mean gene expression values of the top three differentially expressed genes (DEGs) per cluster identify cell-type marker genes. **d** Mean PSI values of the top three differentially spliced gene (DSG) isoforms per cluster identify candidate cell-type-specific isoform markers.

### Supplementary Note 1: Training objective of transcript usage Variational Inference (tuVI)

To determine the optimal parameters $\boldsymbol{\theta}_{TU}, \boldsymbol{\phi}_{TU}$ of the deep neural networks of the encoder and decoder of tuVI, the evidence lower bound (ELBO) is optimised [1]. For a single-cell transcript usage profile $\mathbf{x}_n^{(TU)}$ it is given by

$$\begin{aligned}\log p_{\boldsymbol{\theta}_{TU}}\left(\mathbf{x}_n^{(TU)}\right) &\geq \mathcal{L}\left(\boldsymbol{\theta}_{TU}, \boldsymbol{\phi}_{TU}; \mathbf{x}_n^{(TU)}\right) \\ &= \mathbb{E}_{\mathbf{z}_n \sim q_{\boldsymbol{\phi}_{TU}}(\mathbf{z}_n|\mathbf{x}_n^{(TU)})}\left[\log p_{\boldsymbol{\theta}_{TU}}\left(\mathbf{x}_n^{(TU)}|\mathbf{z}_n\right)\right] - \mathrm{D}_{\mathrm{KL}}\left(q_{\boldsymbol{\phi}_{TU}}(\mathbf{z}_n|\mathbf{x}_n^{(TU)})\middle\|p(\mathbf{z}_n)\right)\end{aligned} \tag{1}$$

where the expected log-likelihood evaluates the data generation fidelity and the Kullback-Leibler regularises the variational posterior towards the standard Gaussian prior. For the specified Gaussian variational posterior $q_{\boldsymbol{\phi}_{TU}}(\mathbf{z}_n \mid \mathbf{x}_n^{(TU)}) = \mathcal{N}\left(\mathbf{z}_n|\boldsymbol{\mu}_{\boldsymbol{\phi}_{TU}}\left(\mathbf{x}_n^{(TU)}\right), \operatorname{diag}\left(\boldsymbol{\sigma}^2_{\boldsymbol{\phi}_{TU}}\left(\mathbf{x}_n^{(TU)}\right)\right)\right)$ and the standard multivariate Gaussian prior $p(\mathbf{z}_n) = \mathcal{N}(\mathbf{0}, \mathbf{I}_L)$, the negative Kullback-Leibler divergence $\mathrm{D}_{\mathrm{KL}}$ can be obtained in closed-form by

$$-\mathrm{D}_{\mathrm{KL}}\left(q_{\boldsymbol{\phi}_{TU}}(\mathbf{z}_n \mid \mathbf{x}_n^{(TU)})\middle\|p(\mathbf{z}_n)\right) = \frac{1}{2}\sum_{l=1}^{L}\left(1 + 2\log(\sigma_{n,l}) - \mu_{n,l}^2 - \sigma_{n,l}^2\right). \tag{2}$$

with $\mu_{n,l} \coloneqq (\boldsymbol{\mu}_{\boldsymbol{\phi}_{TU}}(\mathbf{x}_n^{(TU)}))_l$ and $\sigma_{n,l} \coloneqq (\boldsymbol{\sigma}_{\boldsymbol{\phi}_{TU}}(\mathbf{x}_n^{(TU)}))_l$.

However, the expected log-likelihood in equation (1) cannot be computed analytically. Therefore Monte-Carlo sampling yields an approximate solution. Following [1], sampling from the Gaussian variational posterior $\mathbf{z}_n \sim q_{\boldsymbol{\phi}_{TU}}(\mathbf{z}_n \mid \mathbf{x}_n^{(TU)})$ is enabled through the reparameterisation-trick of the latent cell embedding $\mathbf{z}_n$ as

$$\tilde{\mathbf{z}}_n^{(k)} = \boldsymbol{\mu}_{\boldsymbol{\phi}_{TU}}(\mathbf{x}_n^{(TU)}) + \boldsymbol{\sigma}_{\boldsymbol{\phi}_{TU}}(\mathbf{x}_n^{(TU)}) \odot \boldsymbol{\epsilon}^{(k)} \text{ with } \boldsymbol{\epsilon}^{(k)} \sim \mathcal{N}(\mathbf{0}, \mathbf{I}_L). \tag{3}$$

Then, Monte-Carlo $K$ samples can be drawn and the intractable expected log-likelihood is approximated via

$$\mathbb{E}_{\mathbf{z}_n \sim q_{\boldsymbol{\phi}_{TU}}(\mathbf{z}_n|\mathbf{x}_n^{(TU)})}\left[\log p_{\boldsymbol{\theta}_{TU}}\left(\mathbf{x}_n^{(TU)}|\mathbf{z}_n\right)\right] \approx \frac{1}{K}\sum_{k=1}^{K}\log p_{\boldsymbol{\theta}_{TU}}(\mathbf{x}_n^{(TU)}|\tilde{\mathbf{z}}_n^{(k)}). \tag{4}$$

We used $K = 1$ samples for tuVI as done in standard variational autoencoders [1]. For a dataset containing $N$ cells, the full-data ELBO is the sum of the per-cell ELBOs yielding

$$\mathcal{L}^{\text{total}}\left(\boldsymbol{\theta}_{TU}, \boldsymbol{\phi}_{TU}; \mathbf{X}^{(TU)}\right) = \sum_{n=1}^{N}\mathcal{L}\left(\boldsymbol{\theta}_{TU}, \boldsymbol{\phi}_{TU}; \mathbf{x}_n^{(TU)}\right). \tag{5}$$

Applying uniformly sampled mini-batches $\mathcal{B}$ of size $M = |\mathcal{B}|$, an unbiased estimator of the full-data ELBO is obtained by

$$\widehat{\mathcal{L}}^{\text{total}}_{\mathcal{B}} = \frac{N}{M}\sum_{n \in \mathcal{B}}\widehat{\mathcal{L}}\left(\boldsymbol{\theta}_{TU}, \boldsymbol{\phi}_{TU}; \mathbf{x}_n^{(TU)}\right). \tag{6}$$

For the DM and ZANIDM observation models, this training objective is a valid lower bound on the log evidence (ELBO). When the heuristic ZIDM is used, the expected log-likelihood is replaced by the expected ZIDM surrogate score $H^{\text{ZIDM}}$. The resulting training objective retains the KL-divergence regularisation but is a surrogate objective rather than a formal evidence lower bound because $H^{\text{ZIDM}}$ is a heuristic and not a valid log-likelihood.

### Supplementary Note 2: Observation models of alternative splicing-induced transcript usage data

In contrast to gene expression data, transcript usage data are compositional, because one gene $g \in G$ can encode several isoforms through alternative splicing (Supplementary Fig. 2). Following existing literature for unsupervised modelling of splicing [2, 3], each spliced gene vector $\mathbf{x}_{n,g}^{(TU)}$ of cell $n$ consists of stacked, independent isoform group count vectors $\mathbf{x}_{n,g,i}^{(TU)}$ that are also known as intron group vectors. Each element of an isoform group count vector quantifies an exon junction (intron) through matching read counts indicating the presence of the corresponding spliced isoform. The number of isoform groups $I_g$ can vary for different spliced genes. Similarly, the number of exon junctions per isoform group can also vary. Thus, the data likelihood $p_{\boldsymbol{\theta}_{TU}}\left(\mathbf{x}_n^{(TU)}|\mathbf{z}_n\right)$ of a cell $n$ factorises into

$$p_{\boldsymbol{\theta}_{TU}}\left(\mathbf{x}_n^{(TU)}|\mathbf{z}_n\right) = \prod_{g=1}^{G}\prod_{i=1}^{I_g} p_{\boldsymbol{\theta}_{TU}}\left(\mathbf{x}_{n,g,i}^{(TU)}|\mathbf{z}_n\right). \tag{7}$$

where $p_{\boldsymbol{\theta}_{TU}}\left(\mathbf{x}_{n,g,i}^{(TU)}|\mathbf{z}_n\right)$ is the transcript usage observation model assessing the plausibility of $\mathbf{x}_{n,g,i}^{(TU)}$ given the latent cell embedding $\mathbf{z}_n$ and the parameters $\boldsymbol{\theta}_{TU}$. To train tuVI and TRVI, the log-likelihoods of the transcript usage observation models are necessary. We therefore adopt the notation simplification throughout the remainder of this Supplmentary Note: $\mathbf{x}^{TU} \coloneqq \mathbf{x}_{n,g,i}^{(TU)}$, $\mathbf{z} \coloneqq \mathbf{z}_n$, $\boldsymbol{\alpha} \coloneqq \boldsymbol{\alpha}_{n,g,i}$, $\boldsymbol{\zeta} \coloneqq \boldsymbol{\zeta}_{n,g,i}$, $\boldsymbol{\rho} \coloneqq \boldsymbol{\rho}_{n,g,i}$, and $D_{g,i} \coloneqq D$.

#### 2.1 Log-likelihood of Dirichlet-Multinomial distribution

To optimise the training objectives of tuVI or TRVI, the log-likelihood of the Dirichlet-Multinomial observation model is obtained by

$$\begin{aligned}\log \mathrm{DM}(\mathbf{x}^{(TU)}|N,\boldsymbol{\alpha}) = &\log\Gamma(\alpha_0) + \log\Gamma(N+1) - \log\Gamma(N+\alpha_0)\\ &+\sum_{d=1}^{D}\log\Gamma(x_d^{(TU)}+\alpha_d) - \log\Gamma(\alpha_d) - \log\Gamma(x_d^{(TU)}+1).\end{aligned} \tag{8}$$

#### 2.2 Log-likelihood of Zero-and-N-inflated Dirichlet Multinomial distribution

To optimise the training objectives of tuVI or TRVI with a ZANIDM observation model, the log-likelihood has to be computed. Dependent on the observed isoform group count vector $\mathbf{x}^{(TU)}$, the ZANIDM distribution can be broken down into the four cases and their respective log-likelihood is computed. For case 1 ($N > 0 \wedge \forall d\, x_d^{(TU)} > 0$), the log-likelihood is obtained by

$$\begin{aligned}\log \mathrm{ZANIDM}(\mathbf{x}^{(TU)}|\boldsymbol{\alpha},\boldsymbol{\zeta},N) = &\log\eta^{(D)} + \log\Gamma(\alpha_0) + \log\Gamma(N+1) - \log\Gamma(N+\alpha_0)\\ &+\sum_{d=1}^{D}\log\Gamma(x_d^{(TU)}+\alpha_d) - \log\Gamma(\alpha_d) - \log\Gamma(x_d^{(TU)}+1).\end{aligned} \tag{9}$$

For case 2 ($N = 0 \wedge \mathbf{x}^{(TU)} = \mathbf{0}$), the log-likelihood is simply obtained form the mixture weight as

$$\log \mathrm{ZANIDM}(\mathbf{x}^{(TU)}|\boldsymbol{\alpha},\boldsymbol{\zeta},N) = \log\eta^{(0)}. \tag{10}$$

For case 3 ($N > 0$ and $D-1$ zero-inflated intron counts), the set of zero-inflated subsets $\tilde{\mathcal{K}} = \{\}$ is empty by definition. The log-likelihood is of case 3 then results in

$$\begin{aligned}\log \mathrm{ZANIDM}(\mathbf{x}^{(TU)}|\boldsymbol{\alpha},\boldsymbol{\zeta},N) = &\log(\eta^{(D)}\frac{\Gamma(\alpha_0)\Gamma(N+1)}{\Gamma(N+\alpha_0)}\prod_{d=1}^{D}\frac{\Gamma\left(x_d^{(TU)}+\alpha_d\right)}{\Gamma(\alpha_d)\Gamma\left(x_d^{(TU)}+1\right)}\\ &+\sum_{d=1}^{D}\eta_d^{(N)}\left(\mathbf{1}_0\left(\sum_{k:k\neq d}x_k^{(TU)}\right)\right))\end{aligned}. \tag{11}$$

For case 4 ($N > 0$ and at most $D-2$ zero-inflated intron counts), the log-likelihood is then given by

$$\log \mathrm{ZANIDM}(\mathbf{x}^{(g,i)}|\boldsymbol{\alpha},\boldsymbol{\zeta},N) = \log(\eta^{(D)}\frac{\Gamma(\alpha_0)\Gamma(N+1)}{\Gamma(N+\alpha_0)}\prod_{d=1}^{D}\frac{\Gamma\left(x_d^{(TU)}+\alpha_d\right)}{\Gamma(\alpha_d)\Gamma\left(x_d^{(TU)}+1\right)}$$
$$+\sum_{\mathcal{K}\in\tilde{\mathcal{K}}}\eta_{\mathcal{K}}^{(\tilde{\mathcal{K}})}\left(\mathbf{1}_0\left(\sum_{j\in\mathcal{K}}x_j^{(TU)}\right)\right) \quad . \tag{12}$$
$$\frac{\Gamma(\alpha_{\mathcal{K}})\Gamma(N+1)}{\Gamma\left(N+\alpha_{\mathcal{K}}\right)}\prod_{d\notin\mathcal{K}}\frac{\Gamma\left(x_d^{(TU)}+\alpha_d\right)}{\Gamma(\alpha_d)\Gamma\left(x_d^{(TU)}+1\right)})$$

Although the log-likelihood of the different ZANIDM cases can be computed, the necessity to categorise each isoform group count vector $\mathbf{x}^{(TU)}$ of cell $n$ into one of the different cases, to compute the mixture weights $\boldsymbol{\eta}$ coupling the different ZANIDM cases, and to compute the subsets $\mathcal{K}$ for case 4, prevents an efficient parallelisation and increases the computation time substantially.

#### 2.3 Heuristic approximation of the ZANIDM log-likelihood

To overcome the computational burdens of a ZANIDM observation model, we seek to find a simpler heuristic of a mixture distribution comprising a point mass zero-inflation and the standard DM observation model. Since we only need the observation model to train tuVI but not for inferring latent cell embeddings later, we do not necessarily require a probability mass function. Instead, we can be satisfied with an easier-to-compute heuristic that, when employed within tuVI's objective, results in a latent space with a geometry similar to tuVI trained with a ZANIDM log-likelihood. Particularly, we want to remove the time-demanding computation of the ZANIDM mixture weights $\boldsymbol{\eta}$ from the excess-of-zero parameters $\boldsymbol{\zeta}$, and the computation of the subsets $\mathcal{K}$. Also, we want to reduce the number of cases and only focus on the case 1 and case 2 since a DM observation model in principle could model cases 3 and 4 although ill-suited for N-inflations as shown by [4]. As orientation, we use the zero-inflated negative binomial (ZINB) distribution that is used for gene expression data [5] and was efficiently implemented in scVI without exacerbating the computational time [6]. Accordingly, a putative standard zero-inflation mixture distribution for an isform group count vector $\mathbf{x}^{(TU)}$ consists of a point-mass for full zeros (case 2: $N = 0 \wedge \mathbf{x}^{(TU)} = \mathbf{0}$) and a standard observation model $p_{\boldsymbol{\theta}_{TU}}(\mathbf{x}^{(TU)} \mid \mathbf{z})$ for $N > 0$. The PMF is then obtained by

$$\mathrm{ZI}(\mathbf{x}^{(TU)}|\zeta_0,\boldsymbol{\theta}_{TU}) = \begin{cases}\zeta_0 + (1-\zeta_0)p_{\boldsymbol{\theta}_{TU}}(\mathbf{x}^{(TU)} \mid \mathbf{z}) & \text{if } N = 0\\ (1-\zeta_0)p_{\boldsymbol{\theta}_{TU}}(\mathbf{x}^{(TU)} \mid \mathbf{z}) & \text{if } N > 0\end{cases} \tag{13}$$

where $\zeta_0 \in [0,1]$ is the excess-of-zero probability for an entire isoform group count vector $\mathbf{x}^{(TU)}$. The corresponding, numerically stable log-likelihood for training the tuVI objective is then given by

$$\log \mathrm{ZI}(\mathbf{x}^{(TU)}|\rho_0,\boldsymbol{\theta}_{TU}) = \begin{cases}\mathcal{S}(-\rho_0 + \log p_{\boldsymbol{\theta}_{TU}}(\mathbf{x}^{(TU)} \mid \mathbf{z})) - \mathcal{S}(-\rho_0) & \text{if } N = 0\\ -\rho_0 - \mathcal{S}(-\rho_0) + \log p_{\boldsymbol{\theta}_{TU}}(\mathbf{x}^{(TU)} \mid \mathbf{z}) & \text{if } N > 0\end{cases} \tag{14}$$

where $\rho_0 = \mathrm{logit}(\zeta_0) \in \mathbb{R}$ and $\mathcal{S}(\cdot)$ is the softplus function. This log-likelihood is now used to determine a heuristic surrogate of the ZANIDM log-likelihood (equations 9, 10, 11, 12). To have an excess-of-zero parameter $\zeta_d \in [0,1]$ for each exon junction $d$ of the isoform group count vector $\mathbf{x}^{(TU)}$ as for the ZANIDM, we let the decoder of tuVI compute the logits of the the excess-of-zero parameters for each exon junction

$$\boldsymbol{\rho} = \begin{bmatrix}\rho_1\\ \vdots\\ \rho_D\end{bmatrix} = \mathrm{logit}(\boldsymbol{\zeta}) \in \mathbb{R}^D. \tag{15}$$

Computing the excess-of-zero logits directly not only enhances the numerical stability but also mimics the $\rho_0$ of the ZI log-likelihood (equation 14) although more flexible since $\rho_0$ is only the excess of zero probability for the entire isoform group vector $\mathbf{x}^{(TU)}$. In contrast, $\boldsymbol{\rho}$ can model excess-of-zeros for each intron $d$ in $\mathbf{x}^{(TU)}$. To account for the case of no exon junction counts $\mathbf{x}^{(TU)} = \mathbf{0}, N = 0$, we calculate for the isoform group for each mini-batch $\mathcal{B}$ of size $|\mathcal{B}| = M$ an average number of total exon junction

$$\overline{N}^{(\mathcal{B})} = \left\lceil \frac{1}{M} \sum_{n \in \mathcal{B}} (N_n + 1) \right\rceil \tag{16}$$

that is at least $\overline{N}^{(\mathcal{B})} = 1$. In the case of no exon junction counts $\mathbf{x}^{(TU)} = \mathbf{0}, N = 0$, this average count $\overline{N}^{(\mathcal{B})}$ is used to compute the term

$$h^{\mathrm{DM}}(\boldsymbol{\alpha}, \overline{N}^{(\mathcal{B})}) = \mathrm{sigmoid}\left( \log \frac{\Gamma(\alpha_0)\Gamma(\overline{N}^{(\mathcal{B})} + 1)}{\Gamma(\overline{N}^{(\mathcal{B})} + \alpha_0)} \right) \in [0, 1] \tag{17}$$

taking the parameters of the DM into account. With this term, we define a heuristic surrogate inspired by the ZANIDM log-likelihood as well as the general zero-inflation log-likelihood as

$$H^{\mathrm{ZIDM}}(\mathbf{x}^{(TU)} | \boldsymbol{\alpha}, \boldsymbol{\rho}, N, \overline{N}^{(\mathcal{B})}) = \begin{cases} \sum_{d=1}^{D} (\mathcal{S}(-\rho_d + \log(h^{\mathrm{DM}}(\boldsymbol{\alpha}, \overline{N}^{(\mathcal{B})}))) - \mathcal{S}(-\rho_d)) & \text{if } N = 0 \\ \sum_{d=1}^{D} (-\rho_d - \mathcal{S}(-\rho_d) + \log \mathrm{DM}\left(\mathbf{x}^{(TU)} | \boldsymbol{\alpha}, N\right)) & \text{if } N > 0 \end{cases} \tag{18}$$

where $\mathcal{S}(\cdot)$ is the softplus function. We refer to $H^{\mathrm{ZIDM}}$ as the heuristic ZIDM score and enable tuVI and TRVI to use it in place of the expected log-likelihood to accelerate training. Because this score is not the logarithm of a normalised probability distribution, the resulting training criterion is a surrogate variational objective rather than a strict ELBO. In contrast to the DM log-likelihood, the heuristic ZIDM score permits gradients to propagate from all-zero intron-group observations. We compared the latent cell embeddings obtained by tuVI using the ZANIDM or the heuristic ZIDM as described in Supplementary Note 5 and also benchmarked the computation time.

### Supplementary Note 3: Derivation of multi-modal Transcriptomic Regulation Variational Inference (TRVI)

For a set of modalities $\mathcal{M}$, let $\mathbf{X} = \{\mathbf{X}^{(r)}\}_{r\in\mathcal{M}}$ be the observed data of $N$ cells such that $\mathbf{x}_n = \{\mathbf{x}_n^{(r)}\}_{r\in\mathcal{M}}$ are the observed data modalities of a single cell $n$. We assume that $\mathbf{x}_n$ is generated from a shared latent variable $\mathbf{z}_n \in \mathbb{R}^L$, and $|\mathcal{M}|$ modality-specific latent variables $\{\mathbf{w}_n^{(r)} \in \mathbb{R}^L\}_{r\in\mathcal{M}}$ with respective priors $p(\mathbf{z}_n)$ and $\{p(\mathbf{w}_n^{(r)})\}_{r\in\mathcal{M}}$. Accordingly, the generative model of TRVI factorises as

$$p_{\mathbf{\Theta}}(\mathbf{x}_n, \mathbf{z}_n, \{\mathbf{w}_n^{(r)}\}_{r\in\mathcal{M}}) = p(\mathbf{z}_n) \prod_{r\in\mathcal{M}} p(\mathbf{w}_n^{(r)}) p_{\boldsymbol{\theta}_r}(\mathbf{x}_n^{(r)} \mid \mathbf{z}_n, \mathbf{w}_n^{(r)}) \tag{19}$$

where $\mathbf{\Theta}$ is the set of all parameters. For inference we need to marginalise the latent variables to obtain the evidence

$$p_{\mathbf{\Theta}}(\mathbf{x}_n) = \int p_{\mathbf{\Theta}}(\mathbf{x}_n, \mathbf{z}_n, \{\mathbf{w}_n^{(r)}\}_{r\in\mathcal{M}}) d\mathbf{z}_n \prod_{r\in\mathcal{M}} d\mathbf{w}_n^{(r)} \tag{20}$$

We first marginalise all modality-specific latent variables $\{\mathbf{w}_n^{(r)}\}_{r\in\mathcal{M}}$ from the distribution of the generative model yielding

$$\begin{aligned}
p_{\mathbf{\Theta}}(\mathbf{x}_n, \mathbf{z}_n) &= \int p_{\mathbf{\Theta}}(\mathbf{x}_n, \mathbf{z}_n, \{\mathbf{w}_n^{(r)}\}_{r\in\mathcal{M}}) \prod_{r\in\mathcal{M}} d\mathbf{w}_n^{(r)} \\
&= p(\mathbf{z}_n) \int \prod_{r\in\mathcal{M}} \Big( p(\mathbf{w}_n^{(r)}) p_{\boldsymbol{\theta}_r}(\mathbf{x}_n^{(r)} \mid \mathbf{z}_n, \mathbf{w}_n^{(r)}) \Big) \prod_{r\in\mathcal{M}} d\mathbf{w}_n^{(r)} \\
&= p(\mathbf{z}_n) \prod_{r\in\mathcal{M}} \int p(\mathbf{w}_n^{(r)}) p_{\boldsymbol{\theta}_r}(\mathbf{x}_n^{(r)} \mid \mathbf{z}_n, \mathbf{w}_n^{(r)}) d\mathbf{w}_n^{(r)} \\
&= p(\mathbf{z}_n) \prod_{r\in\mathcal{M}} p_{\boldsymbol{\theta}_r}(\mathbf{x}_n^{(r)} \mid \mathbf{z}_n)
\end{aligned} \tag{21}$$

where the third equality is obtained since each factor depends only on its individual modality-specific latent variable $\mathbf{w}_n^{(r)}$. To obtain the evidence $p_{\mathbf{\Theta}}(\mathbf{x}_n)$ we then marginalise over the shared latent variable $\mathbf{z}_n$ giving

$$\begin{aligned}
p_{\mathbf{\Theta}}(\mathbf{x}_n) &= \int p_{\mathbf{\Theta}}(\mathbf{x}_n, \mathbf{z}_n) d\mathbf{z}_n \\
&= \int p(\mathbf{z}_n) \prod_{r\in\mathcal{M}} p_{\boldsymbol{\theta}_r}(\mathbf{x}_n^{(r)} \mid \mathbf{z}_n) d\mathbf{z}_n.
\end{aligned} \tag{22}$$

However, the marginalisation of latent variables in variational autoencoders is generally intractable. Therefore we need to derive a lower bound on the log evidence and define first the modality-relevance-weighted mixture of experts variational posterior

$$q_{\mathbf{\Phi}^{\mathbf{z}}}(\mathbf{z}_n, \mid \mathbf{x}_n) = \sum_{m\in\mathcal{M}} \pi_n^{(m)} \, q_{\boldsymbol{\phi}_m^{\mathbf{z}}} \left( \mathbf{z}_n \mid \mathbf{x}_n^{(m)} \right) \tag{23}$$

where $\pi_n^{(m)} = \pi_{\boldsymbol{\phi}^\pi}^{(m)}(\mathbf{x}_n)$ is the modality-relevance weight following the constraints

$$\pi_{\boldsymbol{\phi}^\pi}^{(m)}(\mathbf{x}_n) \geq 0 \quad \text{for every}\, m \in \mathcal{M}, \qquad \sum_{m\in\mathcal{M}} \pi_{\boldsymbol{\phi}^\pi}^{(m)}(\mathbf{x}_n) = 1. \tag{24}$$

Although the modality-relevance weights are learnt functions of the observations, for fixed $\mathbf{x}_n$, they are constants with respect to integration over the latent variable $\mathbf{z}_n$. Having defined the modality-relevance-weighted mixture of experts variational posterior, we transform the evidence into

$$
\begin{aligned}
p_{\Theta}(\mathbf{x}_n) &= \int p(\mathbf{z}_n) \prod_{r\in\mathcal{M}} p_{\boldsymbol{\theta}_r}(\mathbf{x}_n^{(r)} \mid \mathbf{z}_n) d\mathbf{z}_n \\
&= \int q_{\boldsymbol{\Phi}^{\mathbf{z}}}(\mathbf{z}_n \mid \mathbf{x}_n) \frac{p(\mathbf{z}_n)}{q_{\boldsymbol{\Phi}^{\mathbf{z}}}(\mathbf{z}_n \mid \mathbf{x}_n)} \prod_{r\in\mathcal{M}} p_{\boldsymbol{\theta}_r}(\mathbf{x}_n^{(r)} \mid \mathbf{z}_n) d\mathbf{z}_n \\
&= \int \sum_{m\in\mathcal{M}} \pi_n^{(m)} q_{\boldsymbol{\phi}_m^{\mathbf{z}}}\left(\mathbf{z}_n \mid \mathbf{x}_n^{(m)}\right) \frac{p(\mathbf{z}_n)}{q_{\boldsymbol{\Phi}^{\mathbf{z}}}(\mathbf{z}_n \mid \mathbf{x}_n)} \prod_{r\in\mathcal{M}} p_{\boldsymbol{\theta}_r}(\mathbf{x}_n^{(r)} \mid \mathbf{z}_n) d\mathbf{z}_n \\
&= \sum_{m\in\mathcal{M}} \pi_n^{(m)} \int q_{\boldsymbol{\phi}_m^{\mathbf{z}}}\left(\mathbf{z}_n \mid \mathbf{x}_n^{(m)}\right) \frac{p(\mathbf{z}_n)}{q_{\boldsymbol{\Phi}^{\mathbf{z}}}(\mathbf{z}_n \mid \mathbf{x}_n)} \prod_{r\in\mathcal{M}} p_{\boldsymbol{\theta}_r}(\mathbf{x}_n^{(r)} \mid \mathbf{z}_n) d\mathbf{z}_n \\
&= \sum_{m\in\mathcal{M}} \pi_n^{(m)} \int q_{\boldsymbol{\phi}_m^{\mathbf{z}}}\left(\mathbf{z}_n \mid \mathbf{x}_n^{(m)}\right) \frac{p_{\boldsymbol{\theta}_m}(\mathbf{x}_n^{(m)} \mid \mathbf{z}_n) p(\mathbf{z}_n)}{q_{\boldsymbol{\Phi}^{\mathbf{z}}}(\mathbf{z}_n \mid \mathbf{x}_n)} \prod_{\substack{r\in\mathcal{M}\\ r\neq m}} p_{\boldsymbol{\theta}_r}(\mathbf{x}_n^{(r)} \mid \mathbf{z}_n) d\mathbf{z}_n
\end{aligned}
\tag{25}
$$

where the penultimate line follows because of the linearity of integration and $\pi_n^{(m)}$ does not depend on the shared latent variable $\mathbf{z}_n$, and the last line by separating the joint distribution

$$
\prod_{r\in\mathcal{M}} p_{\boldsymbol{\theta}_r}(\mathbf{x}_n^{(r)} \mid \mathbf{z}_n) = p_{\boldsymbol{\theta}_m}(\mathbf{x}_n^{(m)} \mid \mathbf{z}_n) \prod_{\substack{r\in\mathcal{M}\\ r\neq m}} p_{\boldsymbol{\theta}_r}(\mathbf{x}_n^{(r)} \mid \mathbf{z}_n)
\tag{26}
$$

explicitly into the marginal likelihoods for the source modality $m$ and target modality $r \neq m$. However, both marginal likelihoods are also generally intractable. Therefore, we define the variational posterior $q_{\boldsymbol{\phi}_m^{\mathbf{w}}}\left(\mathbf{w}_n^{(m)} \mid \mathbf{x}_n^{(m)}\right)$ on the modality-specific latent variable $\mathbf{w}_n^{(m)}$ and insert it into the source modality $m$ marginal likelihood resulting in

$$
\begin{aligned}
p_{\boldsymbol{\theta}_m}(\mathbf{x}_n^{(m)} \mid \mathbf{z}_n) &= \int q_{\boldsymbol{\phi}_m^{\mathbf{w}}}\left(\mathbf{w}_n^{(m)} \mid \mathbf{x}_n^{(m)}\right) \frac{p_{\boldsymbol{\theta}_m}(\mathbf{x}_n^{(m)}, \mathbf{w}_n^{(m)} \mid \mathbf{z}_n)}{q_{\boldsymbol{\phi}_m^{\mathbf{w}}}\left(\mathbf{w}_n^{(m)} \mid \mathbf{x}_n^{(m)}\right)} d\mathbf{w}_n^{(m)} \\
&= \int q_{\boldsymbol{\phi}_m^{\mathbf{w}}}\left(\mathbf{w}_n^{(m)} \mid \mathbf{x}_n^{(m)}\right) \frac{p_{\boldsymbol{\theta}_m}(\mathbf{x}_n^{(m)} \mid \mathbf{z}_n, \mathbf{w}_n^{(m)}) p(\mathbf{w}_n^{(m)})}{q_{\boldsymbol{\phi}_m^{\mathbf{w}}}\left(\mathbf{w}_n^{(m)} \mid \mathbf{x}_n^{(m)}\right)} d\mathbf{w}_n^{(m)}.
\end{aligned}
\tag{27}
$$

In contrast, for the marginal likelihood of the target modality $r \neq m$, we use the auxiliary distribution $r_r(\tilde{\mathbf{w}}_n^{(r)})$ obtaining

$$
\begin{aligned}
p_{\boldsymbol{\theta}_r}(\mathbf{x}_n^{(r)} \mid \mathbf{z}_n) &= \int r_r(\tilde{\mathbf{w}}_n^{(r)}) \frac{p_{\boldsymbol{\theta}_r}(\mathbf{x}_n^{(r)}, \tilde{\mathbf{w}}_n^{(r)} \mid \mathbf{z}_n)}{r_r(\tilde{\mathbf{w}}_n^{(r)})} d\tilde{\mathbf{w}}_n^{(r)} \\
&= \int r_r(\tilde{\mathbf{w}}_n^{(r)}) \frac{p_{\boldsymbol{\theta}_r}(\mathbf{x}_n^{(r)} \mid \mathbf{z}_n, \tilde{\mathbf{w}}_n^{(r)}) p(\tilde{\mathbf{w}}_n^{(r)})}{r_r(\tilde{\mathbf{w}}_n^{(r)})} d\tilde{\mathbf{w}}_n^{(r)}.
\end{aligned}
\tag{28}
$$

With these two transformed marginal likelihoods for the source modality $m$ and target modality $r$, we expand the evidence

$$
\begin{aligned}
p_{\Theta}(\mathbf{x}_n) &= \sum_{m\in\mathcal{M}} \pi_n^{(m)} \int q_{\boldsymbol{\phi}_m^{\mathbf{z}}}\left(\mathbf{z}_n \mid \mathbf{x}_n^{(m)}\right) \frac{p_{\boldsymbol{\theta}_m}(\mathbf{x}_n^{(m)} \mid \mathbf{z}_n) p(\mathbf{z}_n)}{q_{\boldsymbol{\Phi}^{\mathbf{z}}}(\mathbf{z}_n \mid \mathbf{x}_n)} \prod_{\substack{r\in\mathcal{M}\\ r\neq m}} p_{\boldsymbol{\theta}_r}(\mathbf{x}_n^{(r)} \mid \mathbf{z}_n) d\mathbf{z}_n \\
&= \sum_{m\in\mathcal{M}} \pi_n^{(m)} \int g_m(\cdot \mid \mathbf{x}_n) D_{\boldsymbol{\Phi},\boldsymbol{\Theta}}^{(m)} d\mathbf{z}_n d\mathbf{w}_n^{(m)} \prod_{\substack{r\in\mathcal{M}\\ r\neq m}} d\tilde{\mathbf{w}}_n^{(r)} \\
&= \sum_{m\in\mathcal{M}} \pi_n^{(m)} \mathbb{E}_{g_m(\cdot\mid\mathbf{x}_n)}\left[D_{\boldsymbol{\Phi},\boldsymbol{\Theta}}^{(m)}\right]
\end{aligned}
\tag{29}
$$

where all sampling terms are given by

$$g_m(\cdot \mid \mathbf{x}_n) = q_{\phi_m^{\mathbf{z}}}\left(\mathbf{z}_n \mid \mathbf{x}_n^{(m)}\right) q_{\phi_m^{\mathbf{w}}}\left(\mathbf{w}_n^{(m)} \mid \mathbf{x}_n^{(m)}\right) \prod_{\substack{r\in\mathcal{M}\\ r\neq m}} r_r(\tilde{\mathbf{w}}_n^{(r)}), \tag{30}$$

and the importance-weight by

$$\begin{aligned} D_{\mathbf{\Phi},\mathbf{\Theta}}^{(m)} &= \frac{p(\mathbf{z}_n)}{q_{\mathbf{\Phi}^{\mathbf{z}}}(\mathbf{z}_n \mid \mathbf{x}_n)} \frac{p_{\boldsymbol{\theta}_m}(\mathbf{x}_n^{(m)} \mid \mathbf{z}_n, \mathbf{w}_n^{(m)}) p(\mathbf{w}_n^{(m)})}{q_{\phi_m^{\mathbf{w}}}\left(\mathbf{w}_n^{(m)} \mid \mathbf{x}_n^{(m)}\right)} \prod_{\substack{r\in\mathcal{M}\\ r\neq m}} \frac{p_{\boldsymbol{\theta}_r}(\mathbf{x}_n^{(r)} \mid \mathbf{z}_n, \tilde{\mathbf{w}}_n^{(r)}) p(\tilde{\mathbf{w}}_n^{(r)})}{r_r(\tilde{\mathbf{w}}_n^{(r)})} \\ &= \frac{p(\mathbf{z}_n)}{q_{\mathbf{\Phi}^{\mathbf{z}}}(\mathbf{z}_n \mid \mathbf{x}_n)} \frac{p_{\boldsymbol{\theta}_m}(\mathbf{x}_n^{(m)} \mid \mathbf{z}_n, \mathbf{w}_n^{(m)}) p(\mathbf{w}_n^{(m)})}{q_{\phi_m^{\mathbf{w}}}\left(\mathbf{w}_n^{(m)} \mid \mathbf{x}_n^{(m)}\right)} \prod_{\substack{r\in\mathcal{M}\\ r\neq m}} p_{\boldsymbol{\theta}_r}(\mathbf{x}_n^{(r)} \mid \mathbf{z}_n, \tilde{\mathbf{w}}_n^{(r)}) \end{aligned} \tag{31}$$

In TRVI, we choose each auxiliary distribution to equal the corresponding modality-specific latent prior (i.e. $r_r(\tilde{\mathbf{w}}_n^{(r)}) \equiv p(\tilde{\mathbf{w}}_n^{(r)})$) such that $p(\tilde{\mathbf{w}}_n^{(r)})/r_r(\tilde{\mathbf{w}}_n^{(r)}) = 1$. However, the modality-relevance-weighted expectation cannot be solved analytically. Therefore, we approximate it by drawing $K$ i.i.d. Monte Carlo sample tuples

$$\{\mathbf{z}_n^{(m,k)}, \mathbf{w}_n^{(m,k)}, \{\tilde{\mathbf{w}}_n^{(r,k)}\}_{r\neq m}\}_{k=1}^K \overset{\text{i.i.d.}}{\sim} g_m(\mathbf{z}_n, \mathbf{w}_n^{(m)}, \{\tilde{\mathbf{w}}_n^{(r)}\}_{r\neq m} \mid \mathbf{x}_n) \tag{32}$$

conditional on the source modality $m$. This gives the $K$-sample importance-weight estimator

$$\overline{D}_K^{(m)} = \frac{1}{K}\sum_{k=1}^K D_{\mathbf{\Phi},\mathbf{\Theta}}^{(m,k)} \tag{33}$$

where $D_{\mathbf{\Phi},\mathbf{\Theta}}^{(m,k)}$ denotes $D_{\mathbf{\Phi},\mathbf{\Theta}}^{(m)}$ evaluated at the $k$-th sampled tuple. Using this $K$-sample importance-weight estimator and the product proposal

$$g_m^K := \prod_{k=1}^K g_m \tag{34}$$

we apply Jensen's inequality to the log evidence yielding a valid lower bound as

$$\begin{aligned} \log p_{\mathbf{\Theta}}(\mathbf{x}_n) &= \log\left(\sum_{m\in\mathcal{M}} \pi_n^{(m)} \mathbb{E}_{g_m^K(\cdot|\mathbf{x}_n)}\left[\overline{D}_K^{(m)}\right]\right) \\ &\geq \sum_{m\in\mathcal{M}} \pi_n^{(m)} \mathbb{E}_{g_m^K(\cdot|\mathbf{x}_n)}\left[\log\left(\frac{1}{K}\sum_{k=1}^K D_{\mathbf{\Phi},\mathbf{\Theta}}^{(m,k)}\right)\right] \\ &:= \mathcal{L}_K^{\pi}(\mathbf{\Phi}, \mathbf{\Theta}; \mathbf{x}_n) \end{aligned} \tag{35}$$

This ELBO is a modality-relevance-weighted version of the ELBO defined in [7]. Furthermore, for gene expression and AS-induced transcript usage $\mathcal{M} = \{GE, TU\}$, the training objective of TRVI is recovered. Also, this proof shows that TRVI with its modality-relevance-weighted mixture-of-experts variational posterior can mathematically accommodate even more than two single-cell modalities and enable the study of single-cell multiomcs data (e.g. gene expression, AS-induced transcript usage, surface protein abundance, and DNA methylation in parallel) while maintaining a valid training objective. Consequently, TRVI's modality-relevance weighted mixture-of-experts variational posterior could then infer the contribution of each single-cell multiomics modality to represent a cell.

### Supplementary Note 4: Sparsity and compositionality of single-cell splicing data complicate their probabilistic modelling

Gene expression and alternative splicing-induced transcript usage are both captured through single-cell transcriptomics sequencing. This rapidly advancing technology enabled deeper sequencing depths and broader read coverage increasing the data fidelity. However, still only around 10 - 40 % of all transcripts in a cell are detected inducing the uncertainty if a gene or an spliced isoform is not present (biological zero), or was not captured due to technical deficiencies (technical zero) ([8]). Therefore, gene expression matrices and transcript usage matrices are sparse exhibiting a high abundance of zero counts referred to as zero-inflation in statistics. To account for this sparsity, gene expression data can be probabilistically modelled through the zero-inflated negative binomial distribution (ZINB) ([5]).

For alternative splicing-induced transcript usage data, this approach is hard to adopt because a gene is not quantified by just one number of counts. Instead, a spliced gene $g \in G$ in cell $n$ can have different isoforms described by isoform groups (intron groups) $i \in I_g$ where each isoform group is a count vector $\mathbf{x}_{n,g,i}^{(TU)} \in \mathbb{N}_0^{D_{g,i}}$ of of $D_{g,i}$ exon junctions in that group. Thus, transcript usage data are in contrast to gene expression data also compositional further complicating their probabilistic modelling. Adopting the notation simplification $\mathbf{x}^{TU} := \mathbf{x}_{n,g,i}^{(TU)}$, and $D_{g,i} := D$ for the remainder of this Supplmentary Note, the total number of exon junction counts observed by an isoform group count vector $\mathbf{x}^{(TU)}$ is given by $N = \sum_{d=1}^{D} x_d^{(TU)}$. The presence of a spliced isoform indicated through the usage of an exon junction is then analysed through four scenarios:

- Case 1: All nonzero i.e. each exon junction $d$ has counts observed ($N > 0 \wedge \forall d\, x_d^{(TU)} > 0$)
- Case 2: All zero i.e. none of the exon junctions has counts observed ($N = 0$ and $\mathbf{x}^{(TU)} = \mathbf{0}$)
- Case 3: One nonzero i.e. only one exon junction has counts present ($D - 1$ exon junction have zero counts, and one exon junction $d$ has counts $x_d^{(TU)} = N$)
- Case 4: Subsets are zero i.e. $D - 2$ exon junctions have counts observed ($N > 0$ and at most $D - 2$ exon junctions have counts)

Analysing the abundance of each case on the *Tabula Muris* dataset shows that the majority of observed isoform group vectors belong to case 2 followed by case 3 (Supplementary Fig. 3a). Excluding these two cases, the number of isoform group vectors to determine the presence of isoforms through case 1 and case 4 is extremely small. This conclusion is exacerbated by considering the relative numbers of the cases present in the *Tabula Muris* dataset (Supplementary Fig. 3b). The zero-inflation case 1 and the one nonzero case 2 comprise 99.2 % of all observed isoform group count vectors. The Dirichlet Multinomial as state-of-the-art observation model to probabilistically model transcript usage data does not adequately model the case 1 in its event space when learning cell embeddings using variational autoencoders. This case would yield a probablity of 1 and consequently a log-likelihood of 0 such that no information can be backpropagated during optimisation of the variational autoencoder's objective. Thus, a variational autoencoder with a Dirichlet-Multinomial observation model does not gain information from 92.0 % of the data.

Also, considering the case 2 for a hypothetical isoform group of $D = 3$ exon junctions, observed isoform group count vectors constitute events $\mathbf{\Omega} = \{[\hat{N}, 0, 0]^T, [0, \hat{N}, 0]^T, [0, 0, \hat{N}]^T\ \}$ that are winner-takes-all scenarios where one exon junction receives all counts present $\hat{N}$. The event space of the Dirichlet Multinomial observation model includes this scenario but it constitutes a boundary case where the concentration parameter $\boldsymbol{\alpha} = [\alpha_1, \alpha_2, \alpha_3]^T$ has one element maximised and the other ones minimised. This is called N-inflation where recent studies show that the Dirichlet-Multinomial is ill-posed to model it because the concentration should be distributed less extreme over its elements ([4]).

### Supplementary Note 5: Comparison of single-cell alternative splicing-induced transcript usage embeddings learnt by tuVI with a ZANIDM or ZIDM observation model

The ZANIDM is a valid probability distribution and can be used as observation model for the variational autoencoder tuVI to learn a latent space of cell embeddings from single-cell alternative splicing-induced transcript usage data but is difficult to parallelise. In contrast, the heuristic ZIDM surrogate is faster to compute and can be used as a component of the training objective to train tuVI . Considering the optimisation of tuVI-ZANIDM and tuVI-ZIDM on the full *Tabula Muris* atlas, both variational autoencoders converge but with different training durations. Comparing the average epoch time of tuVI-ZANIDM and tuVI-ZIDM (Supplementary Fig. 4a), the former requires $\sim$ 65 min for one run through the Tabula Muris dataset whereas the latter takes only $\sim$ 27 s for the same task. This 27-fold outperformance can be attributed to the ZIDM heuristic being computed in parallel in contrast to the log-likelihood of the ZANIDM. The same conclusion can be drawn by investigating the total training duration (Supplementary Fig. 4b). Here, tuVI-ZIDM completes the optimisation after $\sim$ 53 min which is less than tuVI-ZANIDM would need for one epoch. The total training duration of tuVI-ZANIDM takes $\sim$ 24 h.

Having shown that applying the heuristic ZIDM yields faster computation times, the question remains if it enables tuVI to learn a latent space of cell embeddings from single-cell alternative splicing-induced transcript usage data that preserves the geometry and biology captured by the latent space of tuVI-ZANIDM. Therefore, we consider the cell type centroids of the cell embeddings of tuVI-ZIDM and tuVI-ZANIDM, and compute the Euclidean distance between cell type centroid in each latent space. The distance matrices of the cell type centroid of tuVI-ZANIDM (Supplementary Fig. 4c) and tuVI-ZIDM (Supplementary Fig. 4d) show highly similar relative distances. Not only are the cluster proximities preserved but also the fine-grained distances between cell type centroids are highly similar. This leads to the conclusion that the tuVI-ZIDM yields a latent space where the geometry and biology is highly reminiscent of the latent space of tuVI-ZANIDM but also faster to compute.

### Supplementary Note 6: Model evaluation of tuVI and benchmarking of its cell embeddings against scVI

We evaluated tuVI with a Dirichlet Multinomial observation model (tuVI-DM), a zero-and-$N$-inflated Dirichlet Multinomial observation model (tuVI-ZANIDM) or the zero-inflated Dirichlet Multinomial heuristic (tuVI-ZIDM) on the held-out *Tabula Muris* test set. We first assessed sensitivity to random initialisation and examined the cell-type structure captured by each latent space. We then compared the selected tuVI cell embeddings, learnt from alternative splicing-induced transcript-usage profiles, with gene expression cell embeddings learnt by scVI using a zero-inflated negative-binomial observation model (scVI-ZINB).

Shared architectural and training settings were matched wherever applicable (Supplementary Tabs. 1 and 2). scVI retained its recommended default batch size of 128, whereas tuVI was trained with a batch size of 256.

#### 6.1 Robustness to random initialisation and seed selection

To assess sensitivity to random initialisation, we trained each tuVI variant (tuVI-DM, tuVI-ZANIDM, tuVI-ZIDM) with ten random seeds. Within each observation model, $S = 10$ seeds were compared using the mean conditional NLL on the held-out test set. For a tuVI variant trained with seed $s$ the NLL is then computed as

$$\mathrm{NLL}_{\mathrm{tuVI}}^{(s)} = -\frac{1}{N}\sum_{n=1}^{N} \log p_{\boldsymbol{\theta}^{(s)}}\left(\mathbf{x}_n^{(TU)} \mid \mathbf{z}_n^{(s)}\right) \tag{36}$$

where $N$ is the number of test cells and $\mathbf{z}_n^{(s)}$ is the inferred latent representation of cell $n$. Because DM, ZANIDM and ZIDM define different log-likelihood functions, their absolute NLL values are not directly comparable. We therefore used NLL only to assess variation among seeds within the same observation model and to select one seed per model.

Mean conditional NLL varied only marginally across seeds for all three observation models, and the standard deviation of the per-cell NLL was similarly consistent (Supplementary Tab. 4; Supplementary Figs. 5a,b, 6a,b and 7a,b). These results indicate that conditional reconstruction performance was robust to random initialisation.

We next examined UMAP projections of the embeddings learnt by the models with the lowest and highest mean conditional NLL within each observation model (Supplementary Figs. 5c,d, 6c,d and 7c,d). The tuVI-DM embeddings showed limited separation of annotated cell populations. By contrast, tuVI-ZANIDM and tuVI-ZIDM produced distinct cell-type-associated clusters for both seeds examined. Thus, although all three variants were robust to random initialisation, ZANIDM and ZIDM captured substantially more cell-type-informative patterns than DM.

For subsequent analyses, we selected the seed with the lowest mean conditional NLL within each observation model: seed 5 for tuVI-DM, seed 6 for tuVI-ZANIDM and seed 1 for tuVI-ZIDM (Supplementary Tab. 4).

#### 6.2 The alternative splicing-induced transcript usage cell embeddings learnt by tuVI are competitive with gene expression cell embeddings learnt by scVI

We quantified the biological structure captured by the selected tuVI-DM, tuVI-ZANIDM and tuVI-ZIDM alternative splicing-induced transcript usage cell embeddings and compared it with that captured by gene expression cell embeddings learnt by scVI-ZINB. NMI and ARI measured the agreement between embedding-derived clusters and the Tabula Muris cell-type annotations, whereas ASW measured the separation of annotated cell types in the latent space. AvgBio was calculated as the mean of NMI, ARI and ASW.

tuVI-ZANIDM and tuVI-ZIDM outperformed tuVI-DM across all four metrics (Supplementary Tab. 6), in agreement with the qualitative UMAP analysis. Also, we observe that tuVI-ZIDM as a simpler heuristic of tuVI-ZANIDM has highly similar performance in terms of the AvgBio scores though tuVI-ZANIDM achieved the higher NMI and ASW, whereas tuVI-ZIDM achieved the higher ARI.

The tuVI-ZANIDM and tuVI-ZIDM AS-indcued transcript usage cell embeddings were comparable to the scVI-ZINB cell gene expression embedding. scVI-ZINB achieved the highest NMI and ASW, tuVI-ZIDM achieved the highest ARI, and tuVI-ZANIDM achieved a marginally higher AvgBio score than scVI-ZINB (0.730 versus 0.727). Collectively, these results show that tuVI with ZANIDM or ZIDM learns cell-type-informative structure from alternative splicing-induced transcript usage, with overall agreement to the reference cell-type annotations comparable to that obtained from gene expression cell embeddings learnt by scVI-ZINB.

### Supplementary Note 7: tuVI cell embeddings identify cell-type-specific isoforms that complement gene expression markers in *Tabula Muris* brain non-myeloid tissue

To compare cell annotation based on gene expression and alternative splicing (AS), we analysed the *Tabula Muris* (TM) brain non-myeloid subset comprising astrocytes, brain pericytes, endothelial cells (ECs), oligodendrocytes (OCs), and oligodendrocyte precursor cells (OPCs). We used scVI to learn gene expression embeddings and tuVI to learn alternative splicing-induced transcript usage embeddings, and compared their Leiden clusters, differentially regulated genes and isoforms, and candidate cell-type markers.

#### 7.1 Clustering and differential analysis of gene expression and transcript usage embeddings

We constructed nearest-neighbour graphs and performed Leiden clustering separately for the two embeddings. The scVI embedding yielded five clusters corresponding to OCs, ECs, astrocytes, OPCs and brain pericytes, consistent with the TM reference annotation (Supplementary Fig. 8a,b). Cluster-specific differentially expressed genes (DEGs) were identified using a Wilcoxon rank-sum test. The top six DEGs per cluster included established cell-type markers (Supplementary Fig. 8c), and UMAP projections of the top-ranked DEGs confirmed their cell-type specificity (Supplementary Figs. 8d and 11).

The tuVI embedding yielded six clusters: clusters 0 and 3 corresponded to two OC populations, whereas clusters 1, 2, 4 and 5 corresponded to ECs, astrocytes, OPCs and brain pericytes, respectively (Supplementary Fig. 9a,b). The separation of OCs into two clusters, which was absent from the scVI gene expression cell embedding, suggests AS-related oligodendrocyte subpopulations. Differential splicing analysis identified cluster-specific differentially spliced gene (DSG) isoforms. Despite sparser splicing measurements, the mean percent-spliced-in (PSI) values of the top three DSG isoforms per cluster distinguished the corresponding cell types (Supplementary Fig. 9c). *Scarb1* isoform 1 and *Septin4* isoform 2 were shared across the two OC clusters, whereas *Lims2* isoform 1 and *Mgll* isoform 1 were enriched in cluster 0, nominating them as candidate subpopulation markers (Supplementary Fig. 12a,d). In ECs, the combined PSI profiles of *Scarb1* isoform 2, *Septin4* isoform 1 and *Nrep* isoform 1 distinguished the cluster more clearly than any single isoform (Supplementary Figs. 9d and 12b).

#### 7.2 Gene expression and transcript usage provide complementary cell-type markers

DSGs were substantially less numerous than DEGs and overlapped only partly with them (Supplementary Fig. 10; Supplementary Table 12). For OC cluster 0, ECs, astrocytes, OC cluster 3, OPCs and brain pericytes, respectively, the DEG/DSG counts were 95/11, 181/17, 152/14, 95/11, 193/17 and 129/7, of which 5, 5, 9, 6, 11 and 6 genes were shared. Moreover, only 1, 0, 4, 1, 4 and 0 DSGs, respectively, occurred within equally sized sets of top-ranked DEGs. The overlapping genes were *Septin4* in both OC clusters; *Clu*, *Ntrk2*, *Ntsr2* and *Scg3* in astrocytes; and *Cntn1*, *Cspg5*, *Nnat* and *Rtn1* in OPCs. Thus, most cell-type-associated transcript usage changes were not captured by ranking genes according to expression.

UMAP projections confirmed the cell-type specificity of the top three DEGs and showed that sets of DSG isoforms could similarly distinguish cell types, although individual isoforms were sometimes shared and measurements were sparse (Supplementary Figs. 11 and 12). Split-violin plots further demonstrated that isoform usage could be informative when gene-level expression was not (Supplementary Fig. 13). In OC cluster 0, *Scarb1*, *Lims2* and *Mgll* were not upregulated, whereas their PSI distributions distinguished OCs; *Lims2* and *Mgll* further differentiated cluster 0 from cluster 3 (Supplementary Fig. 13a). In ECs, the *Scarb1*, *Septin4* and *Nrep* isoforms were more specific than the corresponding gene expression profiles (Supplementary Fig. 13b). In astrocytes, *Rtn1*, *Limch1* and *Gpm6a* showed cell-type-specific PSI distributions, although only *Gpm6a* was upregulated at the gene level (Supplementary Fig. 13c). In OC cluster 3, *Scarb1* and *Septin4* supported general OC identity, whereas *Mcam* isoform usage was not cluster-specific (Supplementary Fig. 13d). In OPCs, *Gsn* and *Gpm6a*, but not *Lims2*, showed OPC-specific PSI distributions (Supplementary Fig. 13e). The candidate brain-pericyte isoforms were enriched but not exclusive to this sparsely represented population and therefore require cautious interpretation (Supplementary Fig. 13f).

Several candidate isoform markers are not canonical cell-type markers but have relevant disease contexts. *Scarb1* supports myelin maintenance and has been proposed as a therapeutic target in glioblastoma [9]; *Septin4* contributes to mature central nervous system myelin and has been linked to neurological disorders [10]. *Rtn1* alternative splicing has been associated with Alzheimer's disease [11]. *Limch1* has been implicated in astrocyte dysfunction in amyotrophic lateral sclerosis [12] whereas *Gpm6a* has been reported in OPC and astrocyte populations in glioblastoma [13].

Together, these results show that tuVI identifies candidate cell-type-specific isoforms that are largely distinct from the gene expression cell markers recovered by scVI. Alternative splicing-induced transcript usage additionally resolves OC substructure not apparent from gene expression, supporting gene expression and AS as complementary transcriptomic regulation mechanisms for cell identity.

### Supplementary Note 8: tuVI cell embeddings identify cell-type-specific isoforms that complement gene expression markers in *Tabula Muris* heart tissue

To compare cell annotation based on gene expression and alternative splicing (AS), we analysed the *Tabula Muris* (TM) heart subset comprising endocardial cells (Endocs), coronary endothelial cells (coronary ECs), cardiac fibroblasts (cardiac Fbs), monocytes (Monos) and valve interstitial cells (VICs). We used scVI to learn gene expression embeddings and tuVI to learn alternative splicing-induced transcript usage embeddings, and compared their Leiden clusters, differentially regulated genes and isoforms, and candidate cell-type markers.

#### 8.1 Clustering and differential analysis of gene expression and transcript usage embeddings

We constructed nearest-neighbour graphs and performed Leiden clustering separately for the two embeddings. The scVI embedding yielded six clusters (Supplementary Fig. 14a). Clusters 0–4 corresponded to cardiac Fbs, coronary ECs, VICs, Monos and Endocs, respectively, consistent with the TM reference annotation (Supplementary Fig. 14b). Cluster 5 contained a single cardiac Fb and was excluded from downstream analyses. Cluster-specific differentially expressed genes (DEGs) were identified using a Wilcoxon rank-sum test. The top six DEGs per cluster included established cell-type markers (Supplementary Fig. 14c), and UMAP projections of the top-ranked DEGs confirmed their cell-type specificity (Supplementary Figs. 14d and 17).

The tuVI embedding yielded five Leiden clusters with the same cell-type correspondence (Supplementary Fig. 15a,b). Differential splicing analysis identified cluster-specific differentially spliced gene (DSG) isoforms. Although splicing observations were sparser than gene expression measurements, the mean percent-spliced-in (PSI) values of the top three DSG isoforms per cluster distinguished the corresponding cell types (Supplementary Fig. 15c). For example, the PSI profiles of *Fmo1* isoform 1, *Fbln2* isoform 1 and *Lims2* isoform 1 discriminated cardiac Fbs (Supplementary Figs. 15d and 18a).

#### 8.2 Gene expression and transcript usage provide complementary cell-type markers

DSGs were substantially less numerous than DEGs and overlapped only partly with them (Supplementary Fig. 16; Supplementary Table 13). For cardiac Fbs, coronary ECs, VICs, Monos and Endocs, respectively, we identified 174/14, 160/16, 81/4, 183/8 and 110/3 DEGs/DSGs, of which 8, 11, 3, 3 and 2 genes were shared. Moreover, only two DSGs were represented among the correspondingly sized sets of top-ranked DEGs for cardiac Fbs (*Dcn* and *Gsn*) and coronary ECs (*Cd36* and *Lims2*); none overlapped for VICs, Monos or Endocs. Thus, most cell-type-associated transcript usage changes were not captured by ranking genes according to expression.

UMAP projections confirmed that both the top three DEGs and top three DSG isoforms were cell-type-specific, although DSG measurements were sparser (Supplementary Figs. 17 and 18). Split-violin plots further showed that isoform usage could distinguish cell types when gene-level expression was less informative (Supplementary Fig. 19). In cardiac Fbs, *Fmo1*, *Fbln2* and *Lims2* showed highly cell-type-specific PSI distributions, whereas their gene expression profiles were less specific (Supplementary Fig. 19a). In coronary ECs, the corresponding *Fbln2*, *Lims2* and *Art3* isoforms were cell-type-specific, although *Fbln2* expression was not (Supplementary Fig. 19b). In VICs, *Art3* and *Postn* showed concordant expression and PSI specificity, whereas *Gsn* isoform usage was inconclusive (Supplementary Fig. 19c). In Monos, *Gsn* showed a cell-type-specific PSI distribution despite non-specific gene expression, whereas *Mfge8* and *Crem* remained ambiguous (Supplementary Fig. 19d). No robust isoform marker was resolved for Endocs, potentially because this cell type was sparsely represented (Supplementary Fig. 19e).

Notably, cardiac Fbs and coronary ECs preferentially used different *Fbln2* and *Lims2* isoforms, and coronary ECs and VICs used different *Art3* isoforms (Supplementary Fig. 18). These examples show how gene-level overlap can conceal cell-type-specific isoform usage. Many candidates are not established marker genes but have relevant biological associations. *Fbln2* and *Lims2* have documented roles in cardiac development, repair, fibrosis, angiogenesis and cardiovascular disease [14–17]; *Art3* has primarily been studied in cancer signalling [18]; and monocyte-associated *Gsn* and *Crem* have been linked to acute aortic dissection and cancer, respectively [19, 20].

Together, these results show that tuVI identifies candidate cell-type-specific isoforms that are largely distinct from the gene expression markers recovered by scVI. Gene expression and AS therefore provide complementary transcritpomic regulation mechanisms of cell identity.

### Supplementary Note 9: Model selection and evaluation of TRVI

We evaluated the robustness of TRVI by training ten models with random seeds 0–9 on the *Tabula Muris* dataset. We first quantified modality-specific reconstruction performance on held-out test cells and selected a representative fitted model for visualisation and downstream analyses. We then examined whether variation in reconstruction performance across initialisations was accompanied by changes in the cell-type structure or geometry of the joint gene expression–transcript usage (GE–TU) latent space.

#### 9.1 Robustness to random intitialisation and seed selection

To assess sensitivity to random weight initialisation, we calculated the conditional negative log-likelihood (NLL) for each seed on the held-out *Tabula Muris* test set, separately for gene expression ($m = 1$) and alternative splicing-induced transcript usage ($m = 2$). For every test cell $n$, the relevance-weighted mixture-of-experts variational posterior inferred the joint GE–TU embedding $\mathbf{z}_n^{(s)}$, private gene expression (P-GE) embedding $\mathbf{w}_n^{(1,s)}$ and private transcript usage (P-TU) embedding $\mathbf{w}_n^{(2,s)}$ under seed $s$. These latent cell embeddings were passed to the modality-specific decoders to parameterise the ZINB observation model for gene expression, $p_{\boldsymbol{\theta}_{GE}^{(s)}}\left(\mathbf{x}_n^{(GE)} \mid \mathbf{z}_n^{(s)}, \mathbf{w}_n^{(GE,s)}\right)$, and the heuristic ZIDM surrogate for transcript usage, $\log p_{\boldsymbol{\theta}_{TU}^{(s)}}\left(\mathbf{x}_n^{(TU)} \mid \mathbf{z}_n^{(s)}, \mathbf{w}_n^{(TU,s)}\right)$. The mean per-cell NLL for seed $s$ was calculated as

$$\mathrm{NLL}_s^{(GE)} = -\frac{1}{N}\sum_{n=1}^{N} \log p_{\boldsymbol{\theta}_{GE}^{(s)}}\left(\mathbf{x}_n^{(GE)} \mid \mathbf{z}_n^{(s)}, \mathbf{w}_n^{(GE,s)}\right) \tag{37}$$

for gene expression and

$$\mathrm{NLL}_s^{(TU)} = -\frac{1}{N}\sum_{n=1}^{N} \log p_{\boldsymbol{\theta}_{TU}^{(s)}}\left(\mathbf{x}_n^{(TU)} \mid \mathbf{z}_n^{(s)}, \mathbf{w}_n^{(TU,s)}\right) \tag{38}$$

for transcript usage, where $N$ is the number of test cells. The corresponding standard deviations were calculated across the cell-level NLL values and therefore quantify heterogeneity in reconstruction performance among test cells, rather than uncertainty in the across-seed mean.

The gene expression NLL varied moderately across seeds (Supplementary Fig. 20a,b; Supplementary Table 5). Seed 7 showed the highest mean GE NLL and a substantially greater standard deviation than the other models. The mean and dispersion of the transcript usage NLL also varied across seeds (Supplementary Fig. 20c,d; Supplementary Table 5), with seed 5 showing the largest standard deviation. Performance in the two modalities was not strictly concordant; for example, seed 7 performed poorly for GE reconstruction but comparatively well for TU reconstruction.

Seed 8 provided the strongest balanced performance across the two modalities. It achieved both the lowest mean GE NLL and the lowest GE standard deviation ($1044.347 \pm 429.198$), together with a near-minimal TU NLL ($229.593 \pm 104.584$). Seed 6 was marginally better for TU alone ($229.408 \pm 102.988$), but had a higher GE NLL. We therefore selected seed 8 for representative visualisations and downstream analyses that required a single fitted model. Because run-to-run variability has been reported in benchmarks of single-cell multi-modal integration [21], quantitative conclusions were, wherever applicable, interpreted relative to the distribution across all ten seeds rather than from seed 8 alone.

#### 9.2 Cell-type-associated structure is concentrated in the joint gene expression and transcript usage latent space of TRVI

Using the selected seed-8 model, we examined the five TRVI latent representations: P-GE, P-TU, the shared GE and TU components (S-GE and S-TU), and the joint GE–TU representation. Cell embeddings inferred for the complete *Tabula Muris* dataset were visualised using UMAP and coloured by the reference cell-type annotations (Supplementary Fig. 21).

The P-GE and P-TU projections formed densely occupied regions with limited separation by annotated cell type (Supplementary Fig. 21a,b). By contrast, the S-GE and S-TU projections contained distinct, well-separated clusters that broadly aligned with the cell-type annotations (Supplementary Fig. 21c,d). The joint GE–TU representation, inferred from the shared modality-specific components using the learnt cell-specific modality-relevance weights, also separated annotated cell types and placed several related cell types near one another (Supplementary Fig. 21e). For example, B-cell populations, including immature, pre-, naive and late-pro B cells, occupied neighbouring regions, as did endothelial populations such as EC, coronary EC and aortic EC.

These patterns are consistent with the intended decomposition in TRVI: the private representations capture modality-specific variation that is not required for multimodal alignment, whereas the shared components carry cell-type-associated information used to construct the integrated GE–TU representation. The limited cell-type separation in P-GE and P-TU should therefore not be interpreted as evidence that these representations contain no biological signal. Likewise, because UMAP is a nonlinear visualisation, these projections alone do not establish statistical dependence between gene expression regulation and alternative splicing. They instead show that TRVI can combine complementary GE and TU information into an joint representation with reasonable cell-type structure.

##### 9.3 The geometry of the joint gene expression–transcript usage latent representation is robust to random initialisation

Differences in reconstruction NLL across model initialisations do not necessarily imply changes in latent-space geometry. Because TRVI is used to learn joint cell representations for downstream analyses, we therefore tested whether the relative positions of cell types in the GE–TU latent space were preserved across seeds. We quantified global agreement in pairwise cell-type distances, constructed a consensus distance matrix, measured the variability of individual distances and evaluated the reproducibility of local nearest-neighbour relationships.

For seed $s$, let $\mathcal{I}_c$ denote the set of cells annotated as cell type $c$, with $N^{(c)} = |\mathcal{I}_c|$. The centroid of cell type $c$ in the joint GE–TU representation was defined as

$$\bar{\mathbf{z}}_c^{(s)} = \frac{1}{N^{(c)}} \sum_{n \in \mathcal{I}_c} \mathbf{z}_n^{(s)}. \tag{39}$$

Given $C$ cell types, the Euclidean distance matrix $\mathbf{D}^{(s)} \in \mathbb{R}_{\geq 0}^{C\times C}$ for seed $s$ was then computed element-wise for each cell type centroid pair $(i, j)$ as

$$D_{ij}^{(s)} = \left\| \bar{\mathbf{z}}_i^{(s)} - \bar{\mathbf{z}}_j^{(s)} \right\|_2 = \sqrt{\left(\bar{\mathbf{z}}_i^{(s)} - \bar{\mathbf{z}}_j^{(s)}\right)^T \left(\bar{\mathbf{z}}_i^{(s)} - \bar{\mathbf{z}}_j^{(s)}\right)}. \tag{40}$$

To determine whether different seeds recovered the same rank ordering of cell-type relationships, we vectorised the strict upper triangle of each distance matrix, $\mathbf{d}^{(s)} = (D_{ij}^{(s)})_{1\leq i<j\leq C} \in \mathbb{R}_{\geq 0}^{C(C-1)/2}$, and constructed a seed-similarity matrix $\mathbf{R} \in [-1, 1]^{S\times S}$. Each entry was the Spearman rank correlation between the distance vectors for a pair of seeds $s$ and $t$ obtained by

$$R_{s,t} = \rho_{\text{Spearman}}\left(\mathbf{d}^{(s)}, \mathbf{d}^{(t)}\right). \tag{41}$$

For the $S = 10$ TRVI models and $C = 55$ *Tabula Muris* cell types, the distance vectors were highly correlated across all 45 unique seed pairs (Supplementary Fig. 22a). The median Spearman correlation was $\rho_{\text{Spearman}} = 0.958$, with a range of 0.949–0.968. Thus, the global rank ordering of pairwise cell-type distances was strongly preserved across random initialisations.

Next, we assessed the reproducibility of individual, scale-independent cell-type distances. To account for seed-specific differences in the overall scale of the latent space, each seed distance matrix $\mathbf{D}^{(s)}$ was divided by the median of its strict off-diagonal entries

$$\widetilde{D}_{ij}^{(s)} = \frac{D_{ij}^{(s)}}{\text{median}_{p<q}\, D_{pq}^{(s)}}. \tag{42}$$

Then, the consensus distance matrix $\mathbf{D}^{\text{cons}} \in \mathbb{R}_{\geq 0}^{C\times C}$ was defined element-wise for each cell type pair $(i, j)$ as the median normalised distance across seeds for

$$D_{ij}^{\text{cons}} = \text{median}_s\, \widetilde{D}_{ij}^{(s)}. \tag{43}$$

Projecting the $55 \times 55$-dimensional consensus distance matrix as heatmap and ordering the $C = 55$ cell types according to broader lineages (Supplementary Fig. 22b), we observed the consensus distance matrix contained lineage-associated blocks of shorter distances along the diagonal, whereas many between-lineage distances were larger (Supplementary Fig. 22b). This shows that related cell types occupy nearby regions of the GE–TU latent space across seeds.

To identify cell-type relationships that were more sensitive to initialisation, we calculated an interquartile-range matrix $\mathbf{U} \in \mathbb{R}_{\geq 0}^{C\times C}$ from the normalised distances computed element-wise for each cell type pair $(i, j)$ as

$$U_{ij} = Q_{0.75}\left(\left\{\widetilde{D}_{ij}^{(s)}\right\}_{s=0}^{S-1}\right) - Q_{0.25}\left(\left\{\widetilde{D}_{ij}^{(s)}\right\}_{s=0}^{S-1}\right). \tag{44}$$

The interquartile ranges were predominantly low (Supplementary Fig. 22c), indicating that most normalised pairwise relationships varied little across initialisations. This complements the rank-based analysis by showing that the magnitudes of the scale-normalised distances were also generally reproducible.

Finally, we examined the preservation of local cell-type neighbourhoods in the GE–TU latent space. For each seed, we constructed a five-nearest-neighbour graph from the corresponding cell-type distance matrix and represented its edges by a binary adjacency matrix $\mathbf{A}^{(s)}$. The support of an edge between cell types $i$ and $j$ was defined as

$$H_{ij} = \frac{1}{S}\sum_{s=0}^{S-1} A_{ij}^{(s)}, \qquad H_{ij} \in [0,1]. \tag{45}$$

Thus, $H_{ij} = 0.7$ indicates that the neighbourhood edge was recovered in seven of the ten seed-specific graphs. For visualisation, nodes were positioned using the first two principal components of their consensus distance profiles and coloured by broad lineage. Only edges with support $H_{ij} \geq 0.7$ were shown (Supplementary Fig. 22d). The resulting high-support graph formed lineage-enriched neighbourhoods, demonstrating that many local relationships were reproducible across seeds. A small number of cell types, including microglial cells, OC, skeletal muscle satellite cells (SC (muscle)) and chondrocytes, had no edges above the 0.7 threshold. This indicates lower support for their strongest local relationships under the chosen threshold, rather than an absence of neighbours in the individual seed-specific graphs.

Together, the high correlations between seed-specific distance matrices, the low variability of most normalised distances and the recovery of lineage-associated neighbourhoods show that the relative geometry of the joint GE–TU representation is robust to random initialisation. The observed NLL variation therefore did not substantially disrupt the cell-type relationships encoded by TRVI. These results support the use of the joint GE–TU latent space of cell embeddings for downstream analyses, while motivating the continued reporting of seed-level distributions alongside results from the selected seed-8 model.

### Supplementary Note 10: Cell annotation based on marker genes and isoforms determined from TRVI cell embeddings

The joint GE-TU cell embeddings learnt by TRVI consolidate gene expression and alternative splicing-induced transcript usage patterns per cell. This enables us to perform data-driven cell annotation by determining differentially expressed and spliced genes within the same cell embeddings in contrast to two separate ones as done in Supplementary Notes 7 and 8. Here, we demonstrate this joint analysis for cells of brain non-myeloid, heart, bone marrow, and gonadal adipose tissue (GAT) of *Tabula Muris*.

#### 10.1 Brain non-myeloid

Using the trained TRVI model, we inferred joint GE-TU cell embeddings for the single-cell gene expression and alternative splicing-induced transcript usage profiles. After the construction of a nearest-neighbour graph, cell clusters were obtained using the Leiden algorithm and projected on the UMAP of the GE-TU cell embeddings (Supplementary Fig. 23a). The clusters determined recovered the annotation provided by the *Tabula Muris* dataset (Supplementary Fig. 23b) where the Leiden clusters 0, 1, 2, 3, 4, and 5 correspond to OCs, ECs, astrocytes, OPCs, brain pericytes, and another small mixed cluster. We excluded the Leiden cluster 5 from the subsequent analysis since we need to have at least 50 cells per cluster to perform a differential splicing test.

We then performed a differential gene expression and a differential splicing analysis to determine cluster-specific differentially expressed genes (DEGs) and differentially spliced gene (DSG) isoforms. Per cluster, we visualised the normalised mean expression of the top three significant DEGs (Supplementary Fig. 23c) and the mean PSI value (Supplementary Fig. 23d). We observed that the top three DEGs and DSGs per cluster are not the same but cell type-specfic marker genes and cell-type specific isoforms can be determined for each cluster. For example for OCs (Leiden cluster 0), we found the marker genes *Plp1*, *Cnp*, *Ermn* that are accompanied by isoform markers of *Septin4*, *Scarb1*, and *Gsn*. For ECs (Leiden cluster 1), we determined *Cldn5*, *Pltp*, *Bsg* together with the set of isoforms of *Scarb1*, *Septin4*, and *Nrep*.

#### 10.2 Heart

Joint GE-TU cell embeddings of the single-cell gene expression and alternative splicing-induced transcript usage profiles in heart tissue were inferred using the trained TRVI model. We constructed a nearest neighbour graph and obtained clusters using the Leiden algorithm. Projecting the Leiden clusters on the UMAP of GE-TU cell embeddings (Supplementary Fig. 24a) and comparing them with cell annotation provided by *Tabula Muris* (Supplementary Fig. 24b), confirmed that the clusters determined correspond to cell types (0: Fbs (cardiac), 1: ECs (coronary), 2: VICs, 3: Monos, 4: Endocs, 5: mixed). The fifth Leiden cluster is marginally small and consists of some coronary ECs and cardiac Fbs. Since we need at least 50 cells for a differential splicing test, we exclude it from the downstream analysis. heart.

We determined cluster-specific DEGs and DSG isoform on the Leiden clusters and visualised the top three ranked DEGs (Supplementary Fig. 24c) and DSG isoforms (Supplementary Fig. 24d). Comparing the sets of DEGs and DSGs per cluster group, we found they are not the same. However, we discovered cluster-specific marker genes (e.g. *Gsn*, *Dcn*, *Fbln1* for cardiac Fbs; *Fabp4*, *Cd36*, *Cav1* for coronary ECs) and cluster specific DSG isoforms (e.g. *Fmo1*, *Fbln2*, *Lims2* for cardiac Fbs; *Fbln2*, *Art3*, *Lims2* for coronary ECs).

#### 10.3 Bone marrow

For gene expression and alternative splicing-induced transcript usage profiles of cells belonging to bone marrow tissue, we inferred the GE-TU cell embeddings using the TRVI model trained. Using a nearest-neihbour graph and the Leiden algorithm, we determined clusters (Supplementary Fig. 25a) that largely corresponded to cell type annotation provided by *Tabula Muris* (Supplementary Fig. 25b). Distinct cell type-specific clusters were determined by the Leiden clusters 0 (HSC), 4 (Mono (pro)), 5 (macrophage), and 6 (NK cell). Clusters corresponding to larger cell type groups comprised of different developmental stages include cluster 1,2, and 3. Cluster 1 includes granulocytes and granulocytopoietic cells. The latter summarises precursor stages of granulocytes. Both occupy distinct regions in cluster 1. Cluster 2 and 3 both consist of different B cell lineages. Cluster 2 exhibits distinct sub populations of mostly late pro B cells and pre B cells whereas cluster 3 includes immature and naïve B cells. As expected, these developmental stages cluster closely together but TRVI still assigns distinct intra-cluster regions to the respective developmental stage. We also observed a marginally small cluster of naïve B cells and HSCs which we excluded from downstream analysis since its number of cells were too small for a differential splicing test.

The cluster-specific DEGs and DSG isoforms were yielded by differential gene expression and splicing analysis. We visualised the normalised mean expression for the top three significant DEGs in each cluster (Supplementary Fig. 25c) and the mean PSI value for the top three significant DSG isoforms (Supplementary Fig. 25d). We observed that each cluster group can be identified through a set of DEGs and DSG isoforms although the DSG are not necessarily the DEGs. For cluster 5 (macrophages) only two isoforms could be used for the differential splicing test due the sparsity of splicing profiles.

#### 10.4 Gonadal adipose tissue

The GE-TU cell embeddings of gene expression and alternative splicing-induced transcript usage profiles of cells in gonadal adipose tissue (GAT) were inferred by the TRVI model trained. Through nearest-neighbour graph construction and the Leiden algorithm, we determined clusters and projected them on the UMAP of GE-TU cell embeddings (Supplementary Fig. 26a). We found that the clusters determined correspond to the cell type clusters annotated in the *Tabula Muris* dataset (0: EC, 1: myeloid cells, 2: MSCs (adipose), 3: myeloid cells; Supplementary Fig. 26b). However, we excluded the cluster 1 of myeloid cells since it less than 50 cells insufficient for a differential splicing test.

To annotate the cells, we performed differential gene expression and splicing analysis and visualised the top three significant DEGs (Supplementary Fig. 26c) and DSGs isoforms (Supplementary Fig. 26d) per cluster using the normalised mean expression and the mean PSI value. We observe that the sets of DEG and DSG isoforms are not the same per cluster group. However, both sets are discriminative of cell type. For example, myeloid cells (cluster 3) are identified through the marker genes *Lyz2*, *Lyz1*, and *Ccl6* but also through cell-type specific isoforms of *Gsn*, *Cd36*, and *Igf1*.
